# Interpretable multimodal deep learning for real-time pan-tissue pan-disease pathology search on social media

**DOI:** 10.1101/396663

**Authors:** Andrew J. Schaumberg, Wendy C. Juarez-Nicanor, Sarah J. Choudhury, Laura G. Pastrián, Bobbi S. Pritt, Mario Prieto Pozuelo, Ricardo Sotillo Sánchez, Khanh Ho, Nusrat Zahra, Betul Duygu Sener, Stephen Yip, Bin Xu, Srinivas Rao Annavarapu, Aurélien Morini, Karra A. Jones, Kathia Rosado-Orozco, Sanjay Mukhopadhyay, Carlos Miguel, Hongyu Yang, Yale Rosen, Rola H. Ali, Olaleke O. Folaranmi, Jerad M. Gardner, Corina Rusu, Celina Stayerman, John Gross, Dauda E. Suleiman, S. Joseph Sirintrapun, Mariam Aly, Thomas J. Fuchs

## Abstract

Pathologists are responsible for rapidly providing a diagnosis on critical health issues. Challenging cases benefit from additional opinions of pathologist colleagues. In addition to on-site colleagues, there is an active worldwide community of pathologists on social media for complementary opinions. Such access to pathologists worldwide has the capacity to improve diagnostic accuracy and generate broader consensus on next steps in patient care. From Twitter we curate 13,626 images from 6,351 tweets from 25 pathologists from 13 countries. We supplement the Twitter data with 113,161 images from 1,074,484 PubMed articles. We develop machine learning and deep learning models to (i) accurately identify histopathology stains, (ii) discriminate between tissues, and (iii) differentiate disease states. Area Under Receiver Operating Characteristic is 0.805-0.996 for these tasks. We repurpose the disease classifier to search for similar disease states given an image and clinical covariates. We report precision@k=1 = 0.7618±0.0018 (chance 0.397±0.004, mean±stdev). The classifiers find texture and tissue are important clinico-visual features of disease. Deep features trained only on natural images (e.g. cats and dogs) substantially improved search performance, while pathology-specific deep features and cell nuclei features further improved search to a lesser extent. We implement a social media bot (@pathobot on Twitter) to use the trained classifiers to aid pathologists in obtaining real-time feedback on challenging cases. If a social media post containing pathology text and images mentions the bot, the bot generates quantitative predictions of disease state (normal/artifact/infection/injury/nontumor, pre-neoplastic/benign/ low-grade-malignant-potential, or malignant) and lists similar cases across social media and PubMed. Our project has become a globally distributed expert system that facilitates pathological diagnosis and brings expertise to underserved regions or hospitals with less expertise in a particular disease. This is the first pan-tissue pan-disease (i.e. from infection to malignancy) method for prediction and search on social media, and the first pathology study prospectively tested in public on social media. We will share data through pathobotology.org. We expect our project to cultivate a more connected world of physicians and improve patient care worldwide.

## 1 Introduction

The United Nations’ Sustainable Development Goal 3: Good Health and Well-Being suggests that “ensuring healthy lives and promoting the well-being at all ages is essential”, and “increased access to physicians” should be a focus [1]. We therefore take connecting pathologists worldwide to be important. Indeed, Nix *et al.* [2] find pathologists in developing countries (e.g. India, Brazil, and Pakistan) frequently use social media, and 220/1014 (22%) of the posts they analyzed involved “asking for opinions on diagnosis”. The use of social media by pathologists occurs worldwide for both challenging cases and education [3–5]. This suggests social media can facilitate global collaborations among pathologists for novel discoveries [6]. We expand on these approaches by combining (i) real-time machine learning with (ii) expert pathologist opinions via social media to facilitate (i) search for similar cases and (ii) pathological diagnosis by sharing expertise on a particular disease, often with underserved hospitals.

For machine learning to work in general practice, it must be trained on data (i) of sufficient diversity to represent the true variability of what is observed (ii) in a sufficiently realistic setting that may differ from tightly controlled experimental conditions [7]. We therefore (i) collaborate with pathologists worldwide where we use for training the images that these pathologists share to obtain opinions, which are often histopathology microscopy pictures from a smartphone. We did not observe many images from whole slide scanners, which at a global scale have been adopted slowly, due in part to cost and complexities of digital pathology workflows [8,9].

For machine learning to work accurately, it must be trained on a sufficiently large dataset. Our first aim is therefore to curate a large dataset of pathology images for training a machine learning classifier. This is important because in other machine learning domains, e.g. natural vision tasks, datasets of millions of images are often used to train and benchmark, e.g. ImageNet [10] or CIFAR-10 [11]. Transfer learning allows limited repurposing of these classifiers for other domains, e.g. pathology [12–15]. Indeed, we [16] are among many who start in computational pathology [17] with deep-neural networks pre-trained on ImageNet [18–20], and we do so here.

However, computational pathology datasets annotated for supervised learning are often much smaller than millions of images. For example, there are only 32 cases in the training data for a Medical Image Computing and Computer Assisted Intervention challenge (available at http://miccai.cloudapp.net/competitions/82) for distinguishing brain cancer subtypes, and this includes both pathology and radiology images. Other studies are larger, such as the TUmor Proliferation Assessment Challenge (TUPAC16) dataset of 821 cases [21] – all 821 cases being whole slide images from The Cancer Genome Atlas (TCGA) (http://cancergenome.nih.gov/). TCGA has tens of thousands of whole slide images available in total, but these images are only hematoxylin and eosin (H&E) stained slides, and do not represent non-neoplastic lesions such as infections, which are clinically important to correctly diagnose [22]. The main limitation is that obtaining annotations from a pathologist is difficult due to outstanding clinical service obligations, which prevented our earlier efforts from scaling up [23]. We overcome this limitation by curating a large and diverse dataset of 13,626 images from Twitter and 113,161 images from PubMed, where text annotations came from social media post text, hashtags, article titles, abstracts, and/or figure captions.

Equipped with our large dataset, we then address our second main aim, which is to utilize machine learning trained on this dataset to facilitate prospective disease state predictions and search from pathologists in real-time on social media. To that end, we capitalize on a common and systematic approach to diagnosis in which a disease is in one of three classes [22]. Specifically, we use machine learning on pathology images from social media and PubMed to classify images into one of three disease states: *nontumor* (e.g. normal, artifact (Fig S1), injury, infection, or nontumor), *low grade* (e.g. pre-neoplastic, benign, or low grade malignant potential), or *malignant*.

We then implement a social media bot that in real time applies our machine learning classifiers in response to pathologists on social media to (i) search for similar cases, (ii) provide quantitative predictions of disease states, and (iii) encourage discussion (Fig 1). When this bot links to a similar case, the pathologist who shared that case is notified. The ensuing discussions among pathologists are more informative and context-specific than a computational prediction. For instance, to make a diagnosis of Kaposi’s sarcoma, first-world countries have access to an HHV8 histopathology stain, but a pathologist in a developing country may instead be advised to check patient history of HIV because the HHV8 stain is prohibitively expensive. Obviously, a computational prediction of cancer/non-cancer is far less helpful than what humans do: discuss.

**Fig 1:**
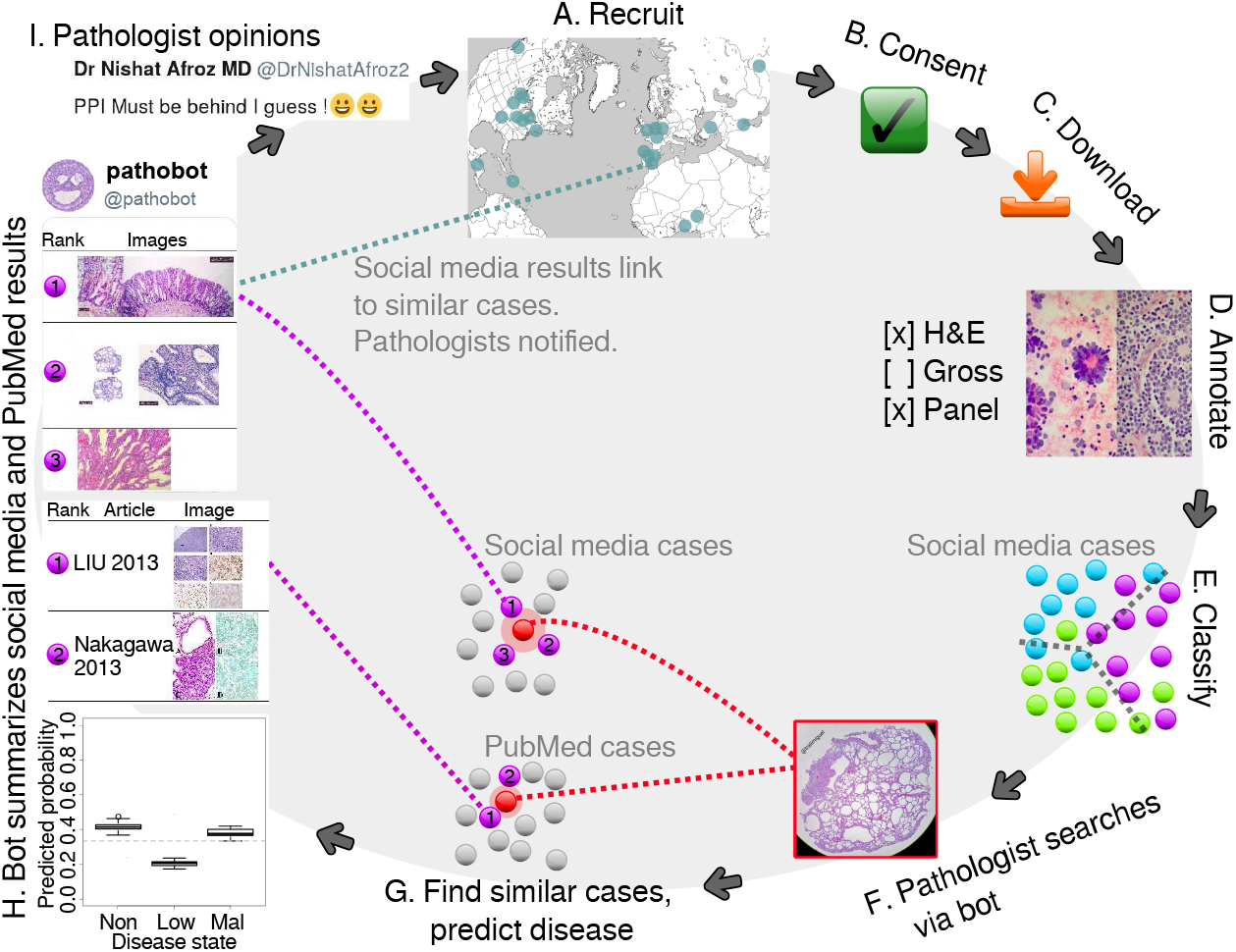
Graphical summary. Pathologists are recruited worldwide (**A**). If a pathologist consents to having their images used (**B**), we download those images (**C**) and manually annotate them (**D**). Next, we train a Random Forest classifier to predict image characteristics, e.g. disease state (**E**). This classifier is used to predict disease and search. If a pathologist posts a case to social media and mentions @pathobot (**F**), our bot will use the post’s text and images to find similar cases on social media and PubMed (**G**). The bot then posts summaries and notifies pathologists with similar cases (**H**). Pathologists discuss the results (I), and some also decide to share their cases with us, initiating the cycle again (**A**). *Procedure overview* in the supplement explains further (Sec S5.4).

In order for machine learning approaches to be useful in a clinical setting, it is critical that these approaches be interpretable and undergo rigorous prospective testing [24]. Furthermore, these approaches need to be accompanied by quantified measures of prediction uncertainty [25]. It may be argued whenever human life is at risk – (i) interpretability, (ii) uncertainty quantification, and (iii) prospective testing are essential – whether the context is medicine or self-driving cars [26,27]. Our social media bot and methods are the first in computational pathology to meet all of these criteria in that (i) we provide multiple levels of interpretability (e.g. Random Forest feature importance and deep learning activation heatmaps), (ii) we statistically quantify prediction uncertainty using ensemble methods, and (iii) we prospectively test in full public view on social media. Concretely, this means (i) a pathologist can interpret what concepts the machine learning finds to be diagnostic in general or what parts of a particular image suggest a specific disease state, (ii) statistical significance, confidence intervals, or boxplots of computational predictions are presented to a pathologist for assessment (e.g. the boxplot in Fig 1 *lower left*), and (iii) in real time a pathologist can interact with our social media bot and method to appraise performance on a case-by-case basis, as well as evaluate the public history of pathologist-bot interactions on social media.

## 2 Materials and methods

### 2.1 Social media data

From Twitter we curate 13,626 images from 6,351 tweets from 25 pathologists from 13 countries. We chose Twitter primarily for its brevity, i.e. one Tweet is at most 280 characters, so we did not expect to need complicated text processing logic to parse tissues or diagnoses. Written permission to download and use the data was obtained from each collaborating pathologist. One pathologist publicly declared their data free to use, so we use these data with acknowledgement. One pathologist donated his glass slide library to another pathologist, and the receiving pathologist shared some received cases on social media, which we treat as belonging to the receiving pathologist. Images are annotated with their tweet text and replies. We use these data for supervised learning.

### 2.2 PubMed data

To represent PubMed data, we download the PubMed Central “Open Access Subset” of 1,074,484 articles. We first trained a classifier to distinguish H&E images from all others on social media (Figs 2, S4, S5), then used the classifier to identify PubMed articles that have at least one H&E figure. From the identified 30,585 articles we retain 113,161 H&E images to comprise our PubMed dataset. Images are annotated with figure caption, article abstract, and article title. This expanded dataset may contain disease that is too rare to be represented in social media data.

**Fig 2:**
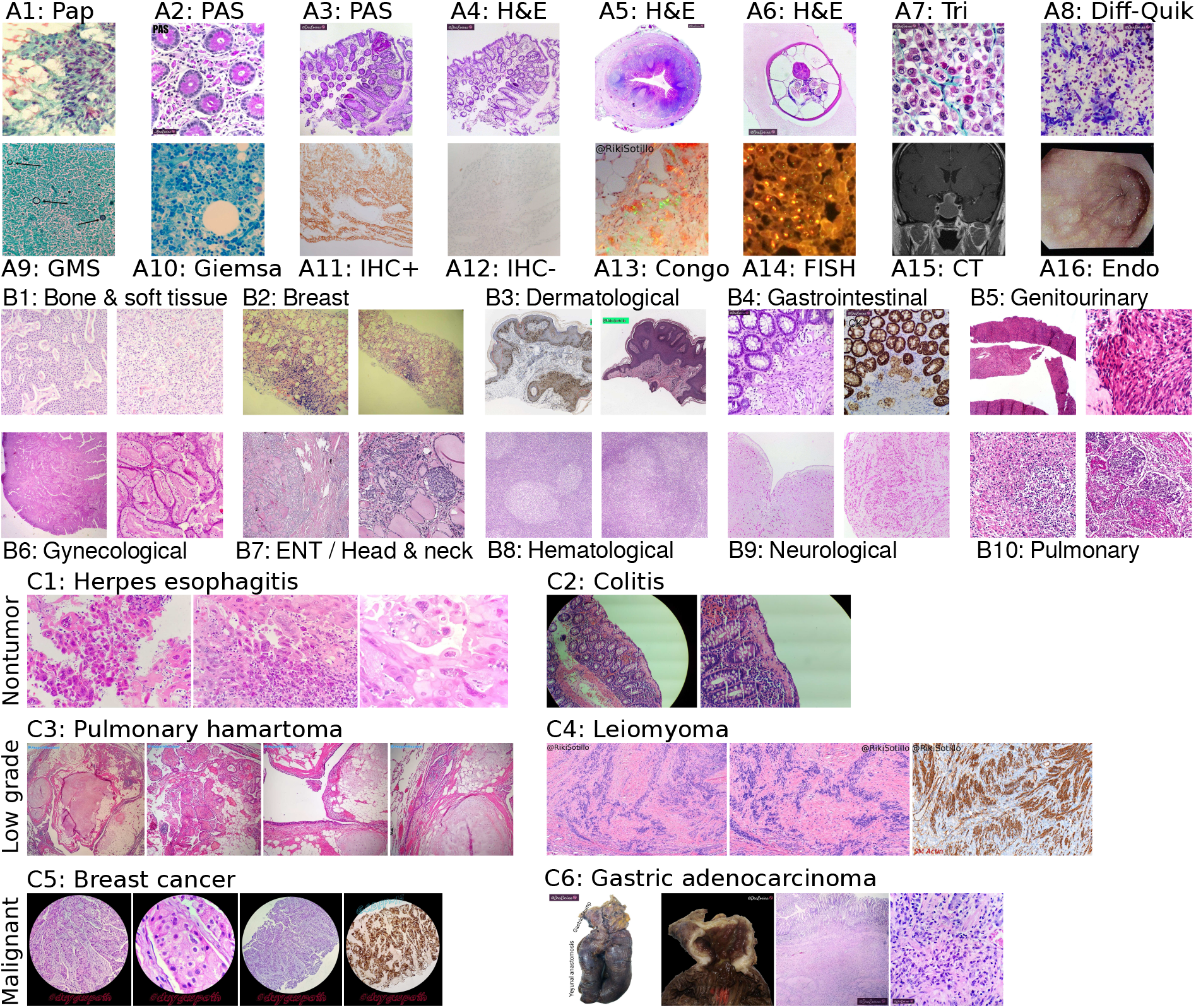
Technique, tissue, and disease diversity. Panel set A shows diverse techniques in our data. Initials indicate author owning image. (**A1**) *R.S.S.:* Papanicolaou stain. (**A2**) *L.G.P*.: Periodic acid-Schiff (PAS) stain, glycogen in pink. (**A3**) *L.G.P*.: PAS stain, lower magnification. (**A4**) *L.G.P.:* H&E stain c.f. Panel A3. (**A5**) *L.G.P.:* H&E stain, human appendix, including parasite *Enterobius vermicularis* (c.f. Fig S2). (**A6**) *L.G.P.:* Higher magnification *E. vermicularis* c.f. Panel A5. (**A7**) *L.G.P.:* Gomori trichrome, collagen in green. (**A8**) *L.G.P.:* Diff-quik stain, for cytology. (**A9**) *R.S.S.:* GMS stain (Intra-stain diversity in supplement details variants, Sec S5.3.1), fungi black. (**A10**) *M.P.P.:* Giemsa stain. (**A11**) *A.M.:* Immunohistochemistry (IHC) stain, positive result. (**A12**) *A.M.:* IHC stain, negative result. (**A13**) *R.S.S.:* Congo red, polarized light, plaques showing green birefringence. (**A14**) *M.P.P.:* Fluorescence *in situ* hybridization (FISH) indicating breast cancer *Her2* heterogeneity. (**A15**) *S.Y.:* Head computed tomography (CT) scan. (**A16**) *L.G.P.:* Esophageal endoscopy. In panel set B we show differing morphologies for all ten histopathological tissue types on Twitter. (**B1**) *C.S.:* bone and soft tissue. We include cardiac here. (**B2**) *K.H.:* breast. (**B3**) *R.S.S.:* dermatological. (**B4**) *L.G.P.:* gastrointestinal. (**B5**) *O.O.F*.: genitourinary. (**B6**) *M.P.P.:* gynecological. (**B7**) *B.X.:* otorhinolaryngological a.k.a. head and neck. We include ocular, oral, and endocrine here. (**B8**) *C.S.:* hematological, e.g. lymph node. (**B9**) *S.Y.:* neurological. (**B10**) *S. M.*: pulmonary. In panel set C we show the three disease states we use: nontumor, low grade, and malignant. (**C1**) *M.P.P.:* Nontumor disease, i.e. herpes esophagitis with Cowdry A inclusions. (**C2**) *K.H.:* Nontumor disease, i.e. collagenous colitis showing thickened irregular subepithelial collagen table with entrapped fibroblasts, vessels, and inflammatory cells. (**C3**) *A.M.:* Low grade, i.e. pulmonary hamartoma showing entrapped clefts lined by respiratory epithelium. (**C4**) *R.S.S.:* Low grade, i.e. leiomyoma showing nuclear palisading. We show IHC completeness but it is not included for machine learning. (**C5**) *B.D.S.:* Malignant, i.e. breast cancer with apocrine differentiation. (**C6**) *L.G.P.:* Malignant, i.e. relapsed gastric adenocarcinoma with diffuse growth throughout the anastomosis and colon. Gross sections (e.g. Fig S3) shown for completeness but not used.

### 2.3 Image processing

We manually curate all social media images, separating pathology from non-pathology images. *Defining an acceptable pathology image* (Sec S5.1.1) details this distinction in the supplement (Fig S4). Some pathologists use our Integrated Pathology Annotator (IPA) tool to browse their data and manually curate the annotations for their cases (Figs S6, S7). We retain non-pathology data publicly posted by consenting pathologists that cannot be publicly distributed to enable building a machine learning classifier that can reliably distinguish pathology from non-pathology images.

### 2.4 Text processing

*Text data overview* (Sec S5.5) in the supplement discusses our text processing to derive ground truth from social media posts (Fig S8). We use hashtags, e.g. #dermpath and #cancer, as labels for supervised learning. We process the text of the tweet and the replies, detecting terms that indicate tissue type or disease state. For instance, “ovarian” typically indicates gynecological pathology, while “carcinoma *in situ*” typically indicates low grade disease (specifically, pre-neoplastic disease in our low grade disease state category). Our text processing algorithm (Fig S8) is the result of author consensus.

### 2.5 Random Forest classifier

We train a Random Forest of 1,000 trees as a baseline for all tasks. A white-balanced image is scaled so its shortest dimension is 512 pixels (px). White balancing helps correct images with reduced blue coloration due to low lighting (Fig S5D). The 512×512px center crop is then extracted, and 2,412 hand-engineered image features are calculated for this crop (Figs 3, S9).

### 2.6 Customized hybrid deep-learning-random-forest model and clinical covariates

#### Image preprocessing and data augmentation

For image preprocessing, a white-balanced image is scaled to be 512 pixels in its shortest dimension, and for deep learning, 224×224px patches are sampled to train a deep convolutional neural network. For deep learning, we use data augmentation of random rotations, random flips, random zoom/rescaling, random brightness variations, Gaussian noise, and Mixup [29]. This means that throughout training hundreds of times over our data we make many small changes to the data each time, e.g. to teach the neural network that rotating an image does not change the diagnosis. *Deep learning* (Sec S5.11.1) discusses further.

#### Deep learning and deep features

To maximize performance by learning disease-state-specific features, we additionally consider deep learning for the most challenging task of disease state prediction. Our deep learning architecture is a ResNet-50 [28] (Fig 3B) pretrained on ImageNet, which we train end-to-end without freezing layers (Fig S13). This means the ResNet-50 deep convolutional neural network is initially trained to classify natural images, e.g. cats and dogs, but every neuron may be adjusted in a data-driven manner for histology-specific learning on our pathology Twitter dataset. To determine how deep feature representations change before and after training the ResNet-50 on histopathology images and covariates, we analyze both (i) ImageNet_2048_ features from the ResNet-50 that has *not* been trained on histopathology data, and (ii) 100 deep features based on the same ResNet-50 where all neurons *have* been further trained on histopathology data. We define ImageNet_2048_ features as the 2,048 outputs from the ResNet-50’s final Global Average Pooling layer, summed over 21 image patches in a grid fashion and concatenated with other features for Random Forest learning (Fig 3C). For histopathology deep learning, we append a 100-neuron fully-connected layer atop the ResNet-50, connecting to the ResNet-50 and covariates, and sum over the same 21 image patches in a grid fashion (Fig 3B). *Deep learning instance and set feature vectors* (Sec S5.8.1) discusses this and the feature interpretability related to the Heaviside step function (Eqns 6, 8).

**Fig 3:**
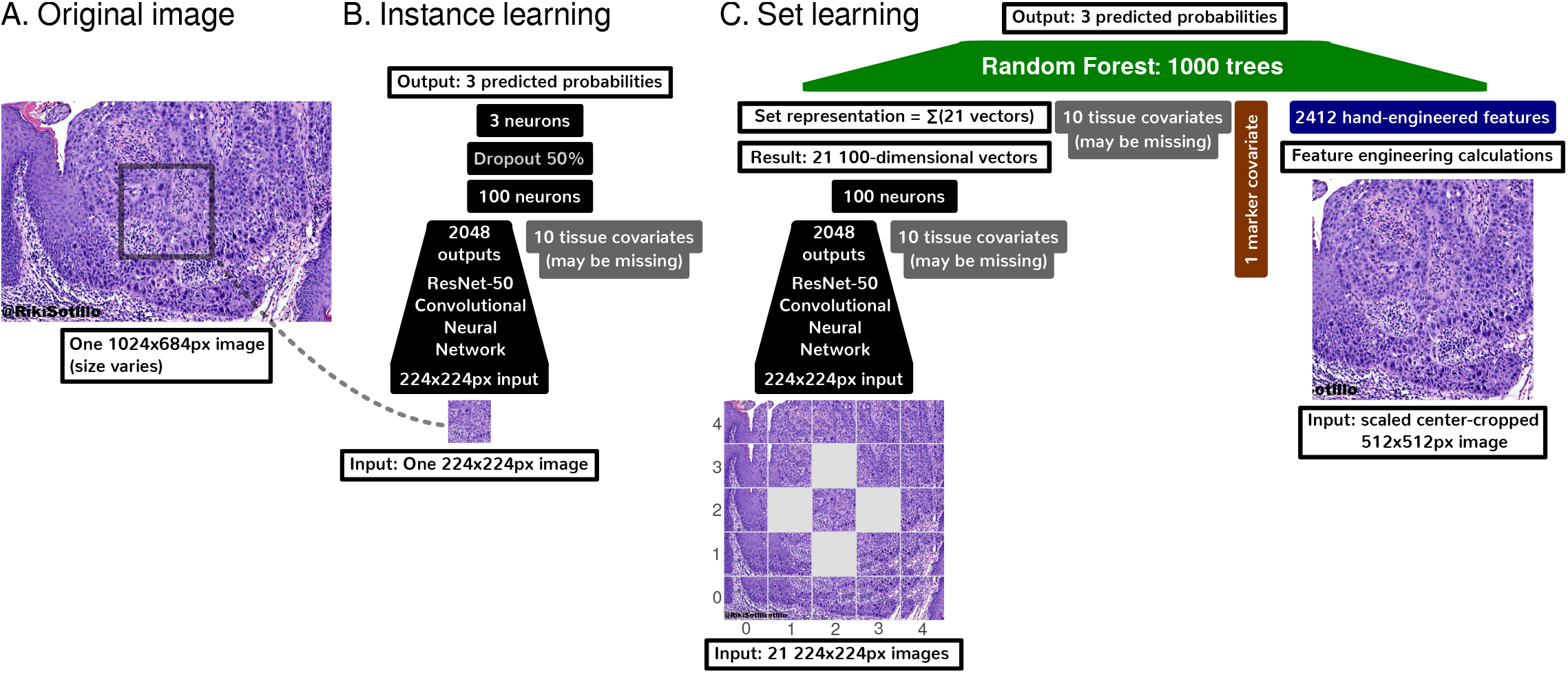
Deep learning methods summary. (**A**) An overall input image may be of any size, but must be at least 512 × 512 pixels (px). (**B**) We use a ResNet-50 [28] deep convolutional neural network to learn to predict disease state (nontumor, low grade, or malignant) on the basis of a small 224×224px patch. This small size is required to fit the ResNet-50 and image batches in limited GPU memory. (**C**) For set learning, this network transforms each of the 21 patches sampled evenly from the image in a grid to a 100-dimensional vector. These 21 patches span the overall input image entirely. For instance, if the overall input image is especially wide, the 21 patches will overlap less in the X dimension. The ResNet-50 converts these 21 patches to 21 vectors. These 21 vectors are summed to represent the overall image, regardless of the original image’s size, which may vary. To represent additional clinico-visual context of a patient case, this sum vector is concatenated with tissue covariates (which may be missing for some images), marker mention covariate, and hand-engineered features. A Random Forest then learns to predict disease state on this concatenation that encodes (i) task-agnostic hand-engineered features (Fig S9) near the image center, (ii) task-specific features from deep learning throughout the image, (iii) whether IHC or other markers were mentioned for this case, and (iv) optionally tissue type. Other machine learning tasks, e.g. histology stain prediction and tissue type prediction, were simpler. For simpler tasks, we used only the Random Forest and 2,412 hand-engineered features, without deep learning.

#### Clinical covariates

To best predict disease state and find similar cases, we seek to include as much patient-related context as possible in our computational pathology machine learning models, so we additionally include clinical information, i.e. tissue type and marker mentions. To represent the tissue type covariate, we include a ten-dimensional one-hot-encoded binary vector to encode which one of the ten possible tissue types is present for this case. If the tissue type is unknown, tissue type is all zeroes for the neural network while being missing values for the Random Forest. We also include a binary one-dimensional marker mention covariate, which is 1 if any pathologist discussing the case mentions a marker test, e.g. “IHC” or “desmin”.

### 2.7 Disease state classifier repurposed for similarity-based search

After we train a *Random Forest classifier* (Sec 2.5) to predict/classify disease state from a variety of deep and non-deep features (Fig 3C), we then use this classifier’s Random Forest similarity metric for search [31,32]. Specifically, our Random Forest consists of 1,000 Random Trees, each of which predicts disease state. If any given Random Tree makes an identical sequence of decisions to classify two histopathology images (each with optionally associated clinical covariates), the similarity of those two images is incremented by one. Aggregating across all Random Trees, the similarity of any two images can therefore be quantified as a number between 0 (not similar according to any Random Tree) and 1,000 (similar according to all 1,000 Random Trees). Equipped with this similarity metric, we repurpose the classifier for search: the classifier takes in a search image and compares it to each other image using this similarity metric, then provides a list of images ranked by similarity to the search image. This approach provides the first pan-tissue (i.e. bone and soft tissue, breast, dermatological, gastrointestinal, genitourinary, gynecological, head and neck, hematological, neurological, pulmonary, etc) pan-disease (i.e. nontumor, low grade, and malignant) patient case search in pathology.

### 2.8 Three levels of sanity checking for search

To inform the physician and to avoid mistakes, sanity checks are important in medicine, or wherever human life may be at risk. Quantifying uncertainty is particularly important [25] in medicine, to assess how much trust to put in predictions that will affect the patient’s care. We are the first to offer three sanity checks for each individual search: (i) prediction uncertainty, (ii) prediction as a check for search, and (iii) prediction heatmaps. *Machine learning sanity checking for search* discusses further (Sec S5.9). Briefly, “prediction uncertainty” relies on an ensemble/collection of classifiers to assess if disease state prediction strength is statistically significant, and if not, the prediction and search using this image should not be trusted. Second, “prediction as a check for search” indicates that if the disease state classification for a given image is assessed as incorrect by a pathologist, search results using this image should not be trusted, because the same classifier is used for both prediction and search.

Third, we use “prediction heatmaps” to show disease-state predictions for each subregion of a given image, based on deep learning. If a pathologist disagrees with these heatmaps, deep-learning-based search for that image cannot be trusted. A failure of any one of these three checks indicates that search results may be incorrect, and they are flagged as such.

### 2.9 Five levels of method interpretability

Interpretability is critical in medicine [24] for physicians to understand whether or not the machine learning is misinterpreting the data. For example, machine learning may uncover that pneumonia patients with a history of asthma have lower mortality risk, suggesting that asthma is protective against pneumonia mortality. However, this would not make sense to a physician, who would instead realize that such patients have lower mortality because they are more likely to be admitted directly to an intensive care unit [33,34]. Asthma is not protective from pneumonia mortality, intensive care is.

Ideally, interpretability facilitates both deductive and inductive human reasoning about the machine learning findings. Deductively, interpretability allows human reasoning about what machine learning finds in specific patient cases, e.g. explaining the malignant prediction overall for a patient by spatially localizing where malignancy-related features are in a histology image. Inductively, interpretability allows human reasoning about broad principles that may be inferred from the machine learning findings overall for a task, e.g. texture features are important in disease state prediction. To the best of our knowledge, it is novel to offer both deductive and inductive interpretability in a pan-tissue pan-disease manner in computational pathology. We do this with (i) hand-engineered feature interpretability (Fig S9), (ii) Random Forest feature importance (Fig 4), (iii) before-and-after-histopathology-training feature importance comparison of deep features to hand-engineered features (Fig 4 vs Fig S10), (iv) deep feature activation maps (Figs 5D, S11), and (v) cluster analyses (Figs 6). *Machine learning interpretability for search* in the supplement discusses further (Sec S5.10).

**Fig 4:**
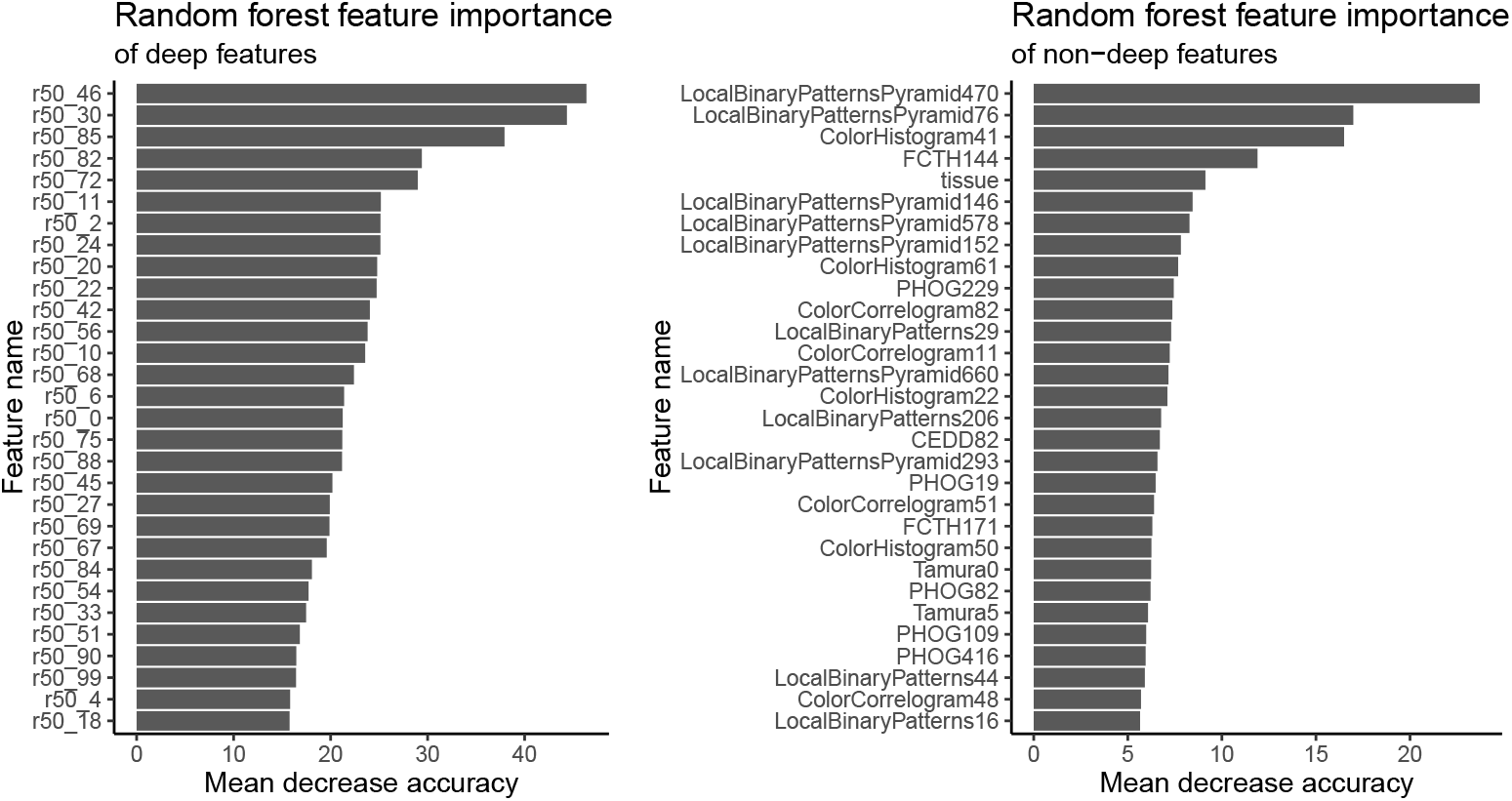
Random Forest feature importance for prioritizing deep features, when non-deep, deep, and clinical features are used together for learning. We use the mean decrease in accuracy to measure Random Forest feature importance. To do this, first, a Random Forest is trained on task-agnostic hand-engineered features (e.g. color histograms), task-specific deep features (i.e. from the ResNet-50), and the tissue type covariate that may be missing for some patients. Second, to measure the importance of a feature, we randomly permute/shuffle the feature’s values, then report the Random Forest’s decrease in accuracy. When shuffling a feature’s values this way, more important features result in a greater decrease in accuracy, because accurate prediction relies on these features more. We show the most important features at the top of these plots, in decreasing order of importance, for deep features (*at left*) and non-deep features (*at right*). The most important deep feature is “r50_46”, which is the output of neuron 47 of 100 (first neuron is 0, last is 99), in the 100-neuron layer we append to the ResNet-50 and train on histopathology images. Thus of all 100 deep features, r50_46 may be prioritized first for interpretation. Of non-deep features, the most important features include Local Binary Patterns Pyramid (LBPP), color histograms, and “tissue” (the tissue type covariate). LBPP and color histograms are visual features, while tissue type is a clinical covariate. LBPP are pyramid-based grayscale texture features that are scale-invariant and color-invariant. LBPP features may be important because we neither control the magnification a pathologist uses for a pathology photo, nor do we control staining protocol. For a before-and-after-training comparison that may suggest the histopathology-trained deep features represent edges, colors, and tissue type rather than texture, we also analyze feature importance of only-natural-image-trained ImageNet_2048_ deep features in conjunction with hand-engineered features (Fig S10). *Marker mention and SIFT features excluded from, Random Forest* feature importance analysis discusses other details in the supplement (Sec S5.10.2).

**Fig 5:**
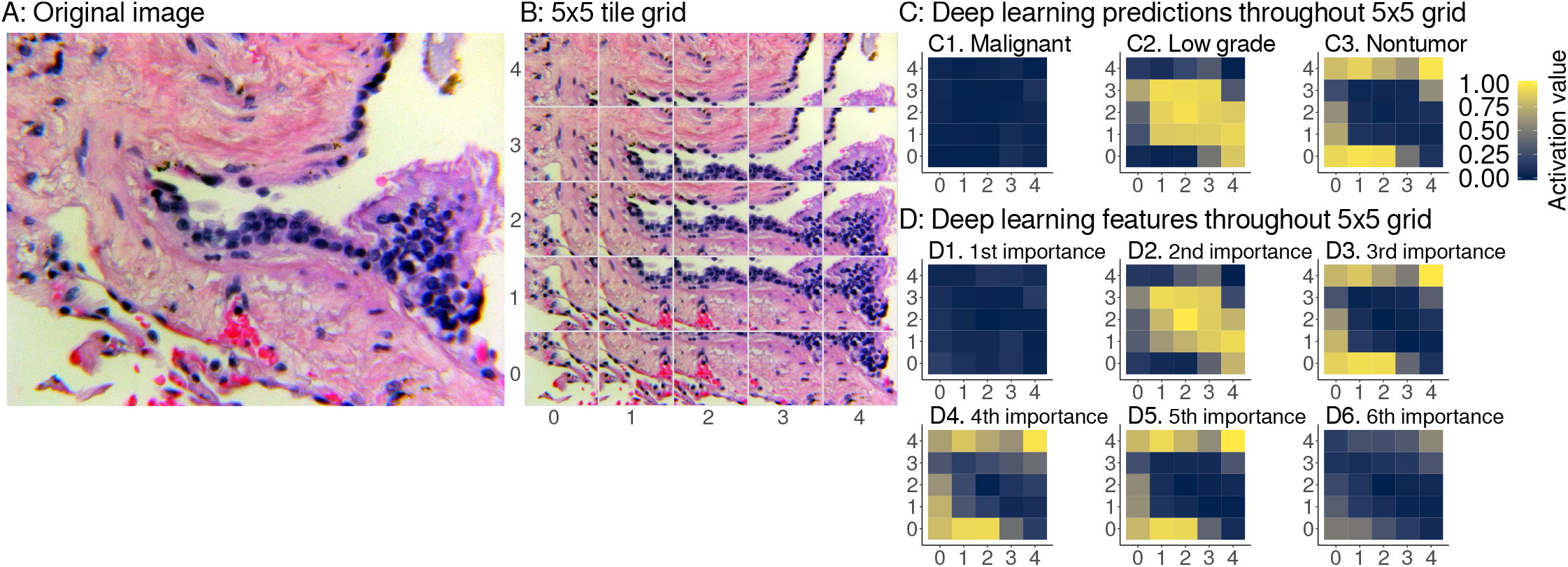
Interpretable spatial distribution of deep learning predictions and features. (**A**) An example image for deep learning prediction interpretation, specifically a pulmonary vein lined by enlarged hyperplastic cells, which we consider to be low grade disease state. Case provided by *Y.R.* (**B**) The image is tiled into a 5×5 grid of overlapping 224×224px image patches. For heatmaps, we use the same 5×5 grid as in Fig 1C *bottom left,* imputing with the median of the four nearest neighbors for 4 of 25 grid tiles. (**C**) We show deep learning predictions for disease state of image patches. (**C1**) throughout the image, predictions have a weak activation value of ~0 for malignant, so these patches are not predicted to be malignant. (**C2**) the centermost patches have a strong activation value of ~1, so these patches are predicted to be low grade. This spatial localization highlights the hyperplastic cells as low grade. (**C3**) the remaining normal tissue and background patches are predicted to be nontumor disease state. Due to our use of softmax, we note that the sum of malignant, low grade, and nontumor prediction activation values for a patch equals 1, like probabilities sum to 1, but our predictions are not Gaussian-distributed probabilities. (D) We apply the same heatmap approach to interpret our ResNet-50 deep features as well. (**D1**) the most important deep feature corresponds to the the majority class prediction, i.e. C1, malignant. (**D2**) the second most important deep feature corresponds to prediction of the second most abundant class, i.e. C2, low grade. (**D3**) the third most important deep feature corresponds to prediction of the third most abundant class, i.e. C3, nontumor. The fourth (**D4**) and fifth (**D5**) most important features also correspond to nontumor. (D6) the sixth most important deep feature does not have a clear correspondence when we interpret the deep learning for this case and other cases (Fig S11), so we stop interpretation here. As expected, we did not find ImageNet_2048_ features to be interpretable from heatmaps, because these are not trained on histpathology (Fig S11A5).

**Fig 6:**
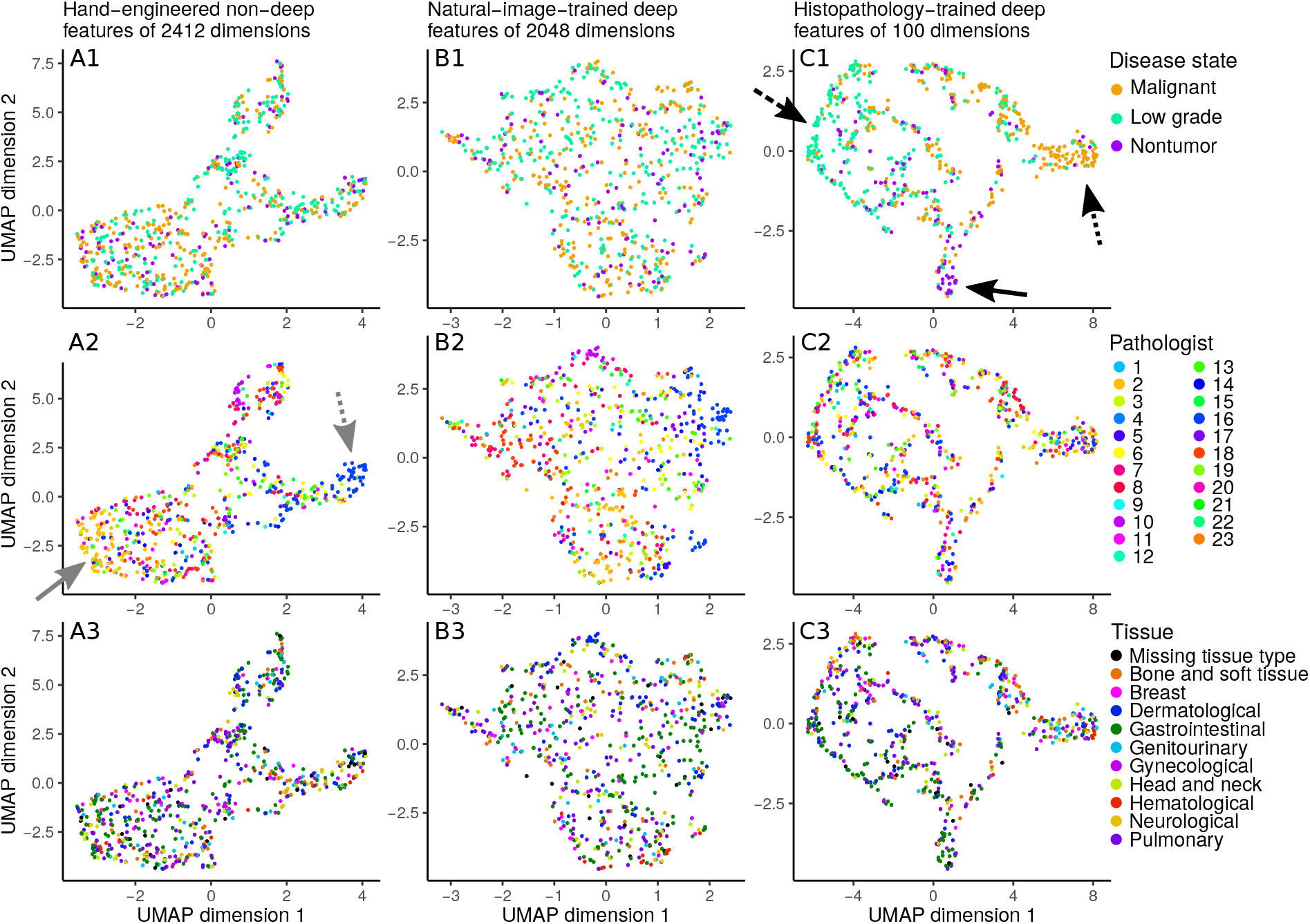
Disease state clusters based on hand-engineered, natural-image-trained deep features, or histopathology-trained deep features. To determine which features meaningfully group patients together, we apply the UMAP [30] clustering algorithm on a held-out set of 10% of our disease state data. Each dot represents an image from a patient case. In general, two dots close together means these two images have similar features. Columns indicate the features used for clustering: hand-engineered features *(at left column*), only-image-trained ImageNet_2048_ deep features *(at middle column*), or histopathology-trained deep features *(at right column*). Rows indicate how dots are colored: by disease state *(at top row*), by contributing pathologist *(at middle row*), or by tissue type *(at bottom row*). For hand-engineered features, regardless of whether patient cases are labeled by disease state (**A1**), pathologist (**A2**), or tissue type (**A3**), there is no strong clustering of like-labeled cases. Similarly, for only-natural-image-trained ImageNet_2048_ deep features, there is no obvious clustering by disease state (**B1**), pathologist (**B2**), or tissue type (**B3**). However, for histopathology-trained deep features, patient cases cluster by disease state (**C1**), with separation of malignant *(at dotted, arrow*), low grade *(at dashed arrow*), and nontumor *(at solid arrow*). There is no clear clustering by pathologist (**C2**) or tissue type (**C3**). The main text notes that hand-engineered features may vaguely group by pathologist (**A2**, pathologists 2 and 16 *at solid and dotted arrows*).

#### Histopathology-trained deep features represent edges, colors, and tissue

To understand what deep features learn to represent after training on histopathology data, we compare Random Forest feature importances of (a) ImageNet_2048_ deep features [not trained on histopathology data] with hand-engineered features and tissue covariate (Fig S10), to (b) 100 deep features [trained on histopathology data] with hand-engineered features and tissue covariate (Fig 4). Before the deep neural network is trained on histopathology data, the tissue covariate as well as edge and color hand-engineered features are important (Fig S10). However, after the deep neural network is trained on histopathology data, tissue is less important while texture hand-engineered features are more important (Fig 4). Therefore, we reason that the deep neural network learns histopathology-relevant edge, color, and tissue features from histopathology data (which reduces the importance of e.g. hand-engineered edge and color features after learning), but the deep neural network may forget histopathology-relevant texture features during learning (which increases the importance of hand-engineered texture features after learning).

#### Interpretability uncovers spatial prediction-to-feature correspondences of disease

Considering both introspective/inductive interpretability (Fig 4) and demonstrative/deductive interpretability (Fig 5), we find a correspondence between important deep features (Fig 4) and the spatial localization of deep learning predictions of disease state (Fig 5). Moreover, we find that using (Eqn 14) the three most important interpretable deep features slightly but significantly improve search performance (Table S1). Deep set learning feature interpretation discusses further (Sec S5.11.2).

#### Deep features trained on histopathology logically cluster patients by disease state, whereas pathology-agnostic features do not

Through cluster analysis we interpret which features (i.e. hand-engineered, only-natural-image-trained, or histopathology-trained), if any, separate patients into meaningful groups, and if the features “make sense” to describe patient histopathology. As expected, neither hand-engineered features (Fig 6A1) nor only-natural-image-trained ImageNet_2048_ deep features (Fig 6B1) cluster patient cases by disease state, presumably because these features are not based on histopathology. These approaches also do not cluster patients by contributing pathologist (Fig 6A2,B2) or by tissue type (Fig 6A3,B3). Additionally, we do not find that reducing dimensionality through principal components analysis qualitatively changes the clustering (Fig S12). In contrast, deep features trained on histopathology data do cluster patients together by disease state (Fig 6C1), but not by pathologist (Fig 6C2) or tissue (Fig 6C3). We conclude that these deep features primarily reflect representations of disease state in a non-tissue-specific manner. It is important to note that any clustering-based result must be carefully scrutinized, because features may suffer from artifacts, e.g. which pathologist shared the patient case. If taken to an extreme, learning to predict disease state on the basis of pathologist-specific staining/lighting/camera artifacts amounts to learning concepts such as, “if pathologist X typically shares images of malignant cases, and a new image appears to be from pathologist X, then this image probably shows malignancy”, which does not “make sense” as a way to predict disease state. Although we did not observe robust clustering by pathologist, even vague grouping by pathologist (Fig 6A2 *at gray arrows*) highlights the importance of critically assessing results. Artifact learning risk is one reason why we (i) rigorously test search through leave-one-pathologist-out cross validation, and (ii) provide sanity checks.

### 2.10 Experimental design and evaluation

We evaluate our classifiers using 10-fold cross validation to estimate bounds of accuracy and Area Under Receiver Operating Characteristic (AUROC) performance metrics. *Supplementary experimental design and evaluation* explains further (Sec S5.14). Because we intend for our methods to accurately find similar cases for any pathologist worldwide, we rigorously test search using leave-one-pathologist-out cross validation and report precision@k. Leave-one-pathologist-out cross validation isolates pathologist cases from one another, so a test set is independent from the corresponding training set. This isolates to a test set pathologist-specific or institution-specific imaging artifacts that may occur from microscopy, lighting, camera, or staining protocol. Thus our leave-one- pathologist-out cross validation measurements quantify our method’s reproducibility, which is critical to measure in medical machine learning [7].

### 2.11 Social media bot for public prospective testing

We present the first pathology-specific social media bot, @pathobot, on Twitter. This bot is a case similarity search tool that applies our methods. Pathologists on Twitter mention the bot in a tweet containing an image. The bot uses our Random Forest classifier to provide disease-state prediction for that image, and search for similar results. Its prediction and search results, along with quantitative assessments of prediction uncertainty, are provided to pathologists in real time. In this way, the bot facilitates prospective tests, and encourages collaboration: as pathologists use the bot, they provide us with complementary qualitative feedback and help us recruit additional collaborators. In this way, the bot facilitates prospective tests, and encourages collaboration: as pathologists publicly use the bot, they provide us with complementary qualitative feedback and these interactions help us recruit additional collaborators.

### 2.12 Computational hardware

For machine learning, we use Weka version 3.8.1 [35] on a laptop. For deep learning, we use Tensorflow Keras [36] on GPUs and a supercomputing cluster. *Supplemetary computational hardware and software* discusses further (Sec S5.15). In R, we perform feature importance analyses with the randomForest package [37] and cluster analyses with the umap package [38].

## 3 Results

### 3.1 Identifying and filtering for H&E images

We ran increasingly difficult tests using increasingly sophisticated machine learning methods. Our first question is the most basic, but arguably the most important: can machine learning distinguish acceptable H&E-stained human pathology images from all others (Figs 2A, S4, S5)? We show acceptable H&E-stained human pathology images can be distinguished from other images – e.g. natural scenes or different histochemistry stains (Fig 7 *at left*) with high performance (AUROC 0.95). Because of the high performance of this classifier, it can be used to partially automate one of our manual data curation tasks, e.g. identifying acceptable images on social media. More importantly, when confronted with over one million PubMed articles, we apply this classifier to filter out all the articles that do not have at least one H&E image. To our knowledge, this is the first H&E image detector to filter PubMed articles. PubMed figures increase our searchable dataset by over an order of magnitude, without any additional manual curation effort. Only with a large dataset may we expect to successfully search for rare diseases, and we currently have 126,787 searchable images. This task also serves as a positive control.

**Fig 7:**
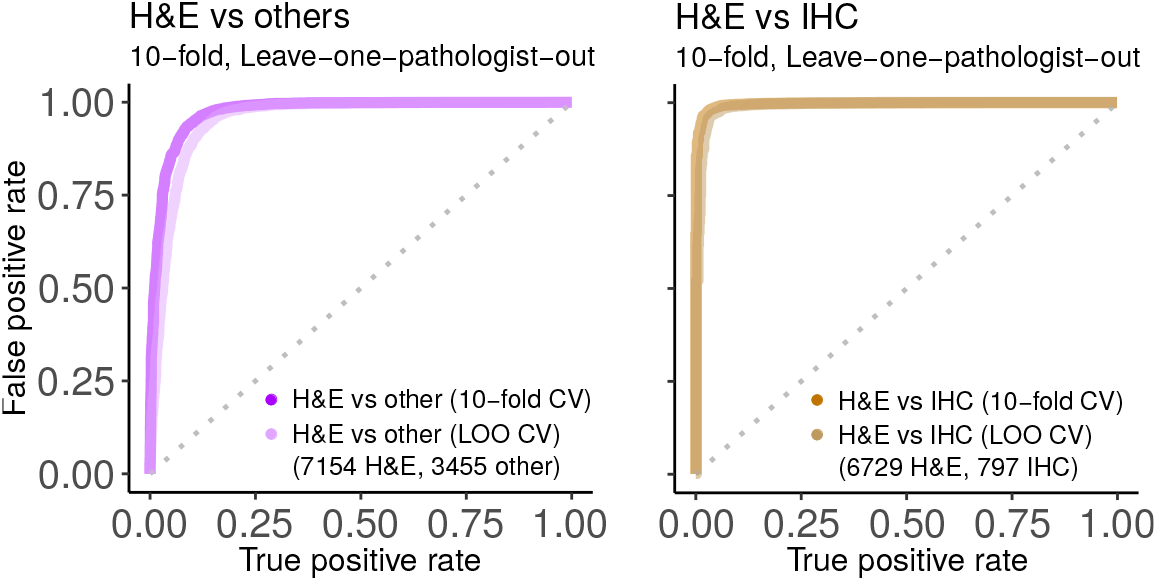
H&E performance. Predicting if an image is acceptable H&E human tissue or not (*at left*), or if image is H&E rather than IHC (*at right*). Ten replicates of ten-fold cross validation (10-fold) and leave-one-pathologist-out cross validation (LOO) had similarly strong performance. This suggests the classifier may generalize well to other datasets. We use the “H&E vs others” classifier to find H&E images in PubMed. Shown replicate AUROC for H&E vs others is 0.9735 for 10-fold (10 replicates of 10-fold has mean±stdev of 0.9746±0.0043) and 0.9549 for LOO (10 reps 0.9547±0.0002), while H&E vs IHC is 0.9967 for 10-fold (10 reps 0.9977±0.0017) and 0.9907 for LOO (10 reps 0.9954±0.0004). For this and other figures, we show the first replicate.

### 3.2 Distinguishing common stain types

H&E and IHC stain types are the most common in our dataset and are common in practice. We therefore ask if machine learning can distinguish between these stain types, which vary in coloration (Fig 2A). Indeed, the classifier performs very well at this discrimination (AUROC 0.99, Fig 7 *at right*). Thus, although IHC coloration can vary between red and brown, machine learning can still successfully differentiate it from H&E. *Intra-stain diversity* explains further (Sec S5.3.1). A well-performing classifier such as this can be useful with large digital slide archives that contain a mixture of H&E and IHC slides that lack explicit labels for staining information. Our classifier can automatically and accurately distinguish these stains, so that downstream pipelines may process each stain type in a distinct manner.

### 3.3 Distinguishing ten histopathology tissue types

We next ask if machine learning can distinguish the ten tissue types present in our Twitter dataset (Fig 2B). *Tissue hashtags and keywords* discusses this further (Sec S5.6.2). The tissue types were distinguishable (AUROC 0.81, Fig 8A) and, as expected, this task was more difficult than stain-related tasks. Being able to identify tissue types may help to detect contaminating tissue in a slide.

**Fig 8:**
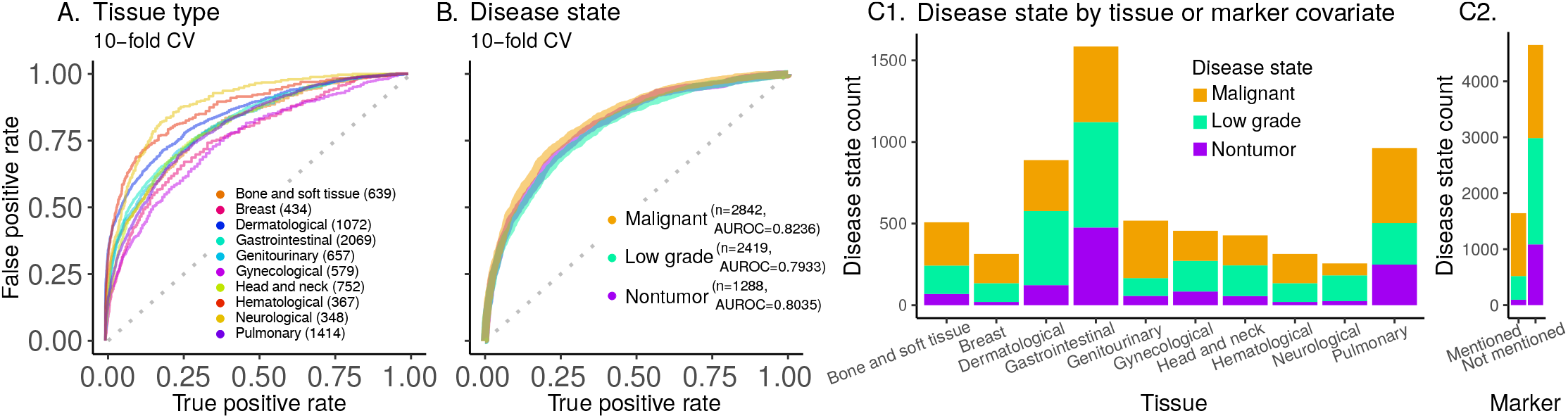
10-tissue type and 3-disease state prediction performance and counts. (**A**) Classifier performance for predicting histopathology tissue type (10 types, 8331 images). (**B**) Classifier performance for predicting disease state (3 disease states; 6549 images). Overall AUROC is the weighted average of AUROC for each class, weighted by the instance count in the class. Each panel (**A** and **B**) shows AUROC (with ten-fold cross-validation) for the chosen classifier. Random Forest AUROC for tissue type prediction is 0.8133 (AUROC for the ten replicates: mean±stdev of 0.8134±0.0007). AUROC is 0.8085 for an ensemble of our deep-learning-random-forest hybrid classifiers for disease state prediction (AUROC for the ten replicates: mean±stdev of 0.8035±0.0043). (**C**) Disease state counts per tissue type. The proportion of nontumor vs. low grade vs. malignant disease states varies as a function of tissue type. For example, dermatological tissue images on social media are most often low grade, but malignancy is most common for genitourinary images. (**D**) Disease state counts as a function of whether a marker test (e.g. IHC, FISH) was mentioned (‘25% of cases) or not. IHC is the most common marker discussed and is typically, but not necessarily, used to subtype malignancies.

### 3.4 Deep learning predicts disease state across many tissue types

Pathologists routinely make decisions about whether a tissue shows evidence of nontumoral disease, low grade disease, or malignant disease, while ignoring spurious artifacts (Fig S1). We therefore ask whether machine learning can perform well on this clinically important task. For this, we use our most common stain type, H&E, including only those images that are single-panel and deemed acceptable (Fig S4). We systematically test increasingly sophisticated machine learning methods (Fig 9) with the goal of achieving the highest possible performance. The simplest baseline model we consider, a Random Forest on the 2,412 hand-engineered features (Fig S9), achieves an AUROC of 0.6843±0.0012 (mean±stdev, Fig 9). Conversely, an ensemble of our deep-learning-random-forest hybrid classifiers achieves much higher performance, with AUROC 0.80 (Fig 9). To our knowledge, this is the first classifier that predicts the full spectrum of disease states, i.e. nontumor, low grade, and malignant (Figs 2, 8B, 9).

**Fig 9:**
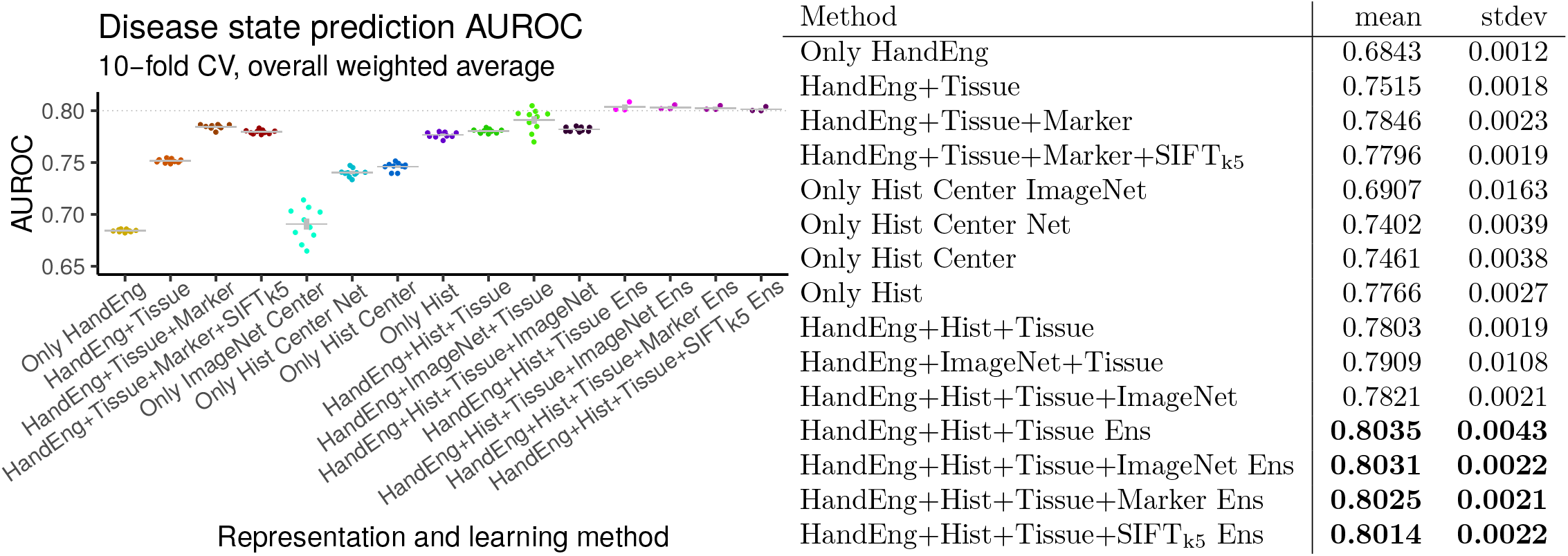
Disease state prediction performance for machine learning methods. For deep learning we use a ResNet-50. For shallow learning we use a Random Forest. We train a Random Forest on deep features (and other features), to combine deep and shallow learning (Fig 3C *top).* Error bars indicate standard error of the mean. Points indicate replicates. Gray lines indicate means. Performance increases markedly when including tissue type covariate for learning (even though tissue type is missing for some patients), when using deep learning to integrate information throughout entire image rather than only the center crop, and when using an ensemble of classifiers. Performance exceeds AUROC of 0.8 (*at right*). We conclude method xii (“HandEng+Hist+Tissue Ens”) is the best we tested for disease state prediction, because no other method performs significantly better and no other simpler method performs similarly. Methods are, from left to right, (i) Random Forest with 2412 hand-engineered features alone for 512×512px scaled and cropped center patch, (ii) Random Forest with tissue covariates, (iii) Random Forest with tissue and marker covariates, (iv) method iii additionally with SIFT_k5_ features for Random Forest, (v) only-natural-image-trained ResNet-50 at same scale as method i with center 224×224px center patch and prediction from a Random Forest trained on 2,048 features from the ResNet-50 (Fig 3) (vi) histopathology-trained ResNet-50 at same scale as method i with center 224× 224px center patch and prediction from top 3 neurons (Fig 3B *top*), (vii) histopathology-trained ResNet-50 with Random Forest trained on 100 features from 224×224px center patch per method vi, (viii) histopathology-trained ResNet-50 features at 21 locations throughout image summed and Random Forest learning on this 100-dimensional set representation with 2,412 hand-engineered features, (ix) method viii with tissue covariates for histopathology-trained ResNet-50 and 2,412 hand-engineered features for Random Forest learning (i.e. Fig 3C sans marker information), (x) method ix with an only-natural-image-trained ResNet-50 instead of a histopathology-trained ResNet-50 for Random Forest learning, (xi) method ix with both an only-natural-image-trained ResNet-50 and a histopathology-trained ResNet-50 for Random Forest learning, (xii) method ix with an ensemble of three Random Forest classifiers such that each classifier considers an independent histopathology-trained ResNet-50 feature vector in addition to 2,412 hand-engineered features and tissue covariate, (xiii) method xii where each Random Forest classifier in ensemble additionally considers only-natural-image-trained ResNet-50 features, (xiv) method xii where each Random Forest classifier in ensemble additionally considers the marker mention covariate (i.e. this is an ensemble of three classifiers where Fig 3C is one of the three classifiers), (xv) method xii where each Random Forest in ensemble additionally considers SIFT_k5_ features for learning.

### 3.5 Texture and tissue are important clinico-visual features of disease

We next determine which features are important to our machine learning classifier for disease state prediction. To do this, we interpret the Random Forest feature importance to gain insight into the clinico-visual features that are predictive of disease state. Our analyses suggest that texture (e.g. Local Binary Patterns) and color (e.g. Color Histograms) features are most important for pathology predictions and search, followed by the tissue type clinical covariate (Fig 4). *Marker mention and SIFT features* excluded, from Random Forest feature importance analysis discusses further (Sec S5.10.2). Our method is therefore multimodal, in that it learns from both visual information in the images and their associated clinical covariates (e.g. tissue type and marker mention). Both modalities improve search performance, as discussed in the following section.

### 3.6 Disease state search, first pan-tissue pan-disease method

In light of pathology-agnostic approaches to pathology search [18,19], we ask if pathology-specific approaches to pathology search may perform better. Indeed, search is the main purpose of our social media bot. Moreover, others have noted task-agnostic features may suffer from poorly understood biases, e.g. features to distinguish major categories (e.g. cats and dogs) in natural images may systematically fail to distinguish major categories in medical images (e.g. ophthalmology or pathology) [39]. To evaluate performance of search, we show precision@k for k =1… 10 (Fig 10). As a positive control, we first test search for similar tissues (Fig 10A), e.g. if the search query image is breast pathology then the top search results should be breast pathology. Here, precision@k=1 = 0.6 means 60% of the time the search query image and top search result image have matching tissue types, e.g. both are breast, or both are gastrointestinal, etc. We subsequently test search for similar disease states (Fig 10B, Table S1), e.g. if the search query image is malignant then the top search results should be malignant. Here, precision@k=1 = 0.76 means 76% of the time the search query image and top search result image have matching disease states (e.g. both malignant, both nontumor, etc), while precision@k=8 = 0.57 means the search query image matches 57% of the top 8 search results, i.e. 4-5 of the top 8 search results are malignant when the search query image is malignant. To estimate performance in general for each method, we perform 10 replicates of leave-one-pathologist-out cross validation with different random seeds (i.e. 0,1,…,9). This allows variance to be estimated for Random Forest learning, but methods based exclusively on the L1 norm are fully deterministic, so these have zero estimated variance (Table S1). We follow two-sample hypothesis testing, where one set of 10 replicates is compared to a different set of 10 replicates. To calculate a U statistic and a p-value, we use the two-tailed Wilcoxon rank-sum test on precision@k=1, which tests for significant differences in precision for the first search result on average. For search’s statistical null model, we train a Random Forest on images with randomly shuffled class labels and evaluate precision@k, as a permutation test (i.e. “RandomForest(2412 + tissue), permutation test” precision@k=1 = 0.3967±0.0044 in Table S1, shown in Fig 10B). We conclude search performs significantly better than chance (0.7618±0.0018 vs 0.3967±0.0044, *U* = 100, *p* = 0.0001817) and offer specifics below.

**Fig 10:**
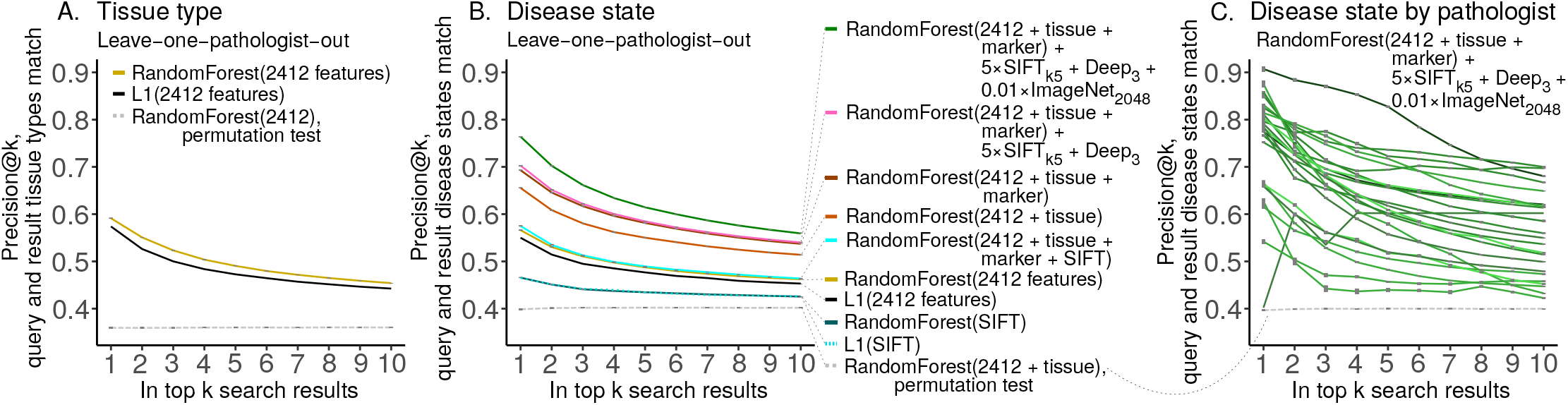
Case similarity search performance. We report search performance as precision@k for leave-one-pathologist-out cross validation for (**A**) tissue and (**B**) disease state. We note search based on SIFT features performs better than chance, but worse than all alternatives we tried. Marker mention information improves search slightly, and we suspect cases that mention markers may be more relevant search results if a query case also mentions markers. SIFT_k5_ and histopathology-trained Deep_3_ features improve performance even less, but only-natural-image-trained ImageNet_2048_ deep features increase performance substantially (Table S1). (C) We show per-pathologist variability in search, with outliers for both strong and weak performance. Random chance performance is shown as a dashed gray line. In our testing, performance for every pathologist is always above chance, which may suggest performance will be above chance for patient cases from other pathologists. We suspect variability in staining protocol, variability in photography, and variability in per-pathologist shared case diagnosis difficulty may underlie this search performance variability. The pathologist where precision@k=1 is lowest shared five images total for the disease prediction task, and these images are of a rare tissue type. Table S2 shows per-pathologist performance statistics.

Results for disease state search are detailed in *supplementary disease state search results* (Sec S5.13). Here, we briefly describe four main findings. First, *clinical covariates improve search performance* (Sec S5.13.1). Both tissue type (0.5640±0.0024 vs 0.6533±0.0025, *U* = 100,*p* = 0.0001796) and marker mention (0.6533±0.0025 vs 6908±0.0021, *U* = 100, *p* = 0.0001796) covariates significantly improve search performance. This suggests that for search these clinical features provide disease state information above and beyond the visual characteristics we have of each image. Second, *in the context of other features, nuclear features of disease are better represented by the most prevalent SIFT clusters rather than all SIFT* (Sec S5.13.2), and the effect of SIFT clusters on search is small but significant (0.6908±0.0021 vs 0.6935±0.0029, U = 19.5, p = 0.02308). This indicates nuclear features, as represented by SIFT, provide limited but complementary disease-related information for search. Third, *deep features synergize with other features, informing search more than nuclear SIFT features, but less than clinical covariates* (Sec S5.13.3). Specifically, deep features improve search performance less than tissue type (0.5720±0.0036 vs 0.6533±0.0025, *U* = 0, *p* = 0.0001806) and less than marker mentions (0.6602±0.0022 vs 0.6908±0.0021, *U* = 0, *p* = 0.0001817, but more than SIFT clusters (0.6983±0.0016 vs 0.6948±0.0032, *U* = 83.5, *p* = 0.01251). Fourth, *deep features trained only on natural images outperform hand-engineered features for search, and offer best performance when combined with other features* (Sec S5.13.4). Particularly, in the context of clinical covariates, ImageNet_2048_ features demonstrate high importance by offering better search performance than the 2,412 hand-engineered features, SIFT_k5_ features, and histopathology-trained Deep_3_ features *combined* (0.7517±0.0025 vs 0.7006±0.0026, *U* = 100, *p* = 0.0001817) – although this may change as more data become available or more advanced methods are used. Moreover, we found that adding only-natural-image-trained ImageNet_2048_ deep features to our best-performing model (incorporating hand-engineered features, tissue type, marker mention, SIFT_k5_ features, and Deep_3_ features) improved search performance further (0.7006±0.0026 vs 0.7618±0.0018, *U* = 0, *p* = 0.0001817), and was the best-performing search method we measured. Taken together, we conclude (i) texture and tissue features are important, (ii) histopathology-trained deep features are less important, (iii) nuclear/SIFT features are least important for disease state search, and (iv) in the context of clinical covariates the only-natural-image-trained ImageNet_2048_ deep features are the most important visual features we tested for search.

## 4 Discussion

### 4.1 Summary

Pathologists worldwide reach to social media for opinions, often sharing rare or unusual cases, but replies may not be immediate, and browsing potentially years of case history to find a similar case can be a time-consuming endeavor. Therefore, we implemented a social media bot that in real-time searches for similar cases, links to these cases, and notifies pathologists who shared the cases, to encourage discussion. To facilitate disease prediction and search, we maintain a large pathology-focused dataset of 126,787 images with associated text, from pathologists and patients the world over. This is the first pan-tissue, pan-disease dataset in pathology, which we will share with the community through pathobotology.org to promote novel insights in computational pathology. After performing stain- and tissue-related baselines with a Random Forest, we performed a number of analyses on this dataset for disease state prediction and search. To accomplish this, we developed a novel synthesis of a deep convolutional neural network for image set representations and a Random Forest learning from these representations (Figs 3, S14). We found this model can classify disease state with high accuracy, and be repurposed for real-time search of similar disease states on social media. This interpretable model, combined with its social media interface, facilitates diagnoses and decisions about next steps in patient care by connecting pathologists all over the world, searching for similar cases, and generating predictions about disease states in shared images. Our approach also allowed us to make a number of important methodological advances and discoveries. For example, we found that both image texture and tissue are important clinico-visual features of disease state – motivating the inclusion of both of feature types in multimodal methods such as ours. In contrast, we find deep features trained only on natural images (e.g. cats and dogs) substantially improve search performance, while pathology-specific deep features and cell nuclei features improve less, although combining all these performed best. Finally, we provide important technical advances, because our novel deep feature regularization and activation functions yield approximately binary features and set representations that may be applicable to other domains. In sum, these advances readily translate to patient care by taking advantage of cutting-edge machine learning approaches, large and diverse datasets, and interactions with pathologists worldwide.

### 4.2 Comparison with prior studies

Our approach builds on, but greatly extends, prior work in the field of computational pathology. We comment on this briefly here, and describe more fully in *supplementary comparison with prior studies* (Sec S5.16). First, much of prior work involves a subset of tissue types or disease states [40–42]. However, our study encompasses diverse examples of each. Second, prior studies investigating pathology search take a variety of pathology-agnostic approaches, e.g. (i) using neural networks that were not trained with pathology data [18,19] or (ii) using scale-invariant feature transform (SIFT) features [19, 43, 44] that do not represent texture or color [45]. Our inclusive approach is different, building a search method for pathology data represented by thousands of features – including SIFT clusters, neural networks, other visual features, and clinical covariates. Our model outperforms pathology-agnostic baselines.

Prior work has found texture and/or color to be important for tissue-related tasks in computational pathology [46–48]. We find texture and color to be important for disease-related tasks. Additionally, we go a step further by comprehensively considering the relative contributions of many clinico-visual features to the prediction and search of disease. Such important features include texture, color, tissue type, marker mentions, deep features, and SIFT clusters.

### 4.3 Caveats and future directions

Below we discuss the primary caveats (also see *supplementary caveats* in Sec S5.17) and future directions (also see *supplementary future directions* in Sec S5.18).

#### Diagnosis disagreement or inaccuracy

First, there is a risk of error in our data because many different pathologists share cases, and they may disagree on the most appropriate hashtags or diagnosis. Moreover, there may be diagnostic inaccuracies from the pathologist who posted the case, or other pathologists. We find these situations to be rare, but if they occur, the case tends to have an increased amount of discussion, so we can identify these situations. Second, our nontumor/low-grade/malignant keyword rules may be incorrect or vague. For these first and second caveats, we take a majority vote approach, manually curate as needed, and discuss. Indeed, as we discussed amongst ourselves the hyperplasia in Fig 5, it became clear we needed to explicitly mention pre-neoplastic disease is included in the low grade disease state category.

#### Dataset case sampling and region of interest biases

Our dataset may have both (i) a case sampling bias and (ii) a region of interest sampling bias. First, there may be case sampling bias if we typically have unusual cases that pathologists consider worth sharing, and our cases by necessity only come from pathologists on social media. We plan to advocate sharing of normal tissue and less unusual cases to circumvent this bias. Second, the pathologist who shares the case chooses which images to share, typically sharing images of regions of interest that best illustrate the diagnosis, while ignoring other slides where the diagnosis is less straightforward. In future work, we will include whole slide images for additional context.

#### Dataset size and granularity

To increase the granularity and accuracy of tissue type predictions, we first plan to expand the size of this dataset by recruiting more pathologists via social media, aiming to have representative images for each organ. There are many organs within the gastrointestinal tissue type, for instance. Additionally, we expect our dataset to broaden, including more social media networks and public pathology resources such as TCGA, with our bot integrating these data for search and predictions.

## Conclusion

We believe this is the first use of social media data for pathology case search and the first pathology study prospectively tested in full public view on social media. Combining machine learning for search with responsive pathologists worldwide on social media, we expect our project to cultivate a more connected world of physicians and improve patient care worldwide. We invite pathologists and data scientists alike to collaborate with us to help this nascent project grow.

## Acknowledgments

A.J.S. thanks Dr. Marcus Lambert and Pedro Cito Silberman for organizing the Weill Cornell High School Science Immersion Program. A.J.S. thanks Terrie Wheeler and the Weill Cornell Medicine Samuel J. Wood Library for providing vital space for A.J.S., W.C., and S.J.C. to work early in this project. A.J.S. thanks Dr. Joanna Cyrta of Institut Curie for H&E-saffron (HES) discussion. A.J.S. thanks Dr. Takehiko Fujisawa of Chiba University for his free pathology photos contributed to social media and this project via @Patholwalker on Twitter.

A. J.S. was supported by NIH/NCI grant F31CA214029 and the Tri-Institutional Training Program in Computational Biology and Medicine (via NIH training grant T32GM083937). This research was funded in part through the NIH/NCI Cancer Center Support Grant P30CA008748.

S.Y. is a consultant and advisory board member for Roche, Bayer, Novartis, Pfizer, and Amgen – receiving an honorarium.

T.J.F. is a founder, equity owner, and Chief Scientific Officer of Paige.AI.

We are grateful to the patients who made this study possible.

## Author Contributions

**Conceptualization**: AJS, MA.

**Data curation**: AJS, WCJ, SJC, LGP, BSP, MPP, NZ, BDS, SY, AM.

**Methodology, Software, Validation, Formal analysis, Investigation, Writing (original draft):** AJS.

**Funding acquisition**: AJS, TJF.

**Project administration**: AJS, WCJ, SJC, MA, TJF.

**Resources (pathology) and discussion:** LGP, BSP, MPP, RSS, KH, NZ, BDS, SY, BX, SRA, AM, KAJ, KRO, SM, CM, HY, YR, RHA, OOF, JMG, CR, CS, JG, DES.

**Resources (computational):** AJS, TJF.

**Supervision:** MA, TJF.

**Visualization, wrote annotation files**: AJS, WCJ, SJC.

**Writing (editing):** AJS, MA.

**Writing (reviewing):** AJS, LGP, BSP, MPP, SY, BX, AM, OOF, JMG, CS, SJS, MA, TJF.

**Answered annotator questions**: LGP, BSP, MPP, NZ, BDS, SY, BX, SRA, AM, SM, CM, HY, YR, RHA, OOF, CS, SJS.

---

**Fig S1:**
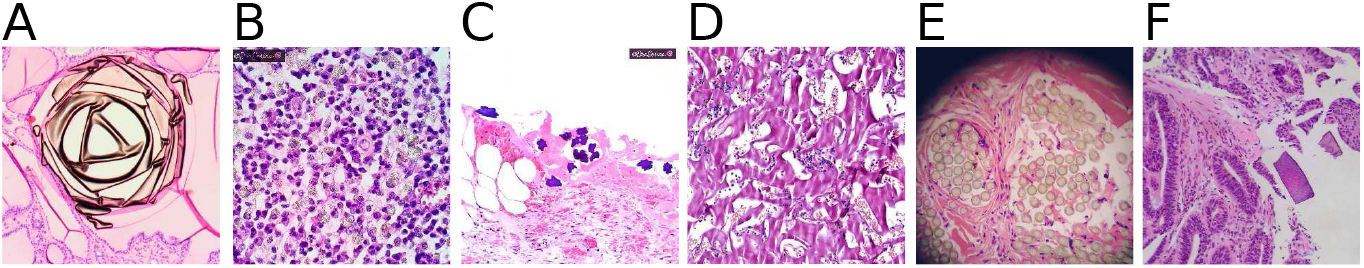
Artifact and foreign body images. Our dataset includes artifacts and foreign bodies which machine learning should not consider prognostic. All panels human H&E. (**A**) *B.X.:* Colloid. (**B**) *L.G.P.:* Barium. (**C**) *L.G.P.:* Oxidized regenerated cellulose, a. k.a. gauze, granuloma may mimic mass lesion [49]. (**D**) *R.S.S.:* Hemostatic gelatin sponge, a.k.a. Spongostan^TM^, may mimic necrosis. (**E**) *S.Y.:* Sutures, may mimic granuloma or adipocytes. (**F**) *L.G.P.:* Crystallized kayexelate, may mimic mass lesion or parasite.

**Fig S2:**
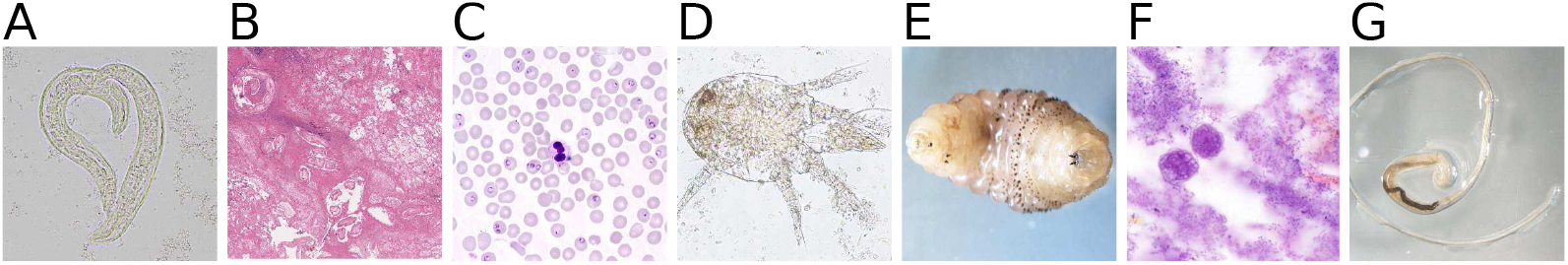
Parasitology images. Our dataset includes diverse parasitology samples. (**A**) *B.S.P.: Strongyloides stercoralis,* light microscopy. (**B**) *B.S.P.: Dirofilaria immitis*, in human, H&E stain. (**C**) *B.S.P.: Plasmodium falciparum,* in human, Giemsa stain. (**D**) B. *S.P.:* Incidental finding of unspecified mite in human stool, light microscopy. (**E**) *B.S.P.: Dermatobia hominis,* live gross specimen. (**F**) *B.S.P.: Acanthamoeba,* in human, H&E of corrective contact lenses. (**G**) *B.S.P.: Trichuris trichiura,* gross specimen.

## S5 Supporting information

### S5.1 Image data overview

The goal of obtaining images from practicing pathologists worldwide is to create a dataset with a diverse and realistic distribution of cases. A worldwide distribution (Fig 1A) may be appropriate to overcome potential biases inherent at any single institution, such as stain chemistries or protocols. Our dataset includes a wide variety of stains and techniques (Fig 2A) – even variety for a single stain, e.g. H&E stains (Fig S5). H&E stain composition may vary by country – e.g. in France, H&E typically includes saffron, which stains collagen fibers. Phyloxin may be used instead of eosin. This helps differentiate between connective tissue and muscle, or to see cell cytoplasm better against a fibrous background. This stain may be referred to as “HES” or “HPS”, and we consider it H&E. *Intra-stain diversity* discusses further (Sec S5.3.1). Our dataset includes gross sections (Fig S3) that pathologists share alongside images of stained slides. In addition to variation in the signal of interest (i.e., stain, tissue, or disease), we find variability in the noise (i.e. pathology artifacts, Fig S1). Such noise may initially seem undesirable, but is likely important for machine learning techniques to robustly predict which image motifs are relatively unimportant rather than prognostic. Finally, our dataset includes a variety of parasites and other [micro]organisms (Fig S2, and Fig S5A,E), an important consideration in developing countries.

#### S5.1.1 Defining an acceptable pathology image

To create our pathology social media database, we first identified pathology images, and second, narrowed down the set of pathology images into those that were of sufficient quality to be used and could be shared publicly. By “pathology image”, we mean images that a pathologist may see in clinical practice, e.g. gross sections, microscopy images, endoscopy images, or X-rays. An image designated as a “pathology image” is not necessarily an image of diseased tissue. After we identified pathology images, we screened them for inclusion in our dataset. “Acceptable images” are those that do not meet rejection or discard criteria defined below. If an acceptable image is personally identifiable or private (see criteria below), we retain the image for some machine learning analyses, but do not distribute the image publicly [for legal reasons].

**Fig S3:**
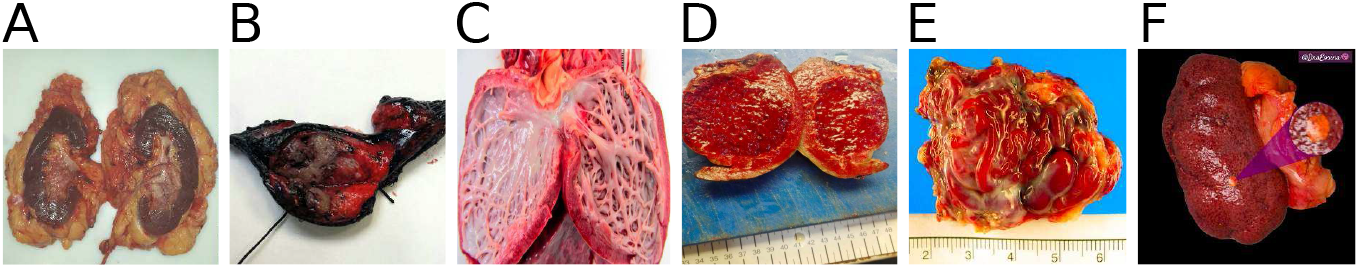
Gross images. Gross sections are represented in our dataset, putting the slide images in context. (**A**) M.P.P: Urothelial carcinoma. (**B**) *M.P.P.*: Lung adenocarcinoma. (**C**) *S.R.A.*: Barth syndrome. (**D**) *N.Z.*: Enlarged spleen. (**E**) *S.R.A.*: Arteriovenous malformation. (**F**) *L.G.P.*: Kidney adrenal heterotopia.

**Fig S4:**
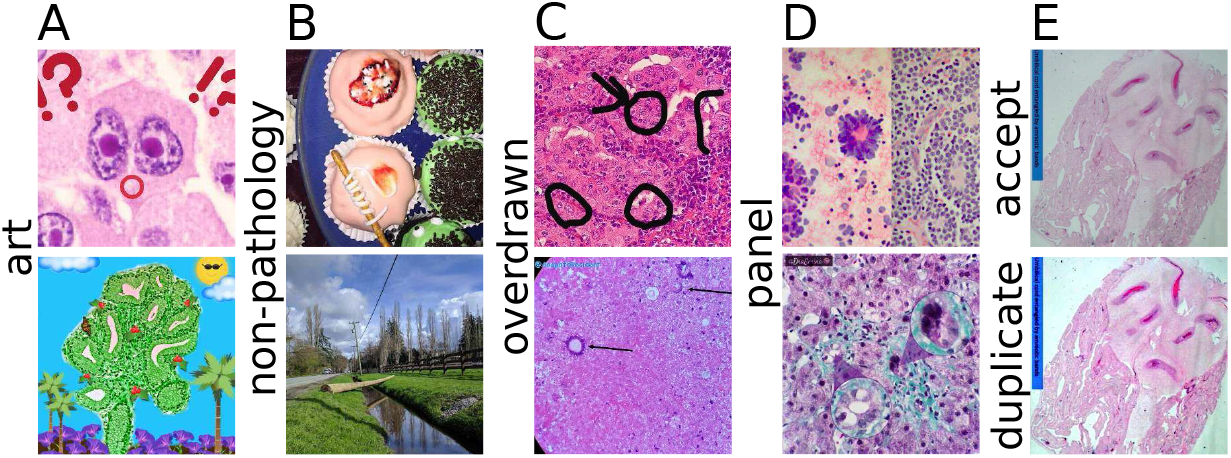
Image acceptability criteria. Examples of images that are rejected, because they are not pathology images that a pathologist would see in clinical practice. (**A**) *top M.P.P., bottom B.D.S*: “art” rejects. (**B**) *top B.S.P.*, bottom S.Y.: “non-pathology” rejects. (**C**) *top B.X., bottom A.M.:* “overdrawn” rejects. (**D**) *top S.R.A., bottom L.G.P.:* “panel” is rejected for some tasks, e.g. H&E vs IHC or disease state prediction, but not for others, e.g. H&E vs others. The H&E vs others task retains multi-panel images because multi-panel images that include an H&E panel should be included in our PubMed search results, and this classifier is used to filter PubMed. (**E**) *top and bottom S.R.A.:* top is acceptable H&E (see Sec S5.1.1 for definition), bottom is “dup” [duplicate] rejection.

##### Criteria for rejected, discarded, private, or acceptable images

For our manual data curation process, we defined several rejection criteria (Fig S4), detailed in Section S5.2.1. Figure S4A shows examples of images rejected as “art”, because they are artistically manipulated H&E pathology microscopy images. Figure S4B shows examples of images rejected as “non-pathology”, e.g. parasitology-inspired cupcakes *(top*) and a natural scene image *(bottom*). Non-pathology images are relatively common on pathologists’ social media accounts, though we try to minimize their frequency by recruiting pathologists who primarily use their accounts for sharing and discussing pathology. Figure S4C shows examples of images rejected as “overdrawn”. Overdrawn images are those that have hand-drawn marks from a pathologist (which pathologists refer to as “annotations”), which prevent us from placing a sufficiently large bounding box around regions of interest while still excluding the hand-drawn marks.

Section S5.2.2 discusses our “overdrawn” criterion in detail. Figure S4D shows examples of images rejected as “panel”, because they consist of small panels (*top*) or have small insets (*bottom*); splitting multi-panel images into their constituent single-panel images would substantially increase our manual curation burden. Figure S4E *top* is an acceptable H&E-stained pathology image. Figure S4E *bottom* is rejected as a duplicate of the S4E *top* image, though the colors have been slightly modified, and the original image is a different size.

**Fig S5:**
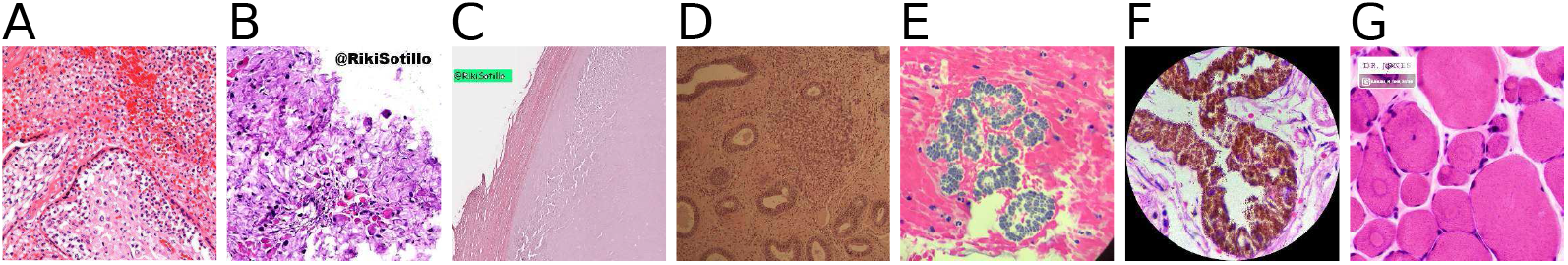
H&E images. Our dataset includes diverse H&E-stained slide microscopy images. (**A**) *S.R.A.:* Acute villitis due to septic *Escherichia coli.* (**B**) *R.S.S.:* Garlic. (**C**) *R.S.S.:* “Accellular” leiomyoma after ulipristal acetate treatment. (**D**) *R.S.S.:* Brownish appearance from dark lighting. (**E**) *K.R.O.: Sarcina* in duodenum. (**F**) *B.D.S.*: Mature teratoma of ovary, pigmented epithelium. (**G**) *K.A.J.:* Central core myopathy.

### S5.2 Supplementary Image processing

#### S5.2.1 Criteria details for rejected, discarded, private, or acceptable images

Though criteria are outlined in *criteria for rejected, discarded, or acceptable images* (Sec S5.1.1) – more formally, we reject the following image types, during our manual data curation process:

1. Non-pathology images, such as pictures of vacations or food.
2. Multi-panel images, such as a set of 4 images in a 2×2 grid. Images with insets are also rejected. For “H&E vs IHC”, “tissue type”, and “disease state” tasks, we only accept single-panel images, and leave for future work the complexities of splitting multi-panel images into sets of single-panel images. We accept multi-panel images for the “H&E vs others” task, because we use the classifier trained for this task to filter PubMed, and many H&E images in PubMed are multi-panel, which are useful as search results. Multi-panel images may have black dividers, white dividers, no dividers, square insets in a corner, or floating circular insets somewhere in the image. There may be two or more panels/insets. Per-pixel labels for each panel may be the best solution here, and would support a machine learning approach to split multi-panel images to reduce this additional manual data curation burden.
3. Overdrawn images, where a 256×256px region could not bound all regions of interest in an image. This occurs most frequently if a pathologist draws by hand a tight circle around a region of interest, preventing image analysis on the region of interest in a way that completely avoids the hand-drawn marks.
4. Images that manipulate pathology slides into artistic motifs, such as smiley faces or trees. In contrast, a picture of a painting would be a non-pathology image.

Moreover, we completely discard from analysis certain types of images:

1. Duplicate images, according to identical SHA1 checksums or by a preponderance of similar pixels. However, duplicate images may be shown in search results, if the images are contained in different tweets, because there may be different replies to these tweets as well.
2. Corrupt images, which either could not be completely downloaded or employed unusual JPEG compression schemes that Java’s ImageIO^1^ library could not open for reading.
3. Pathology images that are owned by pathologists who have not given us explicit written permission to use their images. Consider the following example. When a pathlogist gives us permission to download data, our software bot downloads thousands of that pathologists’s social media posts regardless if some of the images in those posts are actually owned by a different pathologist who did not give us permission. We detect these cases when we manually curate the pathologist’s data, and discard these images belonging to pathologists who have not given us permission. To elaborate, pathology images that are taken by pathologists and shared on social media are treated the same way as pathology images taken from case reports or copyrighted manuscripts, i.e. if the pathologist or publisher has not provided us explicit written permission to use the image, we discard the pathology image and do not use it.

Images that are not rejected or discarded are deemed “acceptable” pathology images. However, for legal reasons, we cannot distribute all of the images we have from social media, namely:

1. Pathology images obtained from children (including fetuses), which may be identifiable. The data shared on social media are anonymized; thus, we do not have contact information for the child’s parent and therefore cannot obtain consent to distribute a picture of e.g. a child’s X-rays or gross specimens. Although unlikely to be identified by the parent if these images were made public, we prefer to err on the side of caution. However, microscopy slide images are not personally identifiable, so we may distribute these.
2. Personally identifiable pictures involving adults, because they have the right to consent or not to their likeness being distributed. We consider faces, body profiles, automobile license plates, etc to all be personally identifiable pictures involving adults, especially because these data may be cross-referenced against timestamp, location, clinician, institution, medical condition, other people in the picture, etc.
3. Copyrighted content, which includes images of copyrighted manuscripts, pictures of slideshow presentations, and pictures of any brand or logo. A lab picture that includes boxes bearing logos would be a non-pathology image that we cannot distribute, because we do not have permission to distribute any images with the protected logos. A picture of a powerpoint slide at a conference that shows some text outlining a new way to make a clinical decision would also be a non-pathology image that we hold privately and do not distribute. We similarly hold privately an image of text taken from a non-open-access manuscript because it may not be possible to identify the original source to provide a proper citation, and even if we could, this poses an additional data curation burden that we would rather avoid. Moreover, we prefer to err on the side of caution and not distribute these images, rather than rely on “fair use” or similar law that may expose us to legal challenges and costs^2^. By retaining these images privately, we can train a machine learning classifier to detect these types of images and potentially reduce our manual data curation burden.

#### S5.2.2 Overdrawn rejection criterion

Here we discuss the details of rejecting images as “overdrawn”. Figure S4C *top* is rejected as “overdrawn”, because the regions of interest (ROIs) in the H&E image that the pathologist refers to in the social media post’s text have hand-drawn circles and arrows such that it is not possible to place a 256×256px square over all ROIs without including these circle and arrow marks. We chose 256×256px because deep convolutional neural networks in computational pathology [12] typically require 227×227px (i.e. AlexNet [50] or CaffeNet [51]) or 224×224px (i.e. ResNet [28]) images, and we have used these sizes in the past [16,23]. We note the Inception [52] family of deep convolutional neural networks takes a 299×299px image input, which is larger than 256×256px and is also frequently used in computational pathology [12]. Ideally, each image would have ROIs and hand-drawn arrows/circles annotated at the pixel level, so each image could be annotated as “overdrawn” to arbitrary bounding box sizes, whether 256×256px or 299×299px, and we leave this to future work. Smaller “overdrawn” bounding boxes may allow more images to pass as acceptable, rather than be rejected. A 256×256px image size allows minor rotations and crops for deep learning data augmentation using 224×224px input image sizes. Minor upsampling and/or image reflection at the image’s outer boundaries may allow a 256×256px image to work for 299×299px input image sizes. Figure S4C *bottom* is rejected as “overdrawn”, because this image was originally 783×720px and the arrow marks prevent us from capturing each of the two indicated regions of interest in their own 256×256px square.

#### S5.2.3 Uniform cropping and scaling of original images

Images shared on social media may be any rectangular shape. However, machine learning methods typically require all images be the same size. To accommodate this, we use the following procedure:

1. Take the minimum of two numbers: the original image’s height and width.
2. Crop from the center of the original image a square with a side whose length is the minimum from the prior step.
3. Scale this square to 512×512px.

This square is intended to be large enough to represent small details, such as arrows and circles drawn one pixel wide by the pathologist. Such arrows and circles may then be used to predict if an image is “overdrawn” or not. Ideally, the tweet’s text would be available alongside the image to give the machine learning the fullest information possible about potential ROIs in the image, for “overdrawn” prediction, but for simplicity here we perform only image-based machine learning.

The motivation for the 256×256px “overdrawn” criterion detailed in Sec S5.2.2 is that there may be an attention layer that scans the original image for 256×256px squares that have no marks from the pathologist. Such marks include circles or arrows for ROI indication, or the pathologist’s name to indicate copyright/ownership. Such mark-free 256×256px images may then be used for machine learning on only patient pathology pixels.

### S5.3 Data diversity discussion

#### S5.3.1 Intra-stain diversity

There is an art and variability in histochemical stains that we have not discussed in the main text, but for completeness mention here. We note that in clinical practice we have observed high variability stains, for instance H&E stains that appear almost neon pink, to GMS stains (discussed below) that had silver (black) deposition throughout the slide. One reason for this is that there are a number of reagents that may be used for staining, each with different qualities that can make the stain darker, brighter, pinker, bluer, etc.

IHC stains typically use an antibody conjugated to a brown stain, namely 3,3’-Diaminobenzidine (DAB). The blue counterstain is typically hematoxylin. However, some laboratories conjugate the antibody to a red stain instead. A small minority of our IHC images are this red variant, which should not be confused with H&E.

There is counterstain variability in Grocott’s modification of the Gömöri methenamine silver stain [GMS stain]. Typically the counterstain is green, but a pink counterstain is also available. We may see the pink variant as we acquire more data. Currently we see only green.

#### S5.3.2 Intra-tissue-type diversity

The tissue type hashtags we use are very broad, e.g. #gipath encompasses several organs, such as stomach, small intestine, large intestine, liver, gallbladder, and pancreas. We note, for instance, liver morphology looks nothing like the stomach. Moreover, gynecological pathology, i.e. #gynpath, includes vulva (which looks just like skin, i.e. dermatological pathology, #dermpath), vagina, cervix, uterus, fallopian tubes and ovaries. Again, vulva looks nothing like uterus. A number of tissue features also overlap, such as adipocytes in breast tissue and adipocytes in the subcutaneous fat layer in skin. The amount and distribution of adipocytes typically differs between these tissues however. However, a lipoma in any tissue has a great deal of adipocytes and should not strictly be confused with breast tissue. For all these motivating reasons, we have a future direction to sample every organ within a tissue type hashtag category, for all tissue type hashtag categories.

### S5.4 Procedure overview

#### S5.4.1 Consent, data acquisition, curation, and review

We follow the procedure outlined in Fig 1, and we first obtain data in steps A-D. In step A, we find pathologists on social media (Twitter) who share many pathology cases, or share infrequently shares tissues, e.g. neuropathology. In step B, we contact the pathologist via social media and ask for permission to use their cases. In step C, we download the consenting pathologist’s cases shared on social media. In step D, we manually annotate these posted cases for acceptability, e.g. if overdrawn, corrupt, duplicate, multi-panel, art, or non-pathology rejecting per Fig S4. *Defining an acceptable pathology image* is explained in further sections. We additionally annotate technique (Fig 2A), species (Figs S2, S5A,B,E), and private status (e.g. personally identifiable pictures of adults or pictures of children). *Image data overview* (Sec S5.1) and *criteria for rejected, discarded, private, or acceptable images* (Sec S5.1.1) explain further, e.g. our definition of “overdrawn” or what is [not] pathology. Moreover, if the nontumor/low-grade/malignant status in a tweet is not clear, we read the Twitter discussion thread for this case and manually annotate the case appropriately if possible. Step D also involves clarifying cases that we have trouble annotating, e.g. if it is not clear what stain was used for the image. We first ask the pathologist who posted this case to social media. If we do not obtain an answer from that pathologist, we (i) ask a pathologist at our local institution (i.e. author *S.J.S*.) for an opinion or (ii) ask pathologist coauthors would have shared cases with us. Pathologist validation of tissue and disease labels is an important part of step D, and we use two tools for this. The first tool is the Interactive Pathology Annotator (IPA) (Figs S6, S7), which pathologists may run on their desktop to browse their case annotations. The second tool is our social media bot “pathobot” (Fig 1), which interacts with collaborating pathologists, then publicly posts results of search and disease state predictions. Pathobot search results may indicate annotation issues, e.g. if a bone and soft tissue pathology search returns a breast pathology result, we may check if the breast result was mistakenly labeled as bone and soft tissue. Though we sometimes manually annotate some cases, most cases are annotated in a crowd sourced fashion. We use social media post hashtags and a pathologist-reviewed rule-based text processing algorithm to determine tissue type and disease state (Fig S8).

##### Interactive Pathology Annotator discussion

For completeness, we show another example of the use of our Interactive Pathology Annotator (IPA) tool (Fig S7), which some pathologists have used to check that tissue and disease state annotations were correct. This is a case of metastatic disease, from breast to gastrointestinal tissue, showing a diffuse pattern of lobular carcinoma that is more common in breast.

**Fig S6:**
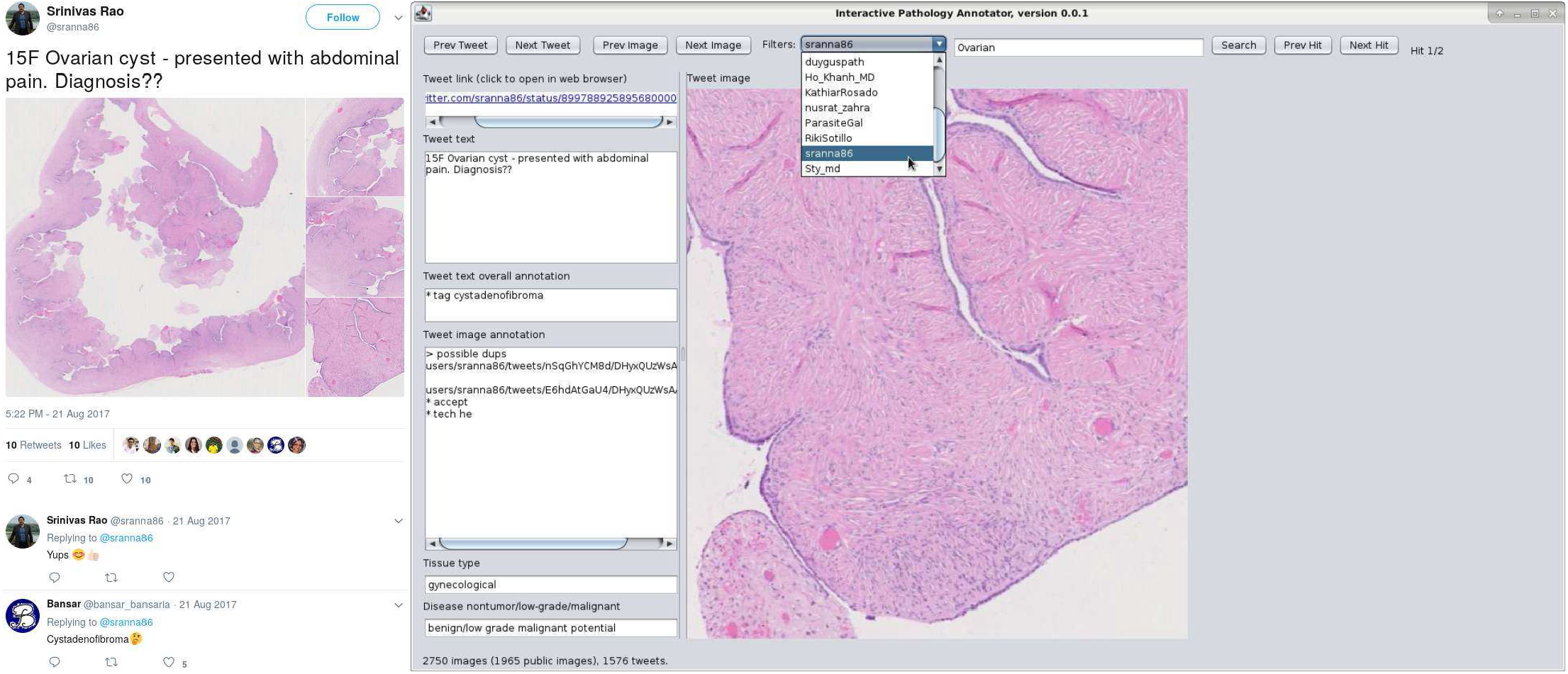
Interactive Pathology Annotator tool and social media dialogue. *At left*: Pathologist (author *S.R.A.*) discusses a case. Without mentioning the diagnosis himself, he confirms the diagnosis suggested by a second pathologist, i.e. cystadenofibroma, which we explicitly annotate. *At right*: Our Interactive Pathology Annotator (IPA) tool displays an image from this case, in the context of the tweet overall. IPA is a portal for pathologists to (i) browse tweets and images in the dataset; (ii) validate our data annotations; (iii) check our tissue type categorization algorithm results, (iv) check our nontumor, low grade, and malignant categorization algorithm results; (v) search tweets for specific keywords or diagnoses; (vi) filter out all cases except those from a specific pathologist; and (vii) click the link to the original tweet on Twitter for context.

#### S5.4.2 Machine learning, search, checks, and social media bot

Our procedure (Fig 1) continues with data analysis in steps E-G. In step E, we use machine learning to train a classifier for a supervised learning task. For example, a task may be to predict the disease state evident in a H&E image: malignant, benign / low grade malignant potential [low grade], or nontumoral pathology. This is a three-class classification task. Our baseline classifier is a Random Forest, [31] which we compare to deep learning. We reuse the classifier to compute a similarity metric for search. In step F, a pathologist posts to social media three key pieces of information together: (i) pathology images, (ii) text descriptions, and (iii) the text “@pathobot”. Our social media bot is triggered when mentioned this way, with parts (i) and (ii) forming the pathology search query. In step G, our bot first searches its social media database of cases, then searches its larger PubMed database. A search result will be ranked highly when the query and result tissue types match, when more clinical keywords are shared between the query and result, and when the images are similar. Additionally, the bot will use an ensemble of classifiers to compute with uncertainties the probability of each disease state in each image. This prediction is a sanity check for search, i.e. if the prediction is uncertain or inaccurate, then the search results may be suspect.

#### S5.4.3 Social media interactions, search, notifications, recruitment

One cycle of our procedure (Fig 1) culminates with concluding social media interactions in steps H and I, before ultimately repeating at step A. The social media bot posts its

**Fig S7:**
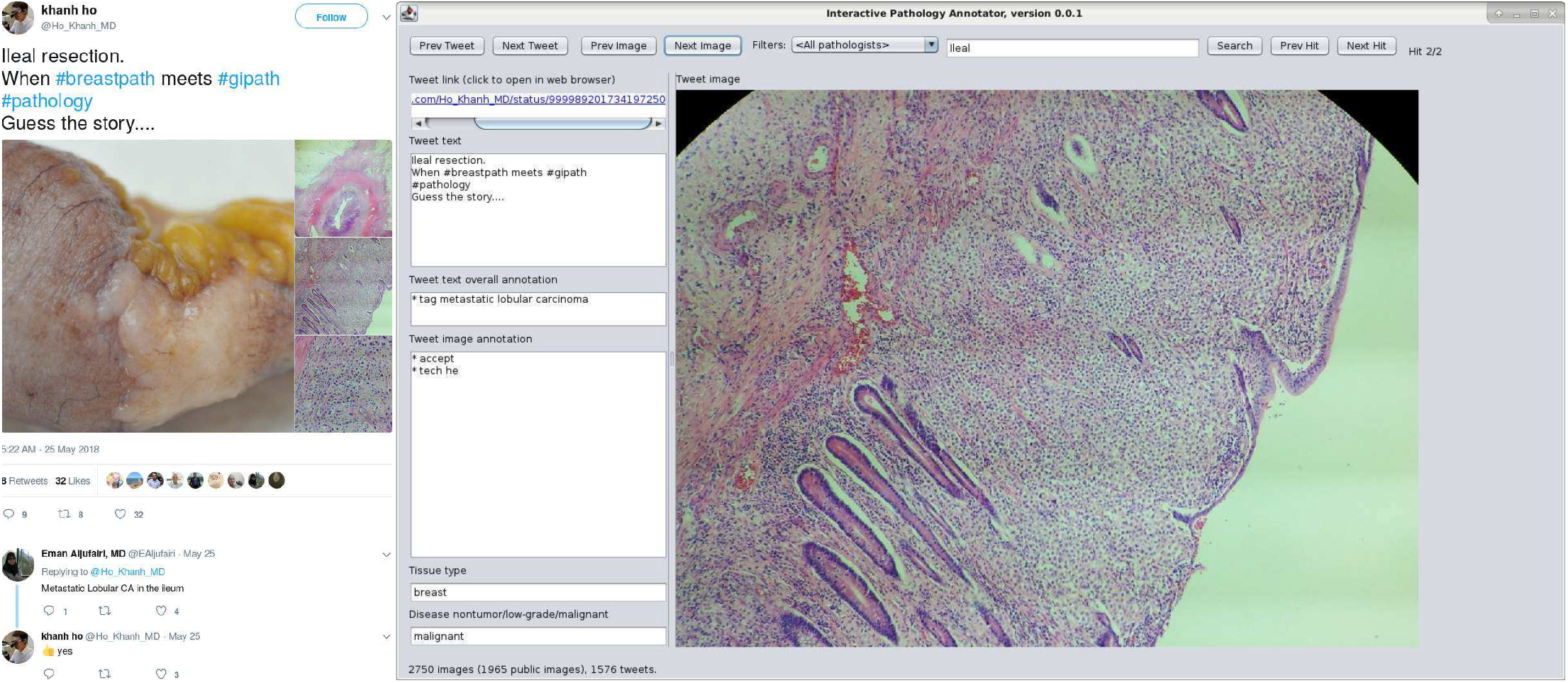
Interactive Pathology Annotator tool and tissue type hashtags. *At left*: Pathologist (author *K.H.)* discusses case. Without mentioning the diagnosis himself, he confirms diagnoses suggested by other pathologists, i.e. lobular breast carcinoma metastasized to ileum, which we explicitly annotate. *At right*: IPA shows that our tissue type categorization algorithm categorizes this tweet as breast pathology rather than gastrointestinal. The primary tumor is in breast. We define the tissue classification task this way to have applications for tumor site of origin prediction.

social media search results, PubMed search results, and disease state prediction results. The social media search results include links to similar cases posted to social media. The social media platform may notify pathologists that their posted case has been linked. These pathologists may discuss the putatively similar case. Our bot leverages text information from the pathologist’s search post and reply posts. In this way our bot’s search is informed by any diagnosis in the differential from any replying pathologist. When multiple pathologists mention the same clinical or diagnostic keywords, those keywords are weighted more highly for search. In effect, search is a collective endeavor by all pathologists in the community discussing the case. The same search repeated over time may be more informed when more pathologist discussion has accumulated over that time. We find that integrating our bot into social media discussions sometimes inspires pathologists to contact us, share with us, and collaborate with us. We then return to step A, and we collect more data for search and classifier training.

### S5.5 Text data overview

For supervised learning, we use regular expressions to detect keywords in a tweet’s text, to determine labels for the tweet’s images. The text and included hashtags may indicate tissue type or disease state.

#### S5.5.1 Tissue type categories from text

Prior work has discussed pathology-related hashtags as a way to make pathology more accessible on social media [53]. Pathologists use hashtags to indicate histopathology tissue types, such as “#gynpath” to indicate gynecological pathology (Fig 2B). Sometimes alternative spellings are used, such as “#ginpath”. Abbreviations are also common, e.g. “#breastpath” and “#brstpath” all mean the same thing: breast pathology (Fig S7). A pathology hashtag ontology is available at https://www.symplur.com/healthcare-hashtags/ontology/pathology/. Because a tweet can have more than one hashtag, we took the first tissue type hashtag to be the “primary” tissue type of the tweet, and ignored the others. Section S5.6.1 discusses a special case. As detailed in Section S5.6.2, we used hashtags and keywords for all tweets in a message thread to identify the ten tissue types on Twitter, finding 233 bone and soft tissue tweets, 155 breast tweets, 415 dermatological tweets, 794 gastrointestinal tweets, 239 genitourinary tweets, 218 gynecological tweets, 308 head and neck tweets, 115 hematological tweets, and 559 pulmonary tweets.

**Fig S8:**
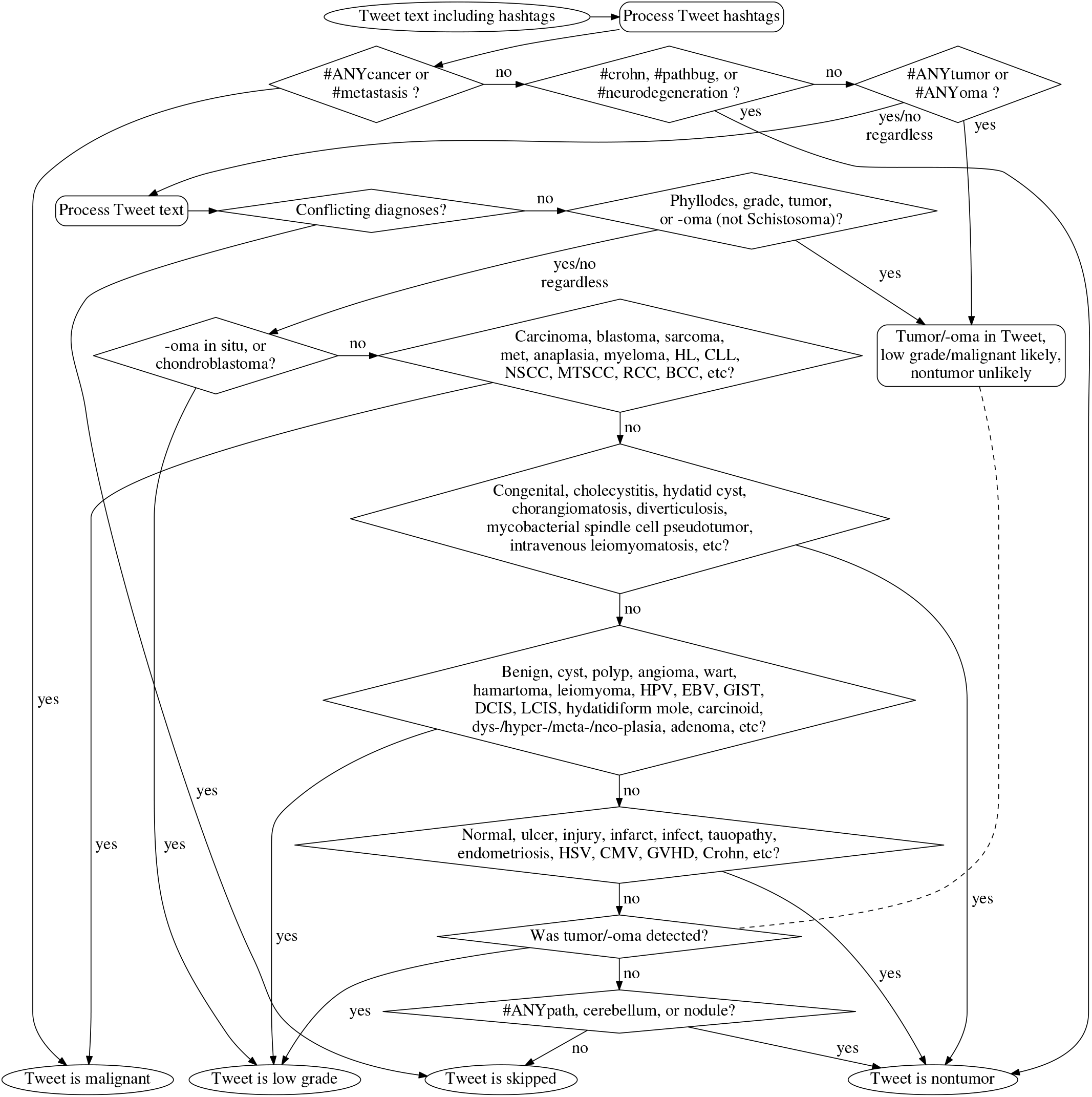
3-disease text processing algorithm flowchart. Flowchart of algorithm that processes a single tweet’s text to categorize it as nontumor (309 images), benign/low grade malignant potential [low grade] (347 images), or malignant (385 images). A tweet may be skipped (132 images, i.e. 11.3% of images) when the pathologist discusses multiple possible diagnoses for this case or when no pathology keywords are found. Dashed line indicates early steps where tumor/-oma detected, and a later step where detected tumor/-oma considered for possible low grade categorization. Nontumor, low grade, and malignant are defined in Sec S5.7. Flowchart steps are detailed in Sec S5.7.1. The algorithm has many steps in order to parse overlapping words that have different diagnoses. For instance, if “Lobular carcinoma in situ of the breast” (which is a low grade disease) was the tweet text, the algorithm has an early step to categorize “carcinoma in situ” as low grade (which is correct here) because a later step categorizes “carcinoma” as malignant (which is not correct here). Indeed, tweet text “Carcinoma of the breast” describes a malignant disease and the algorithm categorizes it malignant because “in situ” is absent. Besides “carcinoma in situ” (low grade) and “carcinoma” (malignant), the algorithm distinguishes “chorangiomatosis” (nontumor) from “angioma” (low grade), “hydatidiform mole” (low grade) from “hydatid cyst” (nontumor) from “ovarian cyst” (low grade) from “cholecystitis” (nontumor), and “intravenous leiomyomatosis” (nontumor) from “leiomyoma” (low grade).

#### S5.5.2 Nontumor, low grade, and malignant categories from text

We define three broad disease state categories (Figs 2C, S8) to use as labels for supervised learning. Our “nontumor” category of 589 tweets includes normal tissue, artifacts, injuries, and nontumoral diseases, e.g. Crohn’s disease, herpes simplex infection, and myocardial infarction. Our “malignant” category of 1079 tweets includes all malignant disease, including carcinoma, blastoma, sarcoma, lymphoma, and metastases. Our definition of malignancy in epithelial cancers is the ability to breach the basement membrane, i.e. a malignant tumor escapes containment and is therefore no longer treatable with surgical resection. Our “pre-neoplastic/benign/low grade malignant potential” [low grade] category of 919 tweets is then all tumors or pre-cancer/neoplastic lesions that are not yet invasive/malignant, e.g. hamartomas, carcinoid tumors, adenomas, and carcinoma *in situ.* Details in Section S5.7. For the nontumoral vs low grade vs malignant task, text processing was more complicated than the tissue type task (Sec S5.6.2) because (i) of a heavy reliance on diagnosis keyword matching (flowchart in Fig S8), and (ii) additional per-tweet and per-image annotations to clarify nontumor/low-grade/malignant state, which may involve feedback from a pathologist. Details in Section S5.7.

### S5.6 Supplementary Text processing

#### S5.6.1 Hashtag special case

A hashtag special case is “#bstpath”, bone and soft tissue pathology, which we include in our breast pathology category only when the social media post’s text also includes the word “breast” or other breast-related keywords. Such keywords are listed further below in this subsection. Examples of such tweets are *“Pleomorphic lobular carcinoma of the breast: Beautiful cells but nasty tumour #pathology #pathologists #BSTPath”* and *“Now at my desk, W(Ą7y-o) breast nodule… Could be it siliconoma?? But it isn’t noted giant cells #pathology #pathologists #BSTpath”.*

#### S5.6.2 Tissue hashtags and keywords

We found a large number of pathology-related hashtags. We grouped alternative spellings, e.g. #ginpath is #gynpath, #brstpath is #breastpath, and #headandneckpath is #entpath. We also grouped less common hashtags with more common hashtags, e.g. #cardiacpath is #bstpath (bone and soft tissue). Some groupings were broad, e.g. #headandneckpath, #thyroid, #salivary, #oralpath, #endocrine, #endopath, #oralpath, #eyepath, and #ocularpath are all #entpath.

To expand the per-tissue tweet counts, we moved beyond the hashtags and next searched for keywords in the tweet using Perl regular expressions. Further, if a tweet’s tissue type could not be determined by hashtags and keywords, we assigned the tissue type of any other tweet in the message thread of tweets. For example, if a tweet of unknown tissue type were a reply to a tweet of known genitourinary type, then we considered both tweets to be genitourinary.

### S5.7 Nontumoral, Low grade, and Malignant task details

Tasks involving distinguishing nontumoral disease, low grade tumors, and malignant tumors (Fig 2C) are our most difficult tasks. The acknowledged definition of “malignant” in epithelial cancers is the ability to breach the basement membrane, i.e. a malignant tumor escapes containment and is therefore no longer “treatable with surgical resection”. A malignant tumor can invade into the adjacent tissue, lymphatics, and blood vessels. For machine learning, we define a three categories of disease: (a) normal tissue and nontumoral disease; (b) benign, low grade, and oncovirus-driven tumoral disease; and (c) malignant tumors – but there are number of caveats with this, because:

1. there is a spectrum of pathology rather than an oversimplified 3-class nontumoral/low-grade/malignant system.
2. the benign/malignant dichotomy may be more vague in certain tissues e.g. central nervous system (CNS) primary tumors such as chordomas.
3. vague terms like adenoma are typically benign but may be malignant, and likewise vague terms like anaplasia are more often associated with malignancy but not always.
4. vague terms like anaplasia and neoplasia make no real reference to the malignancy of lesions i.e. there are benign anaplastic lesions, while neoplasia is almost synonymous with tumor.
5. terms like tumor do not provide information about benign or malignant state, though normal/nontumoral can be ruled out.
6. there may be some disagreement if some terms, e.g. “carcinoma *in situ* “, are more appropriate to include as low grade, or if instead should be considered malignant due to their malignant potential or treatment implications. For instance, ductal carcinoma *in situ* (DCIS) typically needs to be removed with surgery or radiotherapy, whereas lobular carcinoma *in situ* (LCIS) typically does not. DCIS’s lower grade counterpart, atypical ductal hyperplasia, may get surgery or not. We believe treatment implications are a separate task. Typically, tweets do not include a decision to perform surgery or not, so additional annotations may be needed for the surgery task. We assign all pre-cancer and tumoral disease with malignant potential to the “low grade” category, in light of these benign/malignant ambiguities and data limitations.
7. the diagnosis should be known before deciding benign/malignant, but it is very difficult to know the full diagnosis from the brief, generic, descriptive terms in the tweet.

#### S5.7.1 Text processing for Nontumoral, Low grade, and Malignant tasks

To determine if an acceptable H&E human microscopy image is nontumoral, low grade, or malignant, we use regular expressions (Fig S8) as we did for tissue type classification. However, keywords differed and we considered all tweets in a message thread per Sec S5.5.1. To infer these message threads of tweets, we downloaded from Twitter each tweet’s metadata (in JavaScript Object Notation (JSON) format), which describes the parent tweet for each tweet. If tweet A is a reply to tweet B, then tweet A is the parent of tweet B, and both tweets are in the same message thread.

Our heirarchical algorithm for nontumor/low-grade/malignant keyword-matching shown in Fig S8, and details for each step follow. First, to determine if a single tweet indicated nontumoral, low grade, or malignant, we looked for specific hashtags in a tweet’s text that indicated malignancy, tumoral status, or nontumoral status. For illustartion, what follows is a subset of our rules, with commentary.

1. Malignant: `/#[a-z] cancer/i` or `/#metastas[ei]s/i`

- The first regular expression in this set matches #ANYcancer, where ANY can be any non-whitespace characters, e.g. “#bladdercancer” and “#breastcancer” both match, as well as “#cancer”.
- Metastasis is a sign of malignant cancer, so tweets with #metastasis or #metastases hashtags are malignant.
- If any matching keyword is detected, no further keyword processing is performed. The tweet is malignant.
2. Nontumoral: `/#crohn/i` or `/#neurodegeneration/i` or `/#pathbug/i`

- Crohn’s disease and neurodegeneration are not tumoral diseases, so this tweet is in the nontumoral/normal category. This `/#crohn/i` regular expression is case-insensitive, so it matches “#crohn”, “#Crohn”, and “#CROHN”. The #pathbug hashtag indicates a parasite or other microorganism is in the image, which is also nontumoral.
- If any matching keyword is detected, no further keyword processing is performed. The tweet is nontumoral.
3. Tumoral status (ambiguously low grade or malignant): `/#[a-z]*tumou?r/i` or `/#[a-z]*oma/i`

- The first regular expression in this set matches #ANYtumor or #ANYtumour, where ANY can be any non-whitespace characters, e.g. “#BrainTumor” and “#phyllodestumour” both match, as well as “#tumor”.
- The second regular expression matches #ANYoma, e.g. #Lymphoma and #leiomyoma both match.
- Because “tumor” and “-oma” do not necessarily mean a tumor is low grade or malignant, further keyword matching is performed. It is unlikely that the tweet is nontumoral. If no other specific information is found after all further keyword matching is performed, the tumor is presumed to be low grade.

Second, if no hashtags matched, we then analyzed keywords in the tweet text.

1. Skip: `/mistake/i` or `/misinterpret/i` or `/confuse/i` or `/suspect/i` or `/worry/i` or `/surprise/i` or `/mimic/i` or `/simulate/i` or `/lesson/i` or `/\bhelp\b/i` or `/usually/i` or `/difficult/i` or `/pathart/i` or `/pathchallenge/i` or `/pathquiz/i` or `/pathgame/i` or `/^http/`

- We skip tweets where (i) the pathologist discusses points of the case which may be easily mistaken – instead of providing a single diagnosis, (ii) the pathologist provides a diagnosis but may suspect an alternative diagnosis, or (iii) the tweet is simply a link to another tweet. No further keyword matching is performed for this tweet.
2. Tumoral status (ambiguously low grade or malignant): `/phyllod/i` or `/\bgrade\b/i` or `/tumou?r/i` or (`/[a-z]{3,}oma\b/i` and not `/schistost?oma/i`)

- Phyllodes tumors, mentions of “tumor” or “tumour”, mentions of tumor “grade”, and mentions of words that end in “oma” but are not “Schistosoma” – are all detected here.
- Loosely speaking, phyllodes tumors are only slightly more likely to be low grade than malignant. Because “tumor”, “-oma”, and “grade” do not necessarily mean a tumor is low grade or malignant, further keyword matching is performed. It is unlikely that the tweet is nontumoral. If no other specific information is found after all further keyword matching is performed, the tumor is presumed to be low grade.
- *Schistosoma* (and its misspelling “Schistostoma”) refers to a genus of parasitic worm, rather than a tumor, though *Schistosoma* ends in “oma” like many tumor types.
3. Low grade: `/oma in situ/i` or `/chondroblastoma/i`

- If we did not skip this tweet, but the tweet does mention “oma *in situ*’’, e.g. “carcinoma *in situ*” or “melanoma *in situ* “, then we consider this tweet and images to represent low grade disease. Carcinoma *in situ* is pre-cancer, and we consider it more low grade than malignant. If a tweet contains only “carcinoma” but not “*in situ*”, subsequent steps will consider the tweet as malignant.
- If the tweet includes “chondroblastoma”, this tweet is low grade. This is not to be confused with other blastomas, such as glioblastoma or lymphoblastoma, which are malignant and matched in subsequent steps.
- No further keyword matching is performed if these patterns match. The tweet is low grade.
4. Malignant: `/malignant/i` or `/malignancy/i` or `/cancer/i` or `/\bCA\b/i` or `/carc?inoma/i` or `/sarcoma/i` or `/blastoma/i` or `/\bWilms/i` or `/GBM/i` or `/anaplas(?:ia|tic)/i` or `/metastas[ie]s/i` or `/metastatic/i` or `/\bmets?\b/i` or `/adenoca/i` or `/melanoma/i` or `/seminoma/i` or `/lymphoma/i` or `/leuka?emia/i` or `/mesothelioma/i` or `/myeloma/i` or `/hodgkin/i` or `/\bHL\b/i` or `/burkitt/i` or `/plasmoc[yi]toma/i` or (*/paget/i* and `/breast/i`) or `/\bCLL\b/i` or `/PCNSL/i` or `/NSCHL/i` or `/\bCHL\b/i` or `/NSCC/i` or `/\bI[LD]C\b/i` or `/\bASPS\b/i` or `/mtscc/i` or `/sq?cc/i` or `/rcc/i` or `/bcc/i`

- Many diagnoses and abbreviations may indicate cancerous malignancy, e.g. carcinoma, sarcoma, Wilms’ tumor, leukemia, RCC [renal cell carcinoma], NSCC [non-small cell lung carcinoma], or the stand-alone abbreviation “CA” [cancer].
- We consider “anaplastic/anaplasia” to be more malignant than low grade disease.
- No further keyword matching is performed if these patterns match. The tweet is malignant.
5. Nontumoral: `/congenital/i` or `/cholecystitis/i` or `/chorangiomatosis/i` or `/mycobacteri(?:um|al)\s*spindle\s*cell\s*pse?udotumor/i` or `/intravenous\s*leiomyomatosis/i` or `/helicobacter/i` or `/dirofilaria/i` or `/tuberculo/i` or `/enterobius/i` or `/echinococcus/i` or `/hydatid\s*cyst/i` or `/giardia/i` or `/cryptosporidium/i` or `/ascaris/i` or `/sarcina/i` or `/worm/i` or `/spiroquet(?:osis|es)/i` or `/diverticulosis/i` or `/villitis/i` or `/colitis/i` or `/gastritis/i` or `/esophagitis/i` or `/appendicitis/i` or or `/xanthoma/i`

- Many diagnoses and abbreviations may indicate nontumoral disease, e.g. congenital conditions, *Helicobacter* infection, and villitis. Nontumoral disease keywords that contain “cyst”, e.g. “cholecystitis” and “hydatid cyst”, are detected here, because subsequent keyword matching steps will detect “cyst” as a sign of low grade tumoral disease.
- If one of these nontumoral keywords matches, no further keyword matches are attempted, and the tweet is considered nontumoral, even if prior steps detected “tumor” or “-oma”. For instance, a “xanthoma” is a lipid aggregate, not a tumoral disease, even though xanthoma ends in -oma.
6. Low grade: `/benign/i` or `/cyst/i` or `/polyp/i` or `/hamartoma/i` or `/chorangioma/i` or `/ha?ematoma/i` or `/cylindroma/i` or `/fibroma/i` or `/luteoma/i` or `/c[yi]toma/i` or `/cond[yi]loma/i` or `/neoplas(?:ia|tic|m)/i` or `/LCIS/i` or `/DCIS/i` or `/\b[LD]IN\b/i` or `/lipoma/i` or `/carcinoid/i` or `/neuroma/i` or `/meningioma/i` or `/perineurioma/i` or `/cavernoma/i` or `/\bLGG\b/i` or `/\bODG\b/i` or `/oligodendroglioma/i` or `/craniopharyngioma/i` or `/le[yi]om[iy]oma/i` or `/schwannoma/i` or `/osteochondroma/i` or `/ependymoma/i` or `/angioma/i` or `/syringoma/i` or `/acanthoma/i` or `/collagenoma/i` or `/hidradenoma/i` or `/papilloma/i` or `/pilomatrixoma/i` or `/hydatidiform\s*mole/i` or `/wart/i` or `/molluscum/i` or `/\bHPV\b/i` or `/\bEBV\b/i` or `/kerat?osis/i` or `/fibrokeratoma/i` or `/melanoc[iy]tosis/i` or `/brenner/i` or `/granular\s+cell\s+tumou?r/i` or `/metaplas(?:ia|tic)/i` or `/dysplas(?:ia|tic)/i` or `/dysembryoplas(?:ia|tic)/i` or `/hyperplas(?:ia|tic)/i` or `/\bLFH\b/i` or `/\bDNE?T\b/i` or `/\bNET\b/i` or `/\bPTC\b/i` or `/\bGIST\b/i` or `/\bSTIC\b/i` or `/\b[LD]ISN\b/i` or `/adenoma/i` or `/adenosis/i`

- Many diagnoses may indicate benign tumor, e.g. hamartoma, fibroma, condyloma, papilloma, lipoma, adenoma, adenosis, or cyst.
- We consider “neoplastic/neoplasia”, “metaplastic/metaplasia”, “hyperplastic/hyperplasia”, and “dysplastic/dysplasia” to be more indicative of benign/low-grade/non-invasive/pre-malignant disease than malignant disease, but these terms are vague.
- We broadly consider oncovirus-driven tumors and wart-like growths to be in this low grade category also, e.g. HPV [human papilloma virus] warts and *Molluscum contagiosum* “water warts”.
- We similarly consider abbreviations “LCIS” [lobular carcinoma *in situ*], “DCIS” [ductal carcinoma *in situ*], “LISN” [lobular *in situ* neoplasia], and “DISN” [ductal *in situ* neoplasia] to be more benign than malignant disease, so we categorize them as low grade. Though DCIS may require surgical or radiological intervention to be removed while LCIS may not, we consider our “low grade” and “malignant” categories to be defined by the apparent histopathology rather than the appropriate medical intervention. Predicting appropriate medical intervention would be a different machine learning task.
- If one of these keywords match, the tweet is considered low grade and no further keyword matching is performed.
7. Nontumoral: `/normal/i` or `/ulcer/i` or `/embolism/i` or `/thromb/i` or `/rupture/i` or `/infarct/i` or `/aneurysm/i` or `/ha?emorrhag/i` or `/injur(?:y|ed)/i` or `/inflam/i` or `/swell/i` or `/balloon\s*cell\s*na?ev(?:us|i)/i` or `/decidua/i` or `/foreign/i` or `/lymphadenopath?y/i` or `/vasculopathy/i` or `/vasculitis/i` or `/synovitis/i` or `/pulmonary\s*interstitial\s*glycogenosis/i` or `/essential\s*thrombocythemia/i` or `/endometriosis/i` or `/mastoc[iy]tosis/i` or `/castleman/i` or `/herpe(?:s|tic)/i` or `/\bHSV\b/i` or `/\bCMV\b/i` or `/cytomegalovir/i` or `/viral/i` or `/bacteri(?:a|um)/i` or `/fung(?:al|us)/i` or `/mycetoma/i` or `/myco(?:sis|tic)/i` or `/infect(?:ion|ed)/i` or `/tauopathy/i` or `/amyloidosis/i` or `/neurodegen/i` or `/\brabies\b/i` or `/hemosiderosis/i` or `/polymicrogyria/i` or `/status\s*verrucosus/i` or `/\bIUGR\b/i` or `/storage\s*dis(?:ease|order)/i` or `/athero(?:sis|ma)/i` or `/atherosclero(?:sis|tic)/i` or `/gauzoma/i` or `/colchicine/i` or `/\bIBD\b/i` or `/GVHD/i` or `/crohn/i`

- Many diagnoses may indicate normal tissue of nontumoral disease, e.g. normal, embolism, decidua, tauopathy, foreign body, mycetoma, CMV [cytomegaolovirus] infection, GVHD [graft versus host disease], and Crohn’s disease.
- If one of these nontumoral keywords matches, no further keyword matches are attempted, and the tweet is considered nontumoral, even if prior steps detected “tumor” or “-oma”. For instance, a mycetoma is not a tumor, even though mycetoma ends with -oma.
8. Nontumoral: (not tumor/oma) and (`/#[a-z]*path/i` or `/cerebell(?:um|ar)/i` or `/nodul`(*?:e|arity*)/*i*). Low grade if tumor/oma.

- If the tweet does not have tumor or “-oma” keywords detected from prior steps, and if the tweet has a #ANYpath hashtag (e.g. “#pulmpath” or “#pathology”), mention of “nodule”/“nodularity”, or mention of the cerebellum, then we consider the tweet to be nontumoral. If instead the tweet has tumor or -oma keywords, then we consider the tweet to be low grade. The tweet is skipped if no steps identified the tweet as nontumoral, low grade, or malignant.
- Cerebellum is mentioned in several tweets, e.g. to depict normal cerebellar tissue^3^. Currently, we group normal tissue with tissue having nontumoral disease. We expect more tissue-based keywords may be used here in the future, as we expand our study to include more pathologists, tissues, and normal cases.
- In practice, we manually inspect all tweet message text to minimize the number of cases that are classified as nontumoral here. We typically write regular expressions to match specific keywords that indicate if a tweet represents nontumoral, low grade, or malignant disease.
- As part of our manual data curation, if on Twitter there was discussion among pathologists, and a different pathologist mentioned a correct diagnosis, and our consenting contributing pathologist concurred, then we write an auxiliary annotation file for the tweet with a summarized diagnosis^4^. This summary is also used for pattern matching. This is an additional way that we minimize how many cases are handled at this late step.
- Moreover, if the contributing pathologist wrote diagnostic text directly in the image, we will write this text in the auxiliary annotation file for text matching also.^5^
- The way this “default nontumoral or low grade” rule is intended to be used is as a catch-all for unusual but non-malignant conditions^6^. Our motivation for this rule is to minimize our manual data curation burden. We do not wish to write an auxiliary annotation file or make a new regular expression for each unusual type of case, and we observe many of these cases are not malignant. It remains important to inspect the cases manually for correctness.

**Fig S9:**
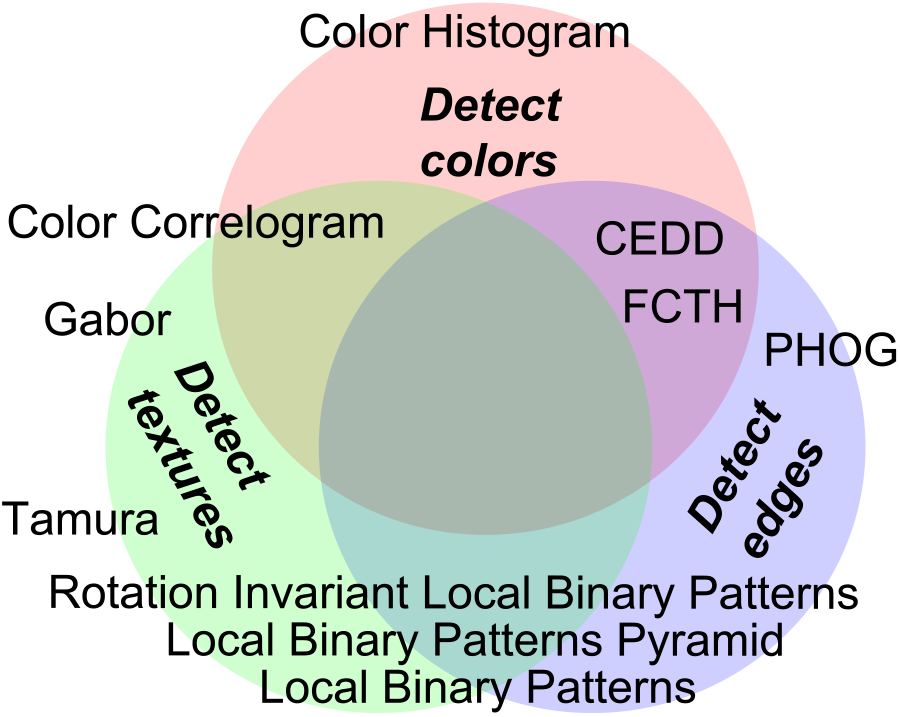
Machine learning features. We use a variety of color, texture, and edge features for baseline machine learning analyses. Some features, such as color histograms, detect only color. Other features, such as Color Correlograms, detect both colors and textures. Pyramid features are scale-invariant. We separately consider SIFT features, which detect edges in a scale-invariant, rotation-invariant, and color-invariant manner, localized at interest points in an image.

Tweets that do not match any nontumoral, low grade, or malignant rules are skipped in the same manner that Tweets matching skip rules are skipped. An additional caveat is this keyword matching may need refinement as we accumulate data.

### S5.8 Image features for machine learning

To perform baseline machine learning analyses on the images from social media, we derive a feature representation for each image, as follows. We crop each image to the center square and resize it to 512×512 pixels [px]. See Sec S5.2.3 for more discussion of the 512×512px image size and how it relates to the 256×256px image size for the “overdrawn” criterion. This 512×512px image is then converted to a feature vector of 2,412 dimensions. The features we use (Fig S9) are available in Apache LiRE [54]. These features, and their dimension counts, are as follows: CEDD (144) [55], Color Correlogram (256) [56], Color Histogram (64) [54], FCTH (192) [57], Gabor (60) [54], Local Binary Patterns (256) [58], Local Binary Patterns Pyramid (756) [59], PHOG (630) [60], Rotation Invariant Local Binary Patterns (36) [61], and Tamura (18) [62].

#### S5.8.1 Deep learning instance and set feature vectors

After training, our appended 100-neuron layer (Fig 3B) is a 100-dimensional disease feature representation for a 224×224px patch. Due to our custom activation function and regularization (Fig S13), these 100 features are approximately binary (Fig S14C). A vector sum of these 100-dimensional approximately binary feature vectors is a 100-dimensional feature counter vector, which we relate to set cardinality (Eqn 8). Inspired by Deep Sets [63] and continuous bag-of-words [64] methods, we sample 21 patches throughout the white-balanced image, then add the 21 100-dimensional feature vectors to derive a 100-dimensional set representation for the white-balanced image (Fig 3C). In the set representation, if a feature’s value is 21, then all 21 image patches have this feature. An approximate intuition follows that if the value is 20, then 1 of the 21 patches does not have this feature. This set-based approach offers limited interpretability and facilitates learning on large pathology images on social media, despite the receptive field of a deep neural network being much smaller (1 million pixels versus 224×224px, respectively). We train a Random Forest on the concatenation of (i) this 100-dimensional set representation, (ii) our 2,412-dimensional hand-engineered feature vector, (iii) the 10-dimensional tissue type covariate vector, and (iv) the 1-dimensional marker mention covariate vector (Fig 3C). Like Deep Sets, we use deep learning for instance learning, and add instance representations for a set representation. However, our approach differs from Deep Sets in that (i) we use a Random Forest to learn on set representations and side information (which is not differentiable end-to-end), (ii) we add approximately binary Centered Soft Clipping (Fig S13) features in the range (0,1) to implement counter-like set representations rather than add ReLU features in the range [0,∞) which do not necessarily count instance features, and (iii) we use Random Forest feature importance to interpret the relative influence of deep features, hand-engineered features, and optional covariates on prediction/classification (Fig 4).

### S5.9 Machine learning sanity checking for search

#### S5.9.1 Prediction uncertainty quantified with ensemble

For disease state predictions posted to social media by the bot, we use an ensemble of classifiers to quantify prediction uncertainty. This ensemble consists of the set of classifiers for leave-one-pathologist-out precision@k search testing, so the ensemble size is equal to the number of pathologists who contributed data. Leaving one pathologist out from each classifier’s training ensures some variability between classifiers. One classifier makes one prediction per image. We use a Z-test to determine if a disease state prediction’s mean is significantly above chance. If no prediction is above chance, then the predictions may be due to chance alone and ignored by a pathologist. Likewise, search results may be ignored. This is our first prospective sanity check, and only requires a statistical interpretation of predictions.

##### Training data detection with ensemble

This leave-one-out ensemble provides an additional non-prospective sanity check – if only one classifier in the ensemble makes a prediction that is strongly different from all the other classifiers in the ensemble, it is possible that the bot was requested to make a prediction on a case that was in the training data. This one “outlier” classifier that makes the strongly different prediction is the classifier that left this pathologist’s case out for training. The distribution of predictions from these classifiers is depicted in a boxplot posted to social media by our bot (Fig 1 *at lower left*). An outlier is indicated by a circle in the boxplot, and a circle in a strongly different direction that the boxplot’s interquartile range may suggest the “outlier classifier sanity check” has been encountered.

#### S5.9.2 Classifier repurposed, so if prediction suspect, then search suspect

If the Z-test of our distribution of predictions indicates the evidence in favor of a particular disease state, e.g. nontumor, is not due to chance alone, but this is surprising to a pathologist’s expectations, the pathologist may consider the prediction to be suspect, so search results may be suspect as well. This is our second prospective sanity check, which requires a pathologists’s expert opinion.

##### Whole-patient prediction and disagreement detection

Our method calculates the probability of a disease state per image, but given 1-4 images are in a tweet, we are left with two questions. First, what is the overall probability that this patient has a particular disease state, e.g. malignant? For this, we make the naïve statistical independence assumption of probabilities, multiply the prediction probabilities, and normalize them to sum to one. Second, what should we do when the prediction for one image differs from another for a given patient? In this case, the bot includes a warning in its tweet message and suggests mistakes are more likely, but we do not consider this a prospective sanity check, because images could indeed show different disease states.

#### S5.9.3 Deep learning prediction heatmaps comparable to pathologist expectations

Heatmaps from our deep learning can localize disease states within an image (Figs 5, S11), offering our third prospective sanity check to pathologists. If localization of predicted disease is suspect, then prediction may be suspect, and search may be suspect.

### S5.10 Machine learning interpretability for search

#### S5.10.1 Hand-engineered feature interpretability

We use existing hand-engineered visual features extensively (Fig S9). *Image features for* machine learning (Sec S5.8) discusses the combination of color, texture, and edge features we use (Fig S9). All of have human-defined mathematical or algorithmic behavior written in software code. We know by definition a color histogram feature is invariant to rotation, because such a feature may simply be the sum of each pixel’s red value in an image. Similarly, rotating an image should not change the diagnosis, so rotation invariance makes sense for disease state prediction. We also know properties of other features, such as the important Local Binary Patterns Pyramid (LBPP) [59]. LBPP is globally color-invariant because it operates on grayscale pixel values, not color pixel values. This may provide robustness to staining protocol differences between institutions. LBPP is globally scale-invariant because it employs a pyramid for multi-scale representation. This is the same pyramid used by PHOG [60]. This pyramid may support robust machine learning despite pathologists sharing images at different magnifications. LBPP is locally rotation-invariant because it consists of rotation-invariant local binary patterns at every level of the multi-scale pyramid representation. These rotation-invariant local binary pattern features are locally robust to localized orientation changes of a pathology image, e.g. minor perturbations in the orientation of a cell or tissue fiber. In contrast, PHOG consists of oriented gradients at every pyramid level, rather than rotation-invariant local binary patterns. LBPP is not globally rotation-invariant because, like PHOG, most pyramid grid cells are spatially localized, so a rotated image will have a different feature representation. LBPP is a texture feature because it compares the value of a center pixel to the value of many pixels at a particular radius from the center. Prior groups have used texture features to distinguish stroma, lymphocytes, necrosis, etc. Hand-engineered feature interpretability provides a simple foundation on which to build more abstract levels of interpretability. We can also reason about what features do not improve disease state prediction, e.g. SIFT features, which are thought to cover nuclei.

**Fig S10:**
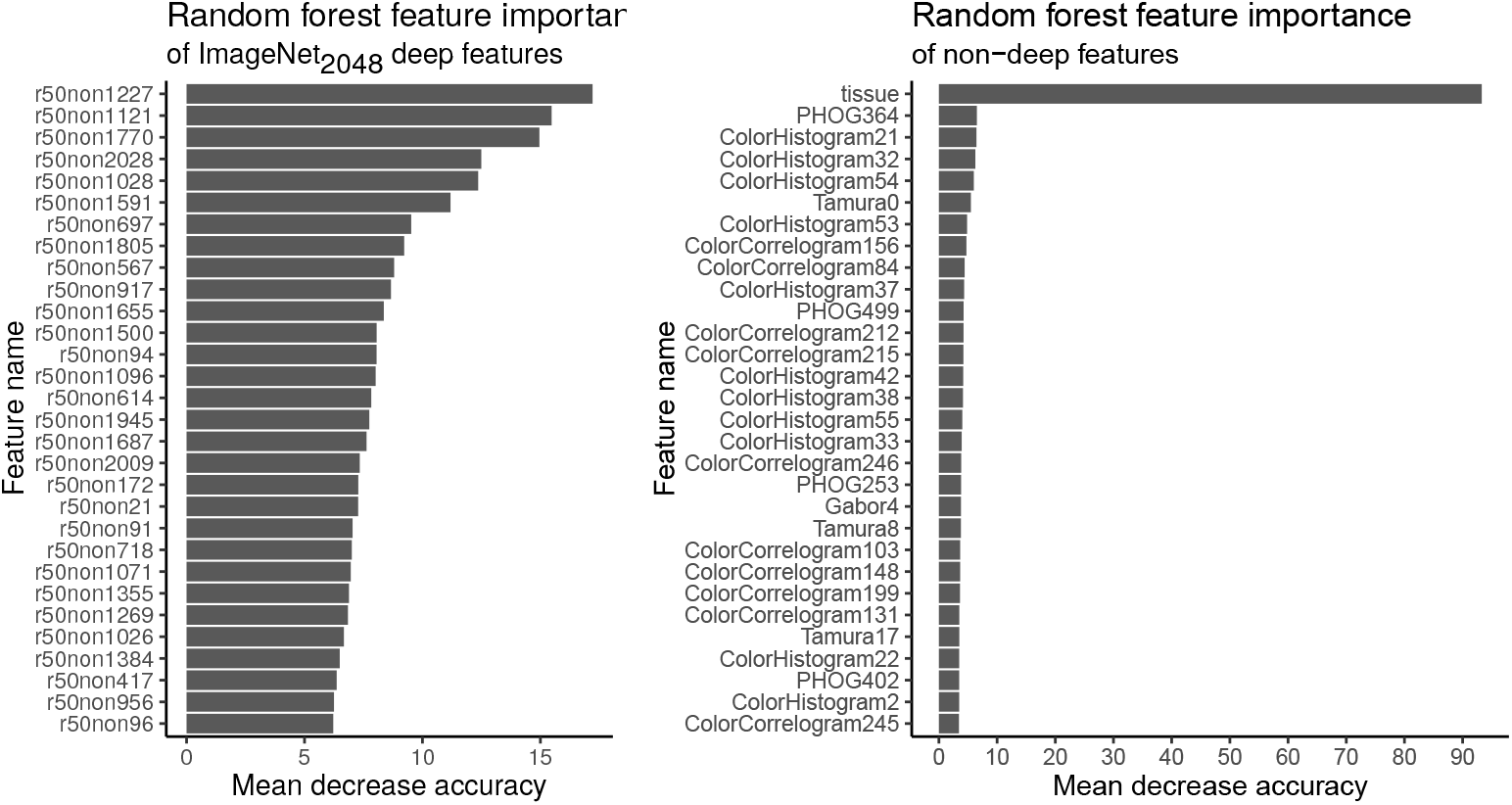
Random Forest feature importance for prioritizing only-natural-image-trained deep features, when non-deep, ImageNet_2048_, and clinical features are used together for learning. As in Fig 4, we use the mean decrease in accuracy to measure Random Forest feature importance. However, we consider here ImageNet_2048_ features, rather than the 100 deep features trained on histopathology images. No visual features here are designed with histopathology in mind or trained on histopathology data. This provides an interpretation of what hand-engineered and only-natural-image-trained deep features are important for disease state prediction, before training the deep neural network on histopathology images and covariates. In this way, we train a Random Forest on 2412 hand-engineered features, ImageNet_2048_ features, and the tissue type covariate. The tissue type covariate is exceedingly important here, highlighting how disease state is reported in our data in a tissue-type-specific manner (Fig 8C1), e.g. infection is more likely reported in lung than breast. Pyramid histogram of oriented gradient (PHOG) features and color features (color histograms and Color Correlograms) are important hand-engineered features complimentary to the ImageNet_2048_ features. PHOG is scale-invariant (due to pyramids) and color-invariant (due to grayscale), while said color histograms and Color Correlograms are rotation-invariant. Taken together, these important hand-engineered features may provide robust pathology representations for Random Forest learning. Such representations may complement ImageNet_2048_ features, with “r50non1227” being the most important of these 2,048 features from the Global Average Pooling layer of a ResNet-50 trained only on the natural images (e.g. cats and dogs) of the ImageNet dataset. Thus of all 2,048 ImageNet_2048_ features, r50non1227 may be prioritized first for interpretation (Fig S11 shows r50non1227 interpretation via heatmaps). *Random Forest feature importance* in the supplement discusses further (Sec S5.10.2).

#### S5.10.2 Random Forest feature importance

We use Random Forest feature importance to infer which features are important for disease state predictions. Hand-engineered visual features, clinical covariates, and deep learning features are concatenated together for a Random Forest to learn to predict disease state, so this single Random Forest classifier provides interpretability of each feature, in context together (Fig 4). We use this to infer broad principles, e.g. important clinico-visual features of disease state are texture (e.g. Local Binary Pattern Pyramid features) and tissue type covariates. Moreover, Random Forest feature importance identified several deep features were more important than the others for disease state prediction, so we focused our analyses on these important deep features.

For a before-and-after-histopathology-image-training comparison, we also consider feature importances when training a Random Forest with ImageNet_2048_ features (Fig S10). ImageNet_2048_ deep features have not been trained on histopathology images or the tissue type covariate. We observe that before histopathology training, these 2048 deep features are complemented by scale-invariant color-invariant edge features (i.e. PHOG) and rotation-invariant color features (color histograms and Color Correlograms). This may suggest disease state prediction benefits from these (i) invariant properties and (ii) features of color/stain intensity and distribution, that are not encoded in the ImageNet_2048_ feature representation. Moreover, the tissue type covariate is strikingly important here (Fig S10). Therefore, after training the ResNet-50 on histopathology images and the tissue type covariate (Fig 3), we find (i) the tissue type covariate importance is reduced presumably because the ResNet-50 has to some extent learned to represent tissue type in its 100-dimensional feature vector, and (ii) scale-invariant and/or color-invariant texture features (e.g. Local Binary Pattern Pyramid [LBPP] and Local Binary Patterns [LBP]) become increasingly important presumably because the ResNet-50 has to some extent learned to represent pathology-relevant edges and color in its 100-dimensional feature vector while texture features are underrepresented. Thus texture features (i.e. LBPP/LBP), are important for disease state prediction, but the deep neural network did not learn similar texture features from the pathology data and learning methods at hand. We likewise reason that LBPP/LBP texture features may have low importance in the context of ImageNet_2048_ features, because ImageNet_2048_ features may represent similar texture, so LBPP/LBP are redundant with ImageNet_2048_ for visual texture features predictive of disease state.

We note several of the most important ImageNet_2048_ features (e.g. r50non1227, r50non1121, …) have an importance measure (i.e. mean decrease in accuracy) greater than the most important hand-engineered features (e.g. PHOG364, ColorHistogram21, …), which may suggest for disease state prediction that ImageNet_2048_ features represent more predictive information than the best hand-engineered features we tested (Fig S10). Alternatively, the high importance of ImageNet_2048_ features may be due to biases in Random Forest learning to choose features that take on many different values [32, 65], as ImageNet_2048_ features do.

In principle, for classification, any interpretable classifier may be used in place of the Random Forest, e.g. logistic regression, support vector machine, or generalized additive model [34]. A careful choice here may demonstrate favorable accuracy and interpretability. We choose a Random Forest as a simple baseline that requires (i) little tuning or preprocessing, (ii) learns interpretable nonlinear relationships among features and covariates, and (iii) provides a measure of similarity for search.

##### Marker mention and SIFT features excluded from Random Forest feature importance analysis

We excluded from our Random Forest feature importance analysis the marker mention covariate and SIFT features, primarily because both did not improve 10-fold cross validation prediction performance when using an ensemble of classifiers, which performed best for prediction (Fig 9: marker 0.8035±0.0043 vs 0.8025±0.0021, *U* = 3, *p* = 0.7; SIFT 0.8035±0.0043 vs 0.8014±0.0022, *U* = 7, *p* = 0.4, two-tailed Wilcoxon rank-sum test). Moreover, SIFT reduces performance when an ensemble is not used (0.7846±0.023 vs 0.7796±0.0019, *U* = 85, *p* = 0.0004114). This may suggest that for sufficiently strong disease state classifiers using H&E images and tissue covariates, the marker mention covariate and SIFT features provide at best only redundant information. For example, if the decision to order a marker test, e.g. IHC, is typically based on the H&E, and a classifier is sufficiently accurate at predicting disease state from H&E, the decision to order a marker test provides no additional disease state information. Secondarily, we excluded the marker mention covariate because it is based on the clinical opinion of all pathologists commenting on this case. Disease state is based on the diagnosis, which is also a clinical opinion. Rather than seeking to explain one opinion in terms of another opinion, we seek to explain opinions in terms of objective information in the H&E or clinicals, e.g. tissue type. We note 10-fold cross validation may provide inflated measures of performance, so for a less inflated examination of the possible contributions of the marker covariate, SIFT features, and deep features, we turn to leave-one-pathologist-out cross validation for search. *Disease state search, first pan-tissue pan-disease method* discusses this (Sec 3.6).

**Fig S11:**
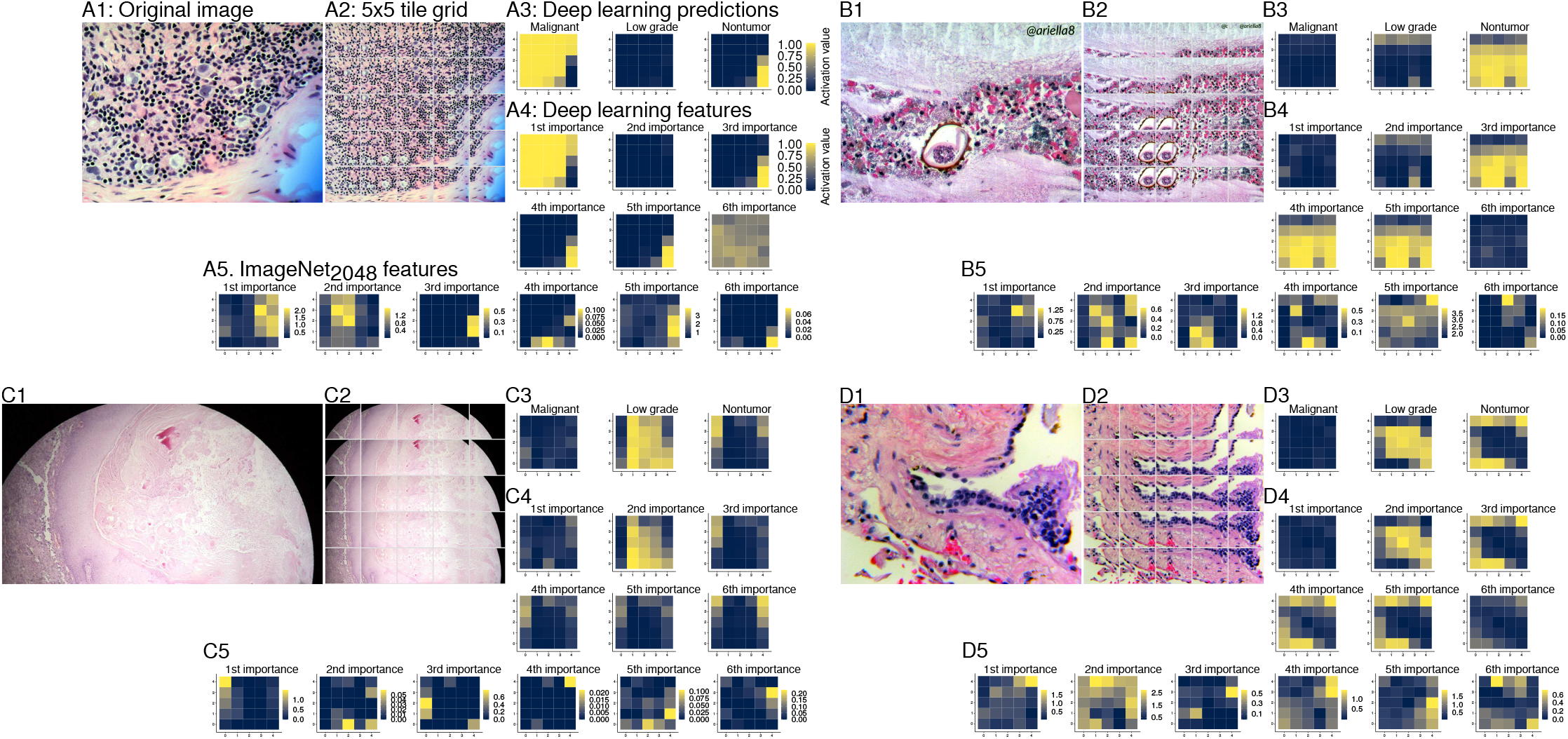
Interpretable spatial distribution of deep learning predictions and features of multiple cases. We compare four cases side-by-side, in the manner of Fig 5. The most important histopathology-trained deep feature is r50_46 (Fig 4 *at left*), which predicts the majority class, e.g. **A3** *left* corresponds to **A4** *upper left.* The second most important feature (r50_30) predicts the second most abundant class, low grade. The third most important feature (r50_85) predicts the third most abundant of three classes, nontumor. The strong class correspondence in important deep features may suggest that removing the top layer’s three class-predictor neurons may incur only a small loss of learned information (Fig 3B *top*). Shown in **A5** *at left* is the most important only-natural-image-trained deep feature (i.e. ImageNet_2048_), “r50non1227” (Fig S10). Continuing left-to-right in **A5** we show ImageNet_2048_ features in decreasing order of importance: r50non1121 (2nd more important), r50non1170 (3rd), r50non2028 (4th), r50non1028 (5th), and r50non1591 (6th). As expected, we find no intelligible pathology-related interpretation of these ImageNet_2048_ features in these heatmaps, because these features are not trained on histopathology data. (**A**) *B.X.:* metastatic lobular carcinoma in satellite lymph node, where malignant activation is high throughout, except the lower right background. (**B**) *C.S.^ĝ^.* juvenile polyp, where nontumor activation is high both for the lentil at lower center (specifically rows 0-2 of columns 1-2 of the 5×5 grid of panel B2, where the lower left corner is row 0 of column 0) and the dark *Ascaris* ova at right (specifically the dark cluster in row 1 of column 4, and evident to some extent in rows 0-2 of columns 3-4), showing the breadth of the nontumor disease state category. (**C**) *R.S.:* is a proliferating epidermoid cyst. Despite viral wart change, we consider this in the low grade disease state (Fig S8). This example also illustrates a different image size and microscope eyepiece field of view artifact. (**D**) *Y.R.:* pulmonary vein lined by enlarged hyperplastic cells, which we consider to be low grade disease state, and **D4** *center top* highlights these low grade cells.

#### S5.10.3 Interpretability of important deep features through activation maps

Deep neural networks have a restricted field of view, but this is an advantage for interpretability, because one can systematically sweep a trained neural network across an image to localize deep feature activations. At each location in the sweep, a 224 ×224px image patch is fed to the trained neural network for interpretation. However, our Random Forest uses a set representation of the deep features, formed as a sum of deep feature vectors systematically sampled throughout the overall image of arbitrary size (Fig 1). Therefore, for spatial localization of disease state we make heatmaps to depict the deep feature activations at those sampled locations. These heatmaps indicate a correspondence between the most important deep features and the class labels. This approach facilitates deductive reasoning about predictions, e.g. (i) in Fig 5 the image overall is predicted by the deep-learning-random-forest hybrid classifier to be low grade (not shown), (ii) this classifier includes deep features (Fig 1C), (iii) a deep feature of ours is by definition a vector sum of images (Fig 1C) shown in the grid (Fig 5D2), (iv) the second most important deep feature (r50_30 in Fig 4) is known to correspond to low grade (e.g. Fig 5C2), (v) r50_30 is active with a value of more than 0.5 for images shown in the grid center (Fig 5D2), (vi) therefore the classifier predicts low grade partly because images near the grid center have the low grade feature. Similar to our work, previous work used Random Forest feature importance for feature selection on a pretrained deep neural network [67], though to the best of our knowledge we are the first to use Random Forest feature importance for feature selection on a deep neural network retrained on the same task as the Random Forest.

**Fig S12:**
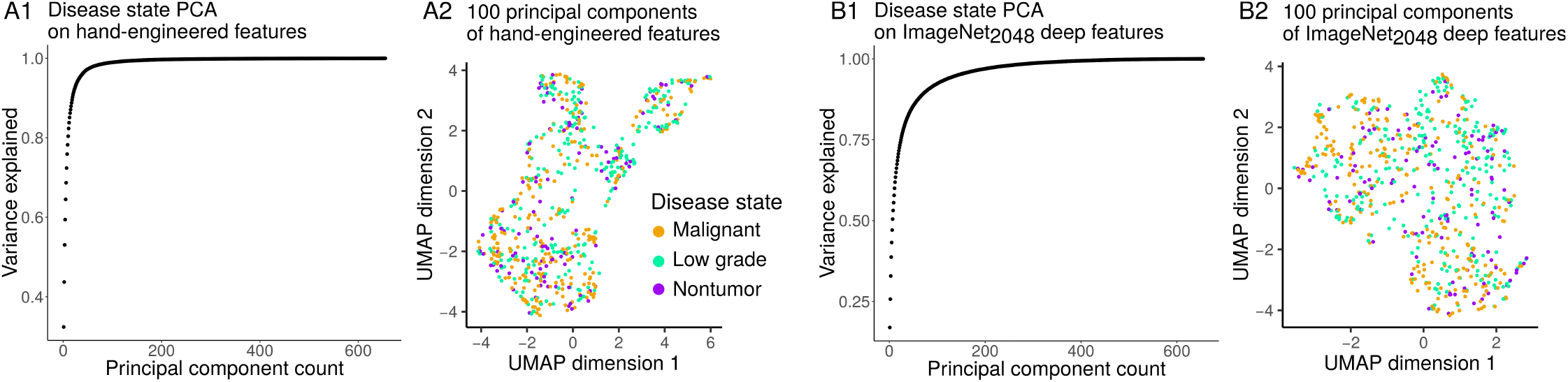
Disease state clusters after dimensionality reduction. As in Fig 6 we apply the UMAP [30] algorithm to determine if there are clusters of patient cases that have meaningful groups of features for the prediction of disease state. However, to investigate if cluster quality can be improved through dimensionality reduction, we first apply principal components analysis (PCA) to reduce hand-engineered feature dimensionality from 2,412 to 100 principal components, and follow the same procedure for ImageNet_2048_ features. In practice, PCA is a common preprocessing step for the t-SNE clustering algorithm [66], but UMAP claims to have no computational restrictions on input dimension (so PCA is not expected to be required for UMAP) [30]. (**A1**) We show that 100 principal components explain 98.98% of the variance of the 2,412 hand-engineered features. Our histopathology-trained deep features are similarly 100 dimensions (Figs 3C, 6C1). (**A2**) As expected, PCA preprocessing does not qualitatively change UMAP clusters based on hand-engineered features. (**B1**) We show that 100 principal components explain 92.35% of the variance of the only-natural-image-trained ImageNet_2048_ deep features. (**B2**) As expected, PCA preprocessing does not qualitatively change UMAP clusters based on ImageNet_2048_ deep features. We conclude the vague clusters from hand-engineered features or lack of clusters from ImageNet_2048_ is not a UMAP-related artifact of their high dimensionality, but instead simply means these features do not clearly group patients by disease state.

### S5.11 Machine learning methods discussion

Because our image feature vectors are so wide, e.g. 2,412 dimensions (Fig S9) or more, we found best results with Random Forests when the number of features to consider for a decision/split was half (rounded up) of the total attribute count. This was especially important for covariates, e.g. the tissue covariate for disease state prediction. For search using Random Forest similarity, we grew each tree to a maximum depth of 10.

#### S5.11.1 Deep learning

##### Cross entropy loss for learning

We optimize the deep neural network with an unweighted cross entropy loss (Eqn 1, where *I_y_i_∈C_i__* is the indicator function being 1 when the example *x* class label *y_i_* is class *C_i_*, and 0 otherwise)^7^, for minibatches of size *N* = 64 224×224px images and typically *C* = 3 classes (nontumor, low grade, malignant). We use stratified bootstrap sampling for each epoch, so all classes have equal weight.

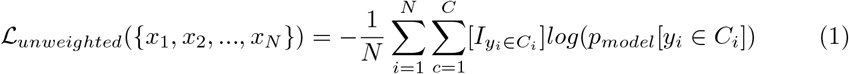

**Fig S13:**
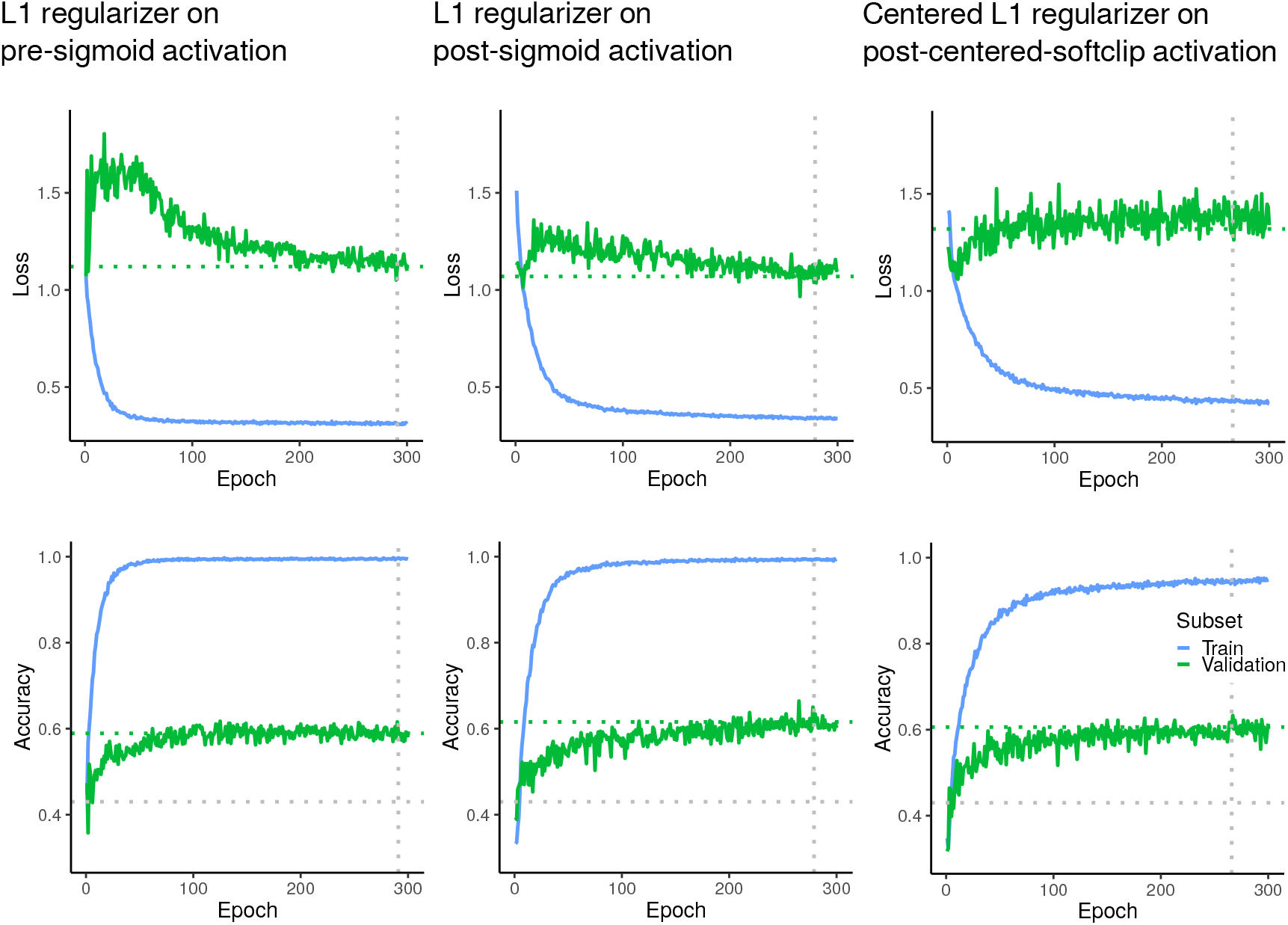
Deep learning performance curves. Loss and accuracy for training and validation sets, for one fold of 10-fold cross validation, using a ResNet-50 to predict disease state, for individual 224×224px image patches. Validation accuracy improves with training, though high validation accuracy may suggest (i) the ground truth label may not apply universally throughout the image which is larger than 512×512px, (ii) samples whose state may not fit with our three-class disease state schema, (iii) mislabeled samples, (iv) deep learning overfit. Indeed, we see in many of our images mixtures of diseases and disease-free tissue, so (i) may be especially likely, and more advanced methods, such as multiple instance learning, may overcome this as we acquire more data. Observing that validation accuracy improved with training, we proceeded with this simple supervised deep learning approach as a proof-of-principle.

##### ResNet-50 learning

For our deep learning, we freeze no layers of the ResNet-50. We train end-to-end with learning rate of 0.01 and Nesterov accelerated gradient momentum of 0.9 [68–70]. We use Keras’ learning rate decay of 10^-5^, which reduces the learning rate each batch. Moreover, we follow a learning rate schedule, where learning rate is divided by 100 until the end of epoch 1, but by 10 until the end of epoch 3. We train with Mixup [29], mixing according to draws from a beta distribution with *alpha* =1.4 and *beta* = 0.4. Our data augmentation is random flips, random free rotations, grayscale Gaussian noise in RGB color space (mean=0, stdev=0.5), and random brightness adjustment (uniform distribution, −0.05 to 0.05). We white-balance images before processing.

Freezing layers lowered validation accuracy. Alternative architectures such as DenseNet, Inception, and Xception trained more slowly and lowered validation accuracy. We do not report these results.

#### S5.11.2 Deep set learning feature interpretation

As our ResNet-50 deep neural network trains (Fig S13), a 100-dimensional feature representation is learned (Fig S14), by the 100-neuron layer we append to the ResNet-50 (Fig 3C). A spatially-localized empirical interpretation of deep learning predictions and feature activations is available in Figure 5. We analytically interpret our deep features through their activation functions, regularization, and relationship with set scalar cardinality, below.

**Fig S14:**
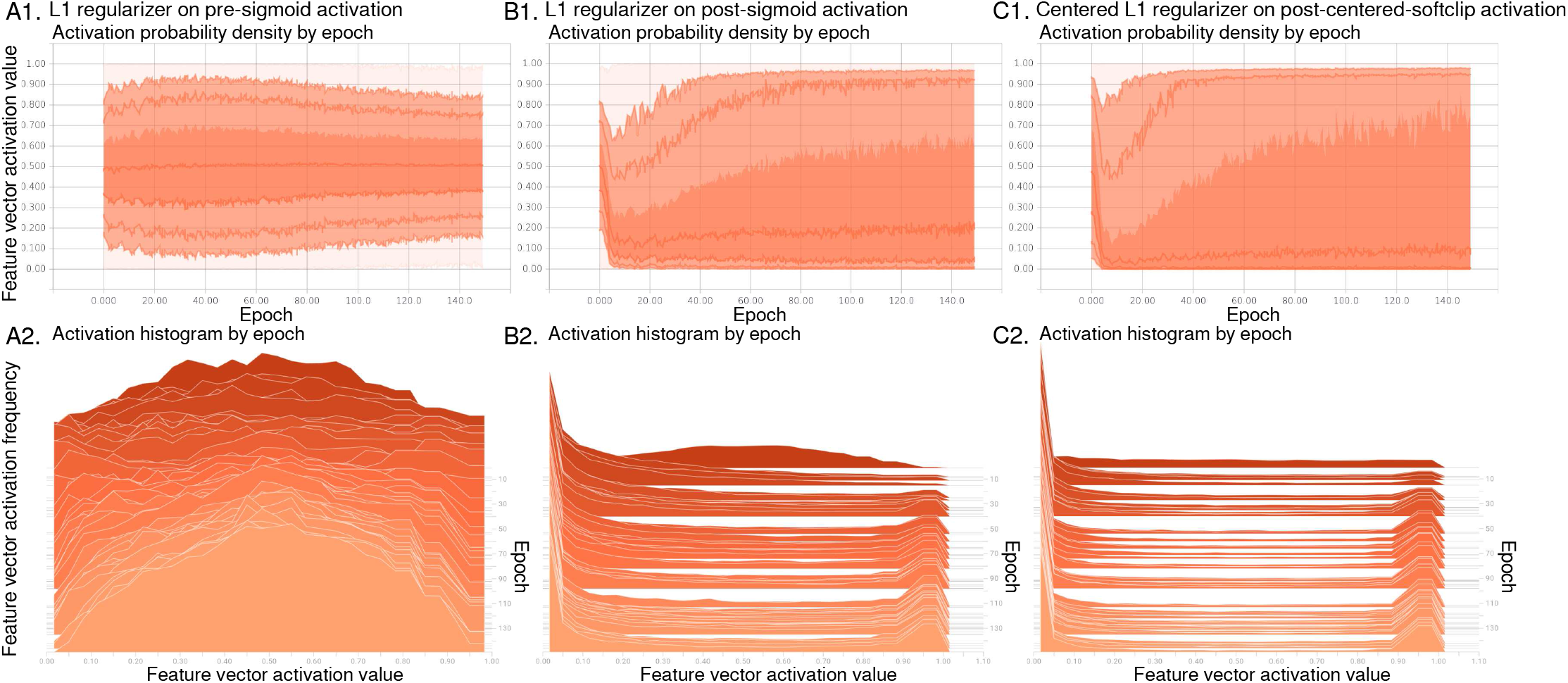
Deep features. We appended a 100 neuron fully connected layer on top of a ResNet-50 to learn a concise 100-dimensional “bottleneck” feature vector representation of a 224×224px image patch (Fig 3). This feature vector takes different distributions, depending on regularization. The top row shows a 100-dimensional activation distribution as standard deviations from the mean, and training proceeds left to right. The bottom row shows the same activation distribution as the top row, but as a histogram, with training proceeding from dark red at the top to orange at the bottom. (A) Feature values are centered at 0.5 when L1 regularization on the activation value before a sigmoid squashing function is applied, because a sigmoid transforms a 0 value to 0.5. (B) Feature values are approximately bimodal when L1 regularization on the activation value after sigmoid squashing is applied, because L1 regularization enforces sparsity by penalizing non-zero values. (C) Feature values are more strongly bimodal when our binarizing regularization (a.k.a. centered L1 regularization, Fig S15C3,D) on the activation value after our centered soft clipping activation function is applied, because this regularization penalizes values near 0.5 and centered soft clipping saturates to 0 or 1 more quickly than a sigmoid. We ultimately chose our centered L1 regularization and centered soft clipping activation to represent the deep feature space for disease state prediction, because (i) it performed better with Random Forest learning over feature vector sums of a set of multiple images (Fig S15A), (ii) it demonstrated comparable validation accuracy in deep learning (Fig S13), (iii) Random Forests have a bias to select features that take many values [32,65] so approximately binary deep features may reduce this bias by having a restricted distribution of values, and (iv) we believe approximately binary deep features to be simple and interpretable as set scalar cardinality, i.e. counters (Fig 5 and Eqn 8).

##### Centered soft clipping activation function definition

To learn sharply binary features from the deep learning, we define a steep activation function, called centered soft clipping (CSC) (Fig S15B and Eqn 3). This is derived from soft clipping (SC) [71], which is not centered at x=0 (Fig S15B and Eqn 2). Like centered soft clipping, the sigmoid and hyperbolic tangent activation functions centered at x=0, but they are not as steep. There is a steepness parameter (p) in [centered] soft clipping, and we let p = 2 for our experiments (Eqns 2, 3).

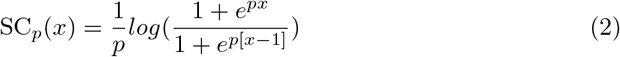

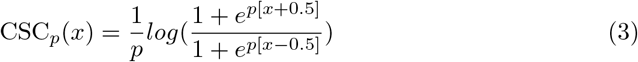

##### Centered soft clipping in the limit as Heaviside step function

Furthermore, we note in the limit *p* → ∞, centered soft clipping converges to the Heaviside step function *H*(*x*), a type of indicator function which is 1 for positive values, 0 for negative

**Fig S15:**
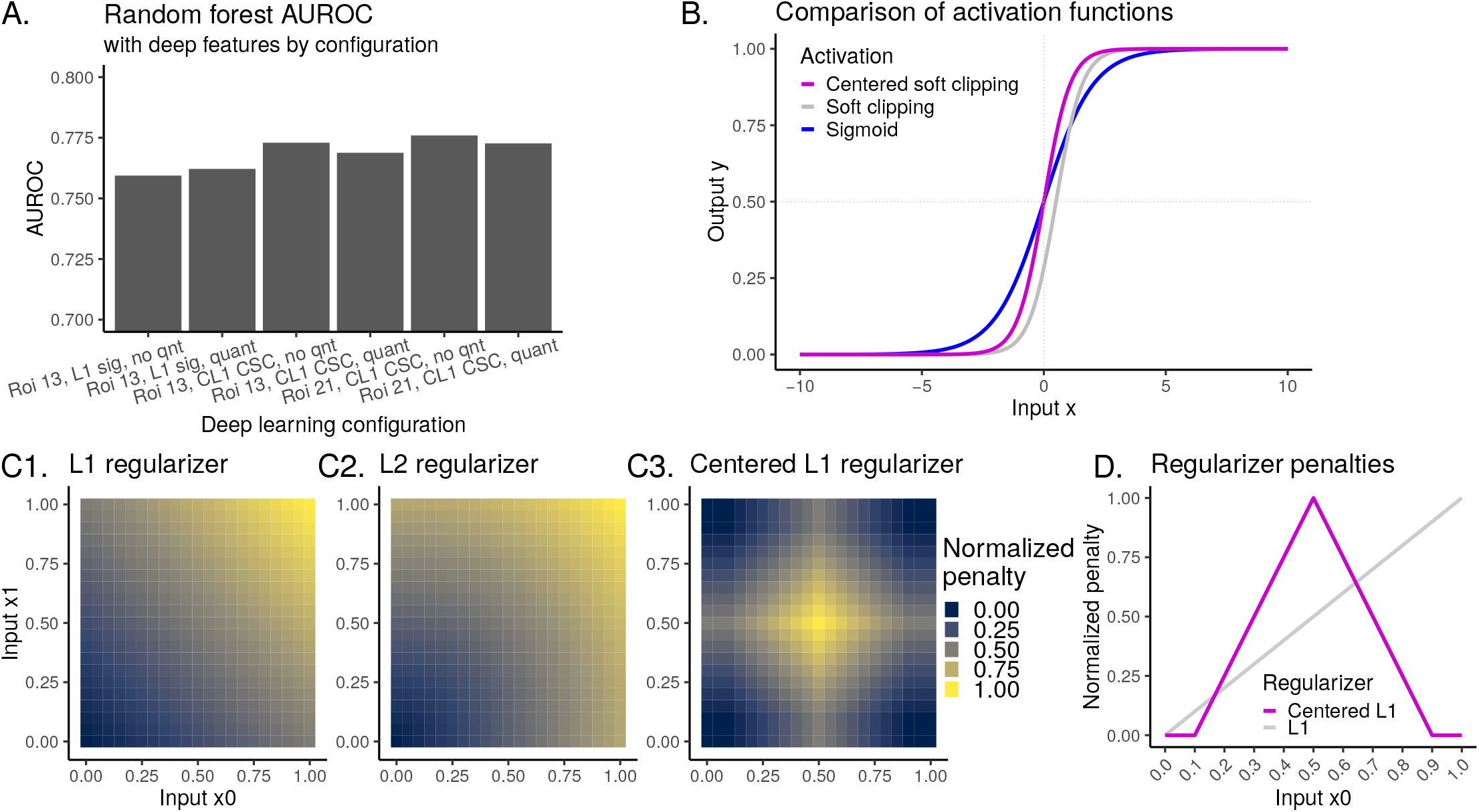
Sigmoid vs centered soft clipping performance. (**A**) Random Forest AUROC when ResNet-50 trained with sigmoid activation function with L1 regularization, versus centered soft clipping activation function with centered L1 regularization. Centered soft clipping demonstrates higher performance. Increasing the number of regions of interest (ROI) images in a deep set from 13 to 21 (Fig 1C shows 21) improves performance slightly. Quantizing features to either 0 or 1 slightly reduces performance when using centered soft clipping. (**B**) Comparison of sigmoid activation function with soft clipping [71] (Eqn 2) and our centered soft clipping (Eqn 3). Our centered soft clipping is centered at x=0 and steeper than sigmoid, which we argue is amenable for learning interpretable (Fig 5) and sharply binary hash codes. (**C**) Comparison of L1 (Eqn 11), L2, and our centered L1 (Eqn 13) regularizers in two dimensions, *x_0_* and *x*_1_. Our L1 regularizer penalizes values close to 0.5, to encourage a binarized feature representation. (**D**) Comparison of L1 and our centered L1 regularizers, in one dimension, for clearer depiction of our centered L1 regularizer’s margin *m* = 0.1 (Eqn 13), for stable learning and vanishing gradient avoidance.

values, and 0.5 for zero values (Eqn 6). In the limit *p* → ∞, CSC_*p*_(*x*) is a binary indicator of the presence or absence of a feature in a 224×224px image patch (Fig 1B), representing binary logic. Smaller values, e.g. *p* =2, allow the representation of a small amount of probabilistic uncertainty regarding this presence or absence, where this area of uncertainty is infinitesimally small for *p* → ∞.

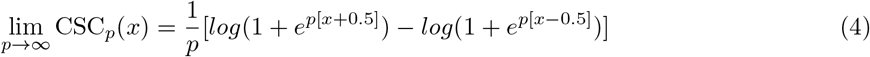

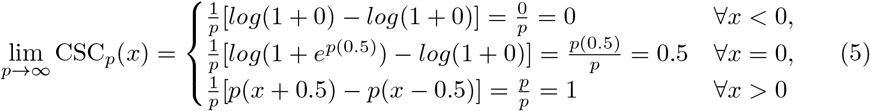

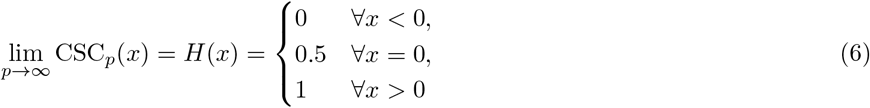

##### Deep feature as set scalar cardinality

To our Random Forest, a deep feature is the scalar cardinality (Eqns 7–8) of the set of these presences measured at 21 different locations throughout the original image (Fig 1C). Here, *x* represents one of the 100 features from the trained layer atop the ResNet-50 in Figure 1B, and *I* is an indicator function of set membership.

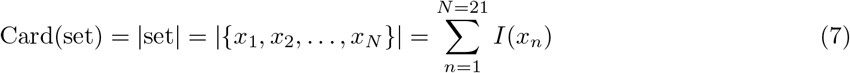

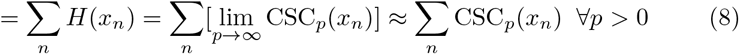

##### Data as Bernoulli process, deep features, and expected value

To consider some input data passed through the ResNet-50, we model a random set as a Bernoulli process, with each set member taking some value on the interval (0, ∞) at probability φ, otherwise some value on the interval (-∞, 0). Let 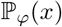 be a Bernoulli trial that is (0, ∞) at probability *φ*, otherwise (-∞, 0) A Heaviside step function (Eqn 6), which our centering soft clipping (CSC) activation function approximates in the limit of *p* → ∞, transforms this 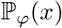 to a standard Bernoulli trial taking value 1 at probability *φ*, otherwise 0. Thus the cardinality of this random set is proportional to the Bernoulli trial’s expected value 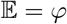 (Eqn 10). In a sense, for a fixed set size e.g. *N* = 21, the Random Forest may learn what thresholds of 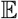 of these deep features are predictive of the classes of interest, in the context of the other available deep and non-deep features (Fig 1C).

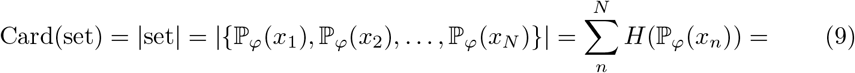

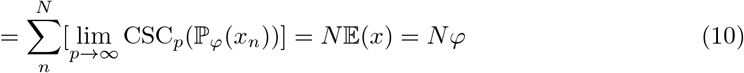

##### Centered L1 regularizer in the context of Gaussian initialization

Neural network weights are often initialized with Gaussian noise having mean of zero [68], so centering the activation function at zero removes initial bias towards learning a 0 or 1 when combined with our centered L1 regularizer (Eqn 3). Whereas a standard L1 regularizer (Eqn 11) penalizes all non-zero values, our centered L1 regularizer penalizes values close to 0.5 (Eqn 13), with the assumption that all values are between 0 and 1, which is true for values from centered soft clipping (Eqn 3).

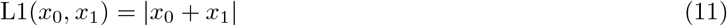

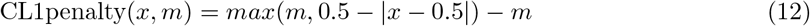

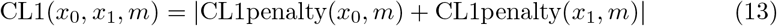

To approximately binarize deep features, our centered L1 regularizer penalizes output values close to 0.5, which occur when *x* = 0 with sigmoid and centered soft clipping activations. (Fig S15D). For stable learning, and to help avoid the vanishing gradient problem [72,73] from values infinitesimally close to 0 or 1, our centered L1 regularizer has no penalty for values less than 0.1 or greater than 0.9. Restated, we let the L1 regularizer’s margin parameter *m* ∈ [0, 0.5] be *m* = 0.1.

##### Binary hash code learning

Lin *et al* [74]. use deep learning with sigmoid activations to learn binary hash codes for search of clothing images. We intend to learn sharply binary hash codes with centered soft clipping and centered L1 regularization together, for interpretability, pathology search, and set representation learning. We find centered soft clipping performed slightly better for classification (Fig S15A) and provided a small but significant increase in search performance (Fig 10B).

### S5.12 Disease state search distance function and performance

Pathology search performance is shown in Figure 10B, and detailed in Table S1. We found a combined distance measure (Eqn 14) worked best for search (i.e. Table S1 top row). The combined distance between two examples (i.e. Dist(*x*_0_, *x*_1_)) is the number of trees in the Random Forest (i.e. 1000), minus the Random Forest similarity of the covariates and non-deep features (i.e. Sim_RF_(*x*_0_, *x*_1_)), plus five times (i.e. *α*_SIFT_ = 5) the L1 norm of the 5 largest SIFT cluster medoids vector sum (i.e. L1_SIFT_k5__ (*x*_0_,*x*_1_)), plus the L1 norm of the top three deep features from each example (i.e. L1_Deep3_(*x*_0_, *x*_1_)). For Deep_3_, we use the top three most class-associated features of example *x*, one such feature for each of the three classes (nontumor, low grade, or malignant per Fig 5C). We initially found these three features using Random Forest similarity (Fig 4), but subsequently identify these features by minimizing the feature’s error with respect to a class label in the training data, for computational expediency.

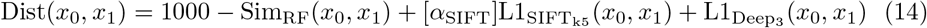

### S5.13 Supplementary Disease state search results

#### S5.13.1 Clinical covariates improve search performance

We ask if pathology-specific clinical covariates improve search performance. Including the tissue type covariate significantly improves performance compared to not using this covariate (0.5640±0.0024 vs 0.6533±0.0025, *U* = 100, *p* = 0.0001796). We find including the marker mention covariate significantly improves performance further (0.6533±0.0025 vs 0.6908±0.0021, *U* = 100, *p* = 0.0001796). Therefore including pathology-specific clinical covariates for pathology search is currently justified, because these improve search performance. We reason that disease states are reported at tissue-type-specific rates, and cases that mention marker tests (e.g. IHC) tend to be more similar to other cases that mention marker tests. Often, marker tests are used to subtype malignancies (e.g. TTF-1), but this is not always the case (e.g. Ki-67). We believe this is the first multimodal classifier that demonstrates improved search performance when combining pathology imaging features with clinical covariates (i.e. tissue type and mention of marker tests) that may be missing for some patients (Fig 10B).

#### S5.13.2 In the context of other features, nuclear features of disease are better represented by the most prevalent SIFT clusters rather than all SIFT

##### Used alone, all SIFT is better than chance, but does not complement other features

We ask if cell nuclei features, as represented by SIFT, may represent disease state, and if so, which SIFT representations perform better than others for pathology search. Inspired by continuous bag-of-words methods that represent a context as a vector sum of words [64] and bag-of-visual-words methods that leverage SIFT [75], our simplest baseline detects all SIFT interest points in an image, then takes the sum of all these SIFT feature vectors to represent an image overall. This way, an image is represented by a 128-dimensional SIFT set representation, where the set cardinality is equal to the number of SIFT interest points in the image, which varies among images. Because SIFT features are non-negative and cover nuclei, the magnitude of this vector may increase as nuclear density increases. This performs better than chance when used alone (0.4636±0.0024 vs 0.3967±0.0044, *U* = 100, *p* = 1.083e-05), but reduces performance when used in the context of our 2,412 hand-engineered features, tissue covariate, and marker covariate (0.6908±0.0021 vs 0.5728±0.0022, *U* = 100, *p* = 0.0001796). We conclude that naïvely using SIFT features this way to estimate nuclear density allows limited discrimination of disease state when used alone, but does not complement other features when used together for pathology search.

**Table S1:**
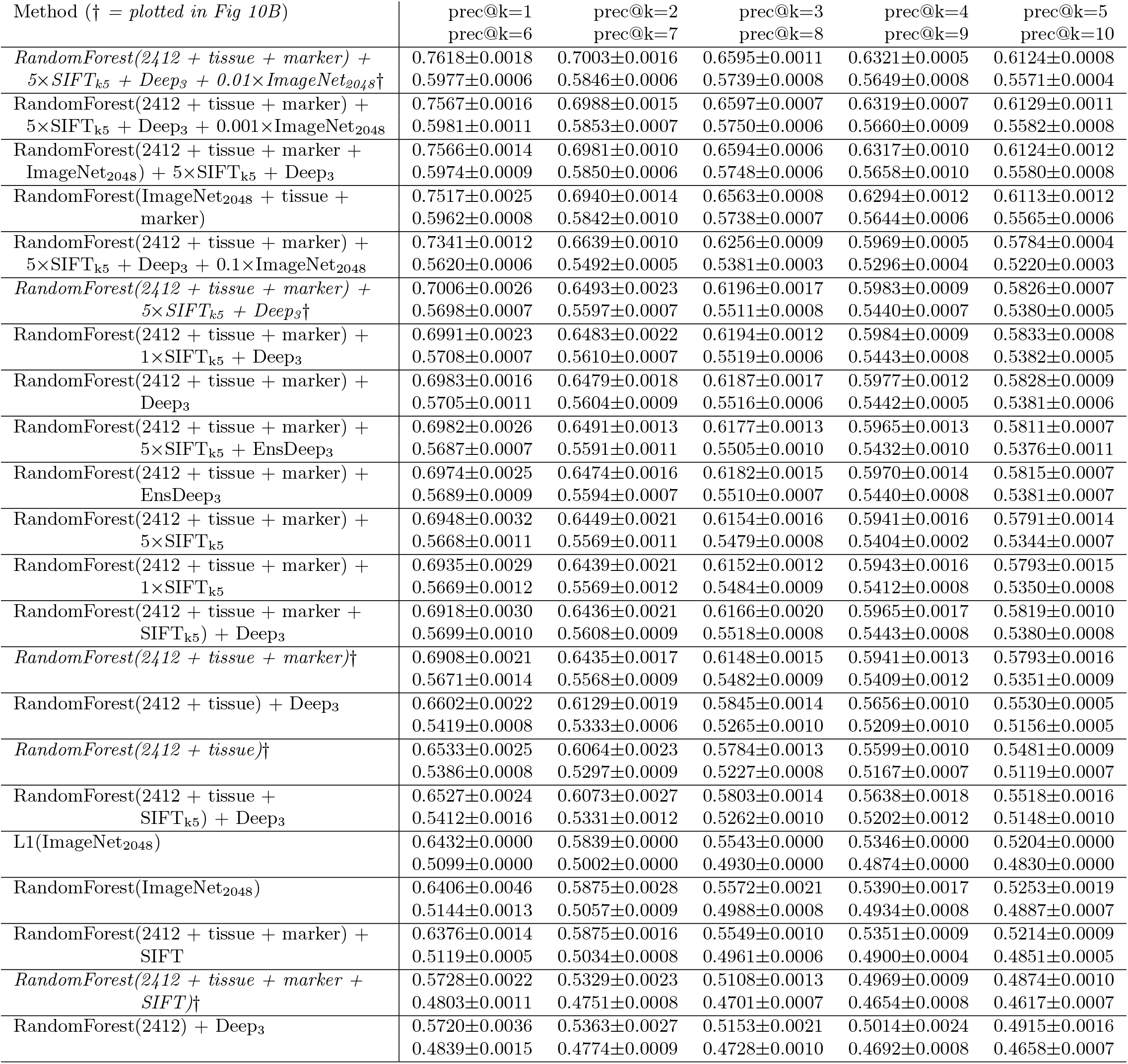
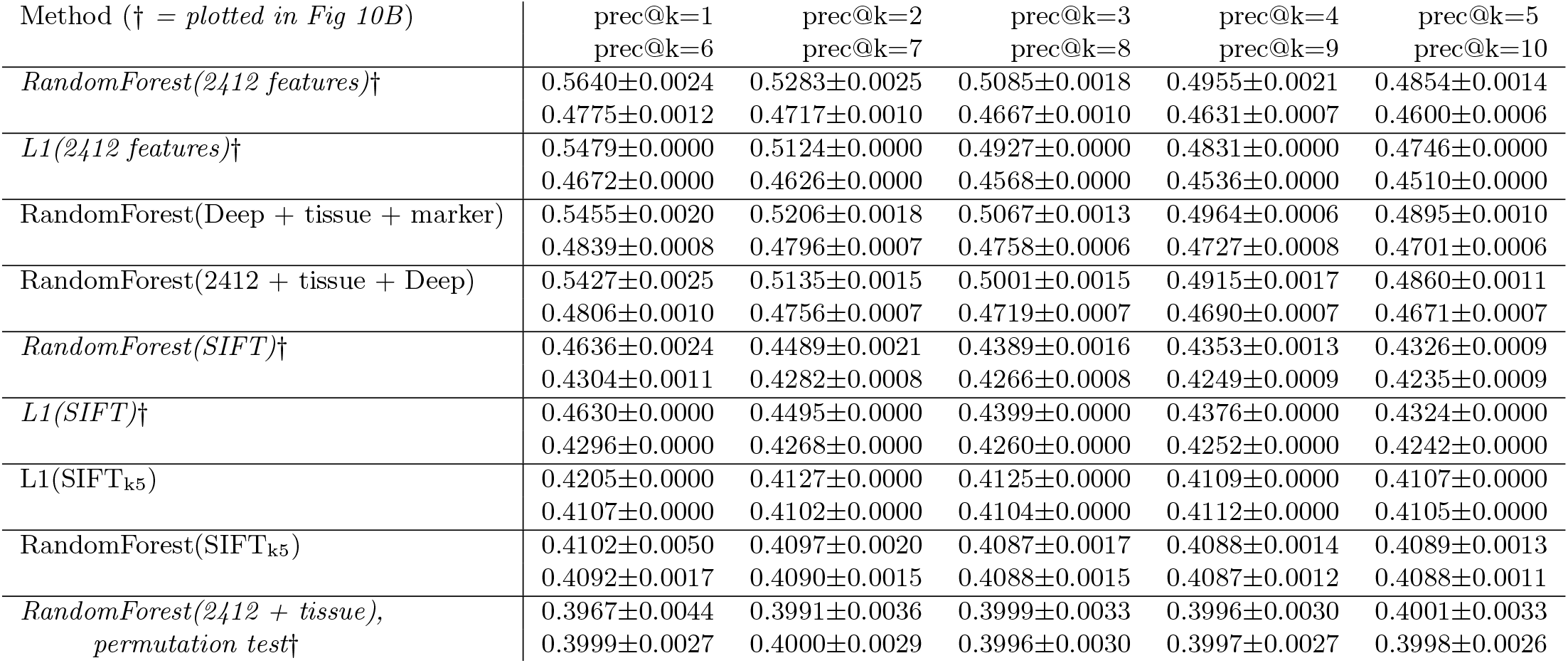
Case similarity search performance in detail. Precision@k for search as shown in Figure 10B, with additional experiments. To estimate 95% confidence intervals, recall standard error of the mean is the sample standard deviation (i.e. stdev) divided by 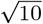, since we perform ten replicates. L1(…) indicates distance is the L1 norm of the feature vector. RandomForest(…) indicates distance is the number of trees in the forest (i.e. 1000) minus Random Forest similarity calculated on the feature vector. Tissue and/or marker covariate use are indicated by “tissue” and/or “marker”, respectively. “2412”/”2412 features” indicates our 2,412 hand-engineered features are used (Fig S9). However, “ImageNet_2048_” indicates the 2,048 top-level features are used from a ResNet-50 trained only on ImageNet images. No histology images are used to train this ResNet-50’s 2,048 features, but these features are summed over 21 locations, just as the ‘histology-image-trained ‘Deep” features are (Fig 1C). “Deep” indicates the full 100-dimensional feature vector is used (Fig 1B,C). “Deep_3_” indicates only the top 3 class-correlated features are used (Fig 5, see Eqn 14 for combining distances). “EnsDeep_3_” indicates the top 3 class-correlated features are used, averaged across three neural networks in an ensemble. “SIFT” indicates all SIFT interest points are summed to represent an image. “SIFT_k5_” indicates the 5 largest of 25 medoids are summed to represent an image. “5×SIFT_k5_” indicates the SIFT_k5_ vector is multiplied by five (i.e. α_SIFT__k5_ = 5), which changes L1-based distances (Eqn 14). We find best performance for “RandomForest(2412 + tissue + marker) + 5×SIFT_k5_ + Deep_3_”, which is 1000 minus Random Forest similarity, plus five times L1(SIFT_k5_), plus L1(Deep_3_).

**Table S2:** Case similarity search performance per-pathologist. Precision@k per-pathologist, corresponding to Fig 10C *at right,* for the best performing search method *RandomForest(2412 + tissue + marker) + 5×SIFT_k5_ + Deep_3_ + 0.01×ImageNet_2048_*, c.f. Table S1. Leave-one-pathologist-out cross validation is repeated ten times. We considered 24 of 25 pathologists for this analysis, excluding one pathologist who shared no human acceptable H&E images for disease state prediction (Fig S4 details acceptability criteria). Each included pathologist’s performance is averaged over these ten repetitions. This average performance is plotted in Fig 10C, where the 24 lines correspond to 24 pathologists, and error bars are standard error for these ten replicates. For precision@k=1 mean, each of the 24 pathologists’ ten-repetition average precision@k=1 is averaged and reported. In this way, the mean prec@k=1 (i.e. 0.7470) is what precision@k=1 may be expected for a pathologist whose data we have not seen before, when averaged over many images for that pathologist. Bootstrapped confidence intervals (CI) for median and mean are shown for 10,000 replicates. There is not a significant difference between the means and medians, so any differences may be due to chance alone.

|  | median | median 95% CI | mean | stdev | mean 95% CI |
| --- | --- | --- | --- | --- | --- |
| prec@k=1 | 0.7819 | 0.7499-0.8065 | 0.7470 | 0.1167 | 0.6978-0.7896 |
| prec@k=2 | 0.7237 | 0.6643-0.7482 | 0.6941 | 0.0967 | 0.6550-0.7300 |
| prec@k=3 | 0.6766 | 0.6320-0.6938 | 0.6523 | 0.1030 | 0.6115-0.6922 |
| prec@k=4 | 0.6373 | 0.6000-0.6769 | 0.6304 | 0.0975 | 0.5920-0.6676 |
| prec@k=5 | 0.6112 | 0.5771-0.6563 | 0.6123 | 0.0962 | 0.5742-0.6501 |
| prec@k=6 | 0.5982 | 0.5401-0.6451 | 0.5984 | 0.0931 | 0.5622-0.6345 |
| prec@k=7 | 0.5858 | 0.5236-0.6378 | 0.5847 | 0.0907 | 0.5494-0.6203 |
| prec@k=8 | 0.5736 | 0.5117-0.6301 | 0.5741 | 0.0879 | 0.5399-0.6079 |
| prec@k=9 | 0.5620 | 0.5042-0.6230 | 0.5644 | 0.0886 | 0.5296-0.6003 |
| prec@k=10 | 0.5526 | 0.4964-0.6132 | 0.5558 | 0.0884 | 0.5210-0.5899 |

##### SIFT learning challenging, but a combined distance function of L1 norm and Random Forest similarity improves search

In light of the aforementioned reduced performance with SIFT, we ask if Random Forest similarity contributes to this performance reduction. We observe the SIFT L1 norm and SIFT Random Forest similarity performance differences may be due to chance alone (0.4630±0.0000 vs 0.4636±0.0024, *U* = 40, *p* = 0.4429). The Random Forest does not appear to learn from this SIFT representation. We additionally observe search performance is reduced less when adding the SIFT L1 norm to Random Forest similarity as a combined distance measure (Eqn 14), rather than training the Random Forest on SIFT features (0.5728±0.0022 vs 0.6376±0.0014, *U* = 0, *p* = 0.0001796). This may suggest the Random Forest overfits or cannot learn from these continuous bag-of-words SIFT features, but this may be mitigated by a combined distance function that does not rely on Random Forest learning from SIFT features (Eqn 14).

##### SIFT clusters provide complimentary information for search

We then ask if SIFT may provide complimentary information for search, when using SIFT to estimate nuclear shape and edge distributions, rather than nuclear density. For this approach, we (i) detect all SIFT interest points in an image, (ii) form 25 clusters using k-medoids++ clustering with the L1 norm of the SIFT feature vectors for all interest points, (iii) retain the 5 medoids corresponding to the 5 most abundant clusters, and (iv) take the sum of the medoid SIFT features to represent the image overall. This way, an image is represented by a 128-dimensional SIFT set representation, where the set cardinality is always 5, the number of retained medoids. We call this a SIFT_k5_ set representation, for k=5 medoids from k-medoids++ clustering. Because SIFT features represent shapes and edges, and because we use only the most abundant cluster medoids, this vector may represent the prevailing shapes and edges of nuclei in the image overall. This performs better than chance when used alone (0.4102±0.0050 vs 0.3967±0.0044, *U* = 100, *p* = 0.0001817), and significantly improves performance when used in the context of 2,412 features, tissue covariate, and marker covariate (0.6908±0.0021 vs 0.6935±0.0029, *U* = 19.5, *p* = 0.02308, though the effect is small.

##### Combined distance function coefficient allows SIFT prioritization, but careful calibration not needed

Because SIFT feature vector magnitude is small, and the number of medoids we retain is small (i.e. 5), we ask if search would benefit from a SIFT coefficient greater than one in the combined distance function (i.e. α_SIFT_ in Eqn 14), to increase the relative contribution of SIFT features to search. We let α_SIFT_ = 5. When combined with deep features, which we discuss later, we report a very small but significant improvement in search performance when α_SIFT_ = 5 (0.6983±0.0016 vs 0.7006±0.0026, *U* = 21.5, *p* = 0.03423) but not when α_SIFT_ = 1 (0.6983±0.0016 vs 0.6991±0.0023, *U* = 34, *p* = 0.2408). However, search performance is not significantly different depending on the choice of α_SIFT_ here (0.6991±0.0023 vs 0. 7006±0.0026, *U* = 30, *p* = 0.14). Therefore, we do not observe compelling evidence in favor of selecting α_SIFT_ carefully, and caution that careful selection risks overfit. We note αSIFT = 5 may be viewed as a micro-optimization, but this effect is very small. It appears this effect is carried by SIFT_k5_ rather than α_SIFT_, so we conclude it may be better to focus on feature engineering or learning, rather than coefficient selection. We report this to show α selection in the combined distance function has only a slight effect.

#### S5.13.3 Deep features synergize with other features, informing search more than nuclear SIFT features, but less than clinical covariates

##### For deep features, a combined distance function of L1 norm and Random Forest similarity also improve search

Because search performance improves when considering SIFT_k5_ set features in a combined distance function, we ask if search performance improves when considering deep set features in a combined distance function (Eqn 14). We observe that naïvely concatenating the full 100-dimensional deep set representation with other features for Random Forest learning reduces performance (0.6533±0.0025 vs 0.5427±0.0025, *U* = 100, *p* = 0.0001796). We then select the interpretable top 3 most class-associated deep features (Figs 4, 5D1-3) and observe using these in a combined distance function improves search performance (0.6533±0.0025 vs 0. 6602±0.0022, *U* = 4, *p* = 0.0005773). Performance remains improved when including the marker mention covariate (0.6908±0.0021 vs 0.6983±0.0016, *U* = 0, *p* = 0.0001817). Given that both SIFT set features and deep set features both perform better in a combined distance function rather than within Random Forest similarity, we suspect our Random Forest parameters facilitate sensitive learning from few covariates, but may be oversensitive to a large number of set features that each take many different values. *Machine learning methods discussion* in the supplement discusses (Sec S5.11). We conclude an interpretable reduced representation of deep set features improves search performance when considered in a combined distance function, though this effect is small. We expect the effect to increase with more data, because deep learning can refine feature representations in a scalable data-driven manner, and because more advanced deep learning methods may be possible with more data.

##### Deep features from supervised learning inform search more than nuclear SIFT features

Given that both SIFT_k5_ and Deep_3_ features improve search performance, we ask if pathology-specific Deep_3_ features improve search performance more than pathology-agnostic SIFT_k5_ features. Indeed, we observe Deep_3_ features improve search performance significantly (i) more than SIFT features (0.6983±0.0016 vs 0. 6376±0.0014, *U* = 100, *p* = 0.0001817), (ii) more than 1×SIFT_k5_ features (0.6983±0.0016 vs 0.6935±0.0029, *U* = 94, *p* = 0.0003248), and (iii) more than 5×SIFT_k5_ features (0.6983±0.0016 vs 0.6948±0.0032, *U* = 83.5, *p* = 0.01251). We conclude learned pathology-specific deep features inform pathology search more than hand-engineered pathology agnostic SIFT features, though SIFT may cover nuclei.

##### Deep features and SIFT features are complementary

To determine if SIFT and deep features represent non-overlapping concepts in pathology, we ask if combining SIFTk5 and Deep3 features improves performance, compared to using either one, in the context of other features. We observe Deep3 features significantly improve performance when considering 5×SIFT_k5_ and all other features (0.6983±0.0016 vs 0.7006±0.0026, *U* = 21.5, *p* = 0.03423). We likewise observe 5×SIFT_k5_ features significantly improve performance when considering Deep_3_ and all other features (0.6948±0.0032 vs 0. 7006±0.0026, *U* = 7, *p* = 0.0004871). These small effects suggest SIFT_k5_ and Deep_3_ features represent complementary, rather than redundant, pathology features for search.

##### Deep features improve search performance less than tissue and marker clinical covariates

Interested in the relative importance of deep features and clinical covariates, we then ask if Deep_3_ features or the tissue type covariate are more important for search. In the context of our 2,412 hand-engineered features, we find Deep3 features improve search performance less than the tissue type covariate (0.5720±0.0036 vs 0. 6533±0.0025, *U* = 0, *p* = 0.0001806). We additionally ask if Deep_3_ features or the marker mention covariate are more important for search. In the context of our 2,412 features and tissue type covariate, we find Deep_3_ features improve search performance less than the marker mention covariate 0.6602±0.0022 vs 0.6908±0.0021, *U* = 0, *p* = 0.0001817). We conclude that for our dataset’s size and diversity, search benefits most from carefully identifying and integrating simple clinical covariates for context, rather than focusing on advanced image analysis techniques such as deep learning.

#### S5.13.4 Deep features trained only on natural images outperform hand-engineered features for search, and offer best performance when combined with other features

##### Deep features trained only on natural images offer best measured performance, when combined with other features

To determine if deep convolutional neural networks trained only on natural images (e.g. cats and dogs) represent useful information for histopathology disease state search beyond what we have represented, we ask if ImageNet_2048_ features improve search performance beyond the best performance we could achieve without ImageNet_2048_ features. Therefore, we consider performance in the context of including 2412 hand-engineered features, tissue type and marker mention covariates, SIFT_k5_ features, and Deep_3_ features. We find ImageNet_2048_ features significantly improve search performance to a substantial degree (0.7006±0.0026 vs 0.7618±0.0018, *U* = 0, *p* = 0.0001817). We conclude for pathology search that there is complementary information represented in the ResNet-50 deep neural network trained only on natural images.

Deep features trained only on natural images outperform hand-engineered features, in the context of clinical covariates Both (a) the ImageNet_2048_ features from a deep convolutional neural network and (b) the 2,412 hand-engineered features from a variety of human-designed published algorithms are made for natural images, rather than histopathology images. The hand-engineered features are intrinsically interpretable, because a human defined each step of the algorithm’s behavior *a priori*. In contrast, deep convolutional features are the result of many layers of nonlinear transformations defined through training on data to minimize a loss function, so deep features are not interpretable as human-designed features are. To determine the pathology search performance penalty, if any, from using (a) less interpretable deep features of natural images rather than (b) more interpretable hand-engineered features of natural images, we compare search performance using (a) ImageNet_2048_ features to (b) the 2,412 hand-engineered features. In the context of tissue type and marker mention clinical covariates, we find ImageNet_2048_ features significantly improve search performance, again to a substantial degree, compared the the 2,412 hand-engineered features (0.6908±0.0021 vs 0.7517±0.0025, *U* = 0, *p* = 0.0001806). Excluding these covariates, to compare only ImageNet_2048_ features to the 2412 hand-engineered features alone, we again find ImageNet_2048_ performs significantly better (0.5640±0.0024 vs 0.6406±0.0046, *U* = 0, *p* = 0.0001817). In the context of tissue type and marker mention clinical covariates, we find (a) ImageNet_2048_ features also significantly improve search performance compared to (b) the 2,412 hand-engineered features combined with both SIFT_k5_ features and histopathology-trained Deep_3_ features (0.7517±0.0025 vs 0.7006±0.0026, *U* = 0, *p* = 0.0001817), which indicates ImageNet_2048_ features are the most important visual feature we measured. In the context of clinical covariates and ImageNet_2048_ features, search performance is significantly (albeit only slighty) improved when also considering the 2,412 hand-engineered features, SIFT_k5_ features, histopathology-trained Deep_3_ features (0.7517±0.0025 vs 0.7618±0.0018, *U* = 0, *p* = 0.0001806), demonstrating the synergy among these features. We conclude deep features are more effective than hand-engineered features for encoding histopathology images for search. This result may be confounded due to ImageNet_2048_ features encoding every image corner-to-corner in a grid fashion (which does not omit pixels, as grid cells have typically >50% overlap), while the 2,412 features are all based on a 512×512px center crop (which omits some pixels from the original image) (Fig 3C). However, we find best search performance when combining deep features, hand-engineered features, and SIFTk5 features.

##### Deep features trained only on natural images have intrinsically general properties that inform histopathology search, rather than learned nonlinear relationships of these features informing histopathology search

Given that natural-image-derived ImageNet_2048_ features provide a powerful representation for histopathology image search, we ask if this representational power comes from (a) general-purpose properties from the ImageNet_2048_ features themselves that hold even for pathology or (b) the Random Forest learning pathology-specific nonlinear relationships among the ImageNet_2048_ features for histopathology applications. To test this, we compare search performance of (a) the L1 norm of the ImageNet_2048_ features to (b) the Random Forest similarity trained on the ImageNet_2048_ features. We find the L1 approach marginally outperforms the Random Forest similarity approach, but this is not statistically significant, so any performance differences may be due to chance alone (0.6432±0.0000 vs 0.6406±0.0046, *U* = 70, *p* = 0.1153). This suggests the Random Forest does not learn nonlinear relationships among ImageNet_2048_ features that improve histopathology search performance. Rather, this suggests general properties of the ImageNet_2048_ features themselves are important for histopathology search. Moreover, we do find some ImageNet_2048_ features are more important than others for disease state prediction (Fig S10). However, we did not observe interpretable correspondences between ImageNet_2048_ feature activations and histopathology (Fig S11). We also do not observe that ImageNet_2048_ features form clusters of patients (Fig 6B). We conclude that although ImageNet_2048_ features empirically perform well, it may be desirable to use features that both empirically perform well and have general properties that “make sense” for histopathology.

### S5.14 Supplementary Experimental design and evaluation

We evaluate our classifiers using 10-fold cross validation, which is the default evaluation scheme in Weka [35]. Our data are saved in ARFF file format, so our findings can be reproduced in Weka without the need for writing software code. This approach allows software code we write to be compared against the unperturbed gold standard of Weka defaults. We follow Weka’s default of ten replicates of 10-fold cross validation, to estimate bounds of accuracy and Area Under Receiver Operating Characteristic (AUROC) performance metrics. This approach will give reproducible results wherever Java and Weka run, e.g. a laptop, a server, a supercomputer, or a cloud computer. This approach will work on all operating systems that support Java, e.g. Linux, Mac, and Windows.

### S5.15 Supplementary Computational hardware and software

We use Weka version 3.8.1 [35] on a ASUS Intel core i7-6700HQ 2.6GHz 4-CPU laptop with 16GB RAM for baseline analyses and Random Forests This laptop was also used for software development and automatically downloading Twitter data from participating pathologists. This laptop ran the Windows 10 operating system, which in turn ran the Oracle VirtualBox virtual machine manager, which in turn ran Debian Jessie 3.16.7-ckt20-1+deb8u3 and Linux kernel 3.16.0-4-amd64. Weka and our other pipeline components ran within Debian.

For deep learning, we use Keras version 2.1.4 and TensorFlow version 1.10.0 on an MSKCC supercomputer with several Nvidia Titan-X GPUs and dozens of Nvidia GTX 1080 GPUs running CUDA 8 and cuDNN 5.

### S5.16 Supplementary Comparison with prior studies

#### S5.16.1 Pathology-agnostic neural nets, SIFT nuclear features, and texture features

Many other groups perform pathology search with deep neural networks or hand-engineered features. Komura *et al* [18]. and Hegde *et al* [19]. use a deep neural network for pathology image search, taking a pathology-agnostic approach by not performing machine learning on histopathology images. Hegde *et al.* go further by comparing search performance of their neural network method, called SMILY, to a simpler baseline method using Scale-invariant feature transform (SIFT) [45]. Zhang *et al.* suggest SIFT interest points tend to cover cell nuclei [76]. However, Lowe, the author of SIFT, notes SIFT features do not represent color or texture, “features […] use only a monochrome intensity image [and] texture measures […] could be incorporated” [45]. For pathology search this may handicap SIFT compared to a neural network that can represent color or texture. Indeed, when we train our neural-network-random-forest hybrid classifier (Fig 3) on pathology images, we find features that represent texture (Local Binary Patterns Pyramid [59], Local Binary Patterns [58]) and color (Color Histogram [54], Color Correlogram [56], etc [55,57]) are the most important non-deep visual features (Fig 4). For decades, texture and color have been known to be important in pathology [46], and this motivates reproducible procedures for staining and slide preparation. Recently, Linder *et al* [47]. report Local Binary Patterns are important texture features to distinguish epithelium from stroma in colorectal cancer. Kather *et al* [48] go further, using Local Binary Patterns and other texture features to distinguish stroma, epithelium, immune cells, normal tissue, etc in colorectal cancer. Linder posit Local Binary Patterns are robust to changes in staining, illumination, and camera settings – useful properties for building a robust classifier from a globally distributed dataset like ours. Though we find using SIFT features alone for search performs better than chance, we find SIFT-based features alone perform worse that every other alternative feature representation we tested (Fig 10, Table S1). This may suggest that the visual signatures of disease state in pathology involve more than covering cell nuclei, and some of these signatures may be uncovered by a Random Forest [31] as nonlinear relationships of texture and color. For pathology search, we use Random Forest similarity derived from our interpretable classifier trained on pathology images, rather than require explicit similarity annotations from pathologists as training data [77]. We find pathology-specific covariates improve our classifier and search performance (Figs 9, 10B). A variety of search approaches for pathology search, also known as Content-based image retrieval (CBIR), have been reported [78,79], including SIFT [43], SIFT with neural networks [44], and dimensionality reduction [80].

#### S5.16.2 Pathology-specific neural networks

Otálora *et al* [20]. take a transfer learning [12–15] approach to pathology search by adding an auxilary layer on top of a frozen pretrained deep neural network. They train this auxilary layer to predict if a prostate image shows Gleason [81] grade of 4 or more, or not. They do not report classification performance. The auxiliary layer feature vector is used for prostate image search, and the search is shown to perform better than a simple baseline, i.e. Color and Edge Directivity Descriptor (CEDD) [55]. We also use CEDD in our work and find it important (Fig 4). However, our classifier training is pan-tissue and pan-disease, neither specifically prostate nor specifically high Gleason. Like Otálora, we search PubMed (Fig 1). However, to filter PubMed for histopathology images, Otálora use a light microscopy detection algorithm [82] while we use a Random Forest trained for H&E detection with leave-one-pathologist-out AUROC of 0.95 (Fig 7). In prior work, we trained a deep neural network end-to-end to predict SPOP mutation in prostate and repurposed the classifier for search [16]. Peng *et al* [83]. jointly train a deep neural network for prediction and search, using colorectal histopathology images labeled with nine possible classes, i.e. adipose tissue, background, cellular debris, lymphocytes, extra-cellular mucus, smooth muscle, normal colon mucosa, cancer-associated stroma, and colorectal cancer epithelium. However Peng do not report search performance as mean average precision or precision@k for k=1,2,3,…,10 (Fig 10B), so Peng’s results may be difficult for others to interpret. Instead, Peng report how many images had “perfect retrieval precision of 10 true neighbours”, and find their method performs 30% higher than their baseline. Peng cite Cao [84] and Cao [85] for reporting this way, but both report mean average precision, so Peng search performance between data sources are not clear to us.

#### S5.16.3 Computational pathology studies not pan-tissue pan-disease

For dermatopathology, Esteva *et al* [40]. predict if an image shows disease that is benign, malignant, or either benign/malignant – but do not consider nontumoral disease such as infections. For a few tissues including prostate, Campanella *et al* [42]. predict if an image shows cancer or not, but similarly do not differentiate non-neoplastic from benign disease, which may be an important clinical consideration [22]. Neither perform search. From optimal coherence tomography (OCT) imaging of eye pathology, De Fauw *et al* [41] predict referral urgency as urgent, semi-urgent, routine, or observation. This is not a prediction of disease state. De Fauw do not mention cancer, but instead focus on diseases we consider to be nontumor, such as diabetes and macular degeneration. We are not aware of any pan-tissue pan-disease datasets other than ours. We believe our pan-disease method serves patients with diseases of poverty (e.g. many forms of infection, 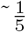 of diseases in our data) as well as patients with diseases of affluence (e.g. many forms of cancer, 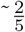 of diseases in our data).

#### S5.16.4 Deep and shallow learning not on same task

In the field of cardiology, deep learning has been trained on a separate task (i.e. epicardial adipose tissue volume), then used by shallow learning (i.e. XGBoost) to predict myocardial infarction [86]. Our approach differs in that we train both deep and shallow learning (i.e. a Random Forest) on the same task, namely disease state prediction (Fig 3B,C). In principle, our approach allows deep learning to, in a data-driven manner, derive features that are important for disease state prediction, which may complement the hand-engineered features and clinical covariates we use for shallow learning. Indeed, we find there are such important and complementary deep features for shallow learning of disease state prediction (Fig 4) and search (Fig 10).

### S5.17 Supplementary Caveats

#### S5.17.1 Patient case across multiple tweets

A feature of these data is that a particular patient might be represented in more than one image. Any given patient might be discussed in multiple tweets, each of which contain one or more images. A patient case may be spread across multiple Twitter threads as well, e.g. the first tweets asks pathologists for their opinions as a quiz, while a second tweets in a new thread provides the correct diagnosis. We believe leave-one-pathologist-out cross validation is a reliable way to prevent the same patient’s data from being in both a training and a test set for machine learning evaluation. However, in the future, we may find a patient is shared by multiple pathologists, e.g. when a pathologist donates his/her slides to another pathologist. Currently, we have not observed the same patient case being shared by multiple pathologists.

#### S5.17.2 Near-duplicate detection

There is room for improvement in automated duplicate detection methods. A pathologist may first tweets an image that has no hand-drawn marks, but later reply with an image that includes hand-drawn marks such as circles and arrows to indicate a region of interest. Pathologists may also re-share the cases of other pathologists, with minor modifications, such as white balancing. In future work, these near-duplicates should be automatically detected. Duplicates may artificially inflate performance metrics.

#### S5.17.3 Disease state annotations vary in text

Our dataset is only as good as the accuracy of the hashtags and diagnoses made by the collaborating pathologists and pathologists who comment on the cases. The more pathologists that contribute to the database, the higher the risk for errors and inconsistencies. Indeed we note some uses of the #bstpath hashtag to describe breast pathology (Section S5.6.1). We should remember the fun and voluntary nature of sharing cases on social media.

While our disease state text processing algorithms take a consensus vote among the pathologists discussing the patient case, these methods are not perfect, and our manual annotations to correct this may be incomplete. We hope that by sharing our data with the community, more corrections may be made, improving the quality of our dataset.

#### S5.17.4 Disease state evidence varies in images

For our Random Forest baselines, we crop images to convert rectangular images to be uniformly square for the machine learning. However, pathologists may include diagnostic information only at the extreme edges of an image that are cropped out. A case of this from B.X. involves a hydatid cyst in the extreme right of an image, which would be cropped out^8^. This hydatid cyst indicates *Echinococcus* infection, so the case is nontumoral. Our set-based deep learning approach is an incomplete remedy to this problem, where we train on 224×224px image patches sampled throughout the original image, then test using 21 224×224px patches systematically sampled throughout the original image. Although this approach samples the entire image, the remedy is incomplete in that the ground truth is not uniform throughout the image. For example, it is only based on one corner of the image that there is histological evidence of nontumor disease. It may be helpful here to use more advanced methods, which make fewer assumptions about the ground truth and allow weaker supervision, but such methods may come at a cost of requiring more data than we have currently.

#### S5.17.5 Algorithm and labeling inaccuracies

We do not expect that our text classification algorithm (Fig S8) perfectly interprets disease state from the text associated with an image. Moreover, we also do not expect our manual annotation process to be perfect, e.g. some stains may be incorrectly labeled as H&E, IHC, etc. We manually curate 10,000+ images, so human error as low as 1% means a handful of images are incorrectly labeled, but there could be more. By sharing our data with more pathologists and data scientists, we intend to gather feedback and correct any inaccuracies here, then measure disease state classification performance changes.

#### S5.17.6 No automated quality control

Finally, the size of the dataset is both a blessing and a curse. A large and diverse dataset is required to provide the most benefit to computational pathology. However, quality control for such large datasets is most feasible if done automatically, and automated quality control cannot deal with all issues. For example, some pathology images include marks designating a particular pathologist as the contributor of that image. Other pathology images have been marked by pathologists with arrows and circles. Our automated quality control pipeline enables us to rapidly discriminate pathology from non-pathology images, but is not able to address these other challenges. Future steps will need to be taken for more specialized quality control.

### S5.18 Supplementary Future directions

#### S5.18.1 Acquiring more data

The first step is to expand the size of this dataset by recruiting more pathologists via social media. With more data, we hope to improve performance on discriminations that were the most difficult (e.g. those involving gynecological pathology). More data may facilitate machine learning methods that discriminate between similar but less frequently used stains, such as H&E vs Diff-quik, rather than H&E vs IHC. More data might also enable us to distinguish particular organs or tissues within a histopathology tissue type, e.g. distinguish kidney tissue from bladder tissue. With increased sample size and increased tissue of origin granularity, it may be possible to predict metastatic tissue of origin. Finally, a larger dataset might also include more rare cases that can be useful for machine learning techniques that can support diagnoses.

#### S5.18.2 Expanded and specific hashtags

A second step is advocacy on social media, for (i) sharing normal tissue data, and (ii) expanded pathology hashtags. Normal tissue complements our existing “relatively unimportant” artifact and foreign body data, such as colloids and gauze, which are typically not prognostic of disease. Normal tissue also complements the description of tissue morphology in our data, if we tend to have only cancerous or diseased tissue. Separately, more descriptive hashtags may reduce our manual annotation burden, and obviate the need for us to ask pathologists to clarify what stain was used or what the tissue is. Moreover, molecular hashtags may complement the histology we see. However, we understand that for pathologists sharing cases on social media is probably a fun and voluntary activity, rather than a rigorous academic endeavor, so it may not be appropriate for us to suggest pathologists use terms from synoptic reporting in hashtag format in their Tweets. Moreover, the size of tweets is limited to 280 characters, so more than 3-4 hashtags per tweets is probably infeasible. Some pathologists are already close to this limit without using additional hashtags.

We encourage the adoption of hashtags that explicitly identify what stains or techniques are used (this is not an exhaustive list):

1. #he indicates there are one or more H&E-stained images in the tweets.
2. #ihc indicates there are one or more IHC-stained images in the tweets.
3. #pas indicates there are one or more periodic acid-Schiff images in the tweets.
4. #diffq indicates there are one or more diff-quik images in the tweets. There is a common misspelling of diff-quick, so our hashtag avoids this misspelling.
5. #gross indicates one or more gross section images are in the tweets. This is typically fresh cut tissue, e.g. an entire excised tumor or a large piece of an organ.
6. #macro indicates an unmagnified picture of a microscopy slide. Unfortunately, such pictures are occasionally referred to as gross.
7. #endo indicates one or more endoscopy images are in the tweets.
8. #ct indicates one or more CT scan images are in the tweets.
9. #xray indicates one or more X-ray images are in the tweets.

We encourage hashtags to describe not only the histological features of a case, but also the molecular features of a case. Again, this hashtag list is far from exhaustive.

1. #braf indicates the BRAF gene is known to be mutated, perhaps through sequencing.
2. #msi indicates micro-satellite instability, which again may be evident from sequencing.
3. #desmin indicates that the IHC used targets desmin.

We encourage the adoption of hashtags that explicitly identify any artifacts, art, or pathologist annotations/marks on the image.

1. #artifact or #artefact indicates there are artifacts or foreign bodies in one or more images, such as colloids, barium, sutures, Spongostan™, gauze, etc. We encourage the tweets message text to identify what the artifact or foreign body is.
2. #pathart is a hashtag in use today, but unfortunately it is used in two ways: (i) to identify naturally-occurring and unmodified pathology images that are “pretty” or “interesting” as natural works of art, and (ii) to identify images that have been modified by the pathologist herself/himself to be “funny” or “interesting”. The trouble is (i) is “acceptable” pathology for analysis while (ii) is not. We advocate for the continued use of the #pathart hashtag, but with clarification, below:
3. #drawn or #annotated indicates the pathologist made hand-drawn marks on one or more images, such as arrows, circles, or artistic manipulations. Artistic manipulations may include drawing exclamation points, question marks, eyes, mouths, faces, skulls, cartoon bodies, etc on the image. So, “#pathart #drawn” is likely a pathology image with artistic drawn marks that prevents the image from being an “acceptable” pathology image for analysis, while “#pathart” without “#drawn” is likely a pathology image that is a naturally occurring unmodified histology image that is an “acceptable” pathology image for analysis.
4. Alternatively, “#pathfun” or “#pathdrawing” may refer artistically manipulated pathology pictures, leaving “#pathart” exclusively for naturally-occurring pathology that are “pretty” or “interesting” from an artistic perspective.

We encourage the adoption of hashtags that give other information about the image.

1. #pathbug is an existing hashtag that indicates a parasite or other co-occurring non-human organism is depicted in one or more images in the tweets. The #parasite tag is sometimes used instead.
2. #panel indicates one or more multi-panel images are in the tweets.

We encourage all adopted pathology-related hashtags to be registered in an ontology, e.g. https://www.symplur.com/healthcare-hashtags/ontology/pathology/. A hashtag ontology can standardize the hashtags used, which in turn can (i) help pathologists in the same subspecialty find each other, and (ii) simplify computational analyses of hashtag text.

## Footnotes

1 ImageIO documentation available here: https://docs.oracle.com/javase/7/docs/api/javax/imageio/Image10.html

2 Courts in the United States have ruled that images posted to social media are still owned by their authors and are not public domain. Indeed, in Morel v. AFP, AFP was ordered to pay Morel $1.2 million for copyright infringement because AFP used images that Morel posted to social media.

3 Normal cerebellum case by S.Y. at https://twitter.com/Sty_md/status/821840894634565632

4 A case of this is from author *K.H.*, where a different pathologist gave the diagnosis, and he agreed. We summarized this as “metastatic lobular carcinoma” in the auxiliary annotation file for the tweet https://twitter.com/Ho_Khanh_MD/status/999989201734197250.

5 A case of this is from author *M.P.P.,* where M.P.P. wrote “IDC DIN LISN” directly on a shared histology image in the tweet https://twitter.com/dr_MPrieto/status/890118713155997696 so we wrote this text in the auxiliary annotation file for the tweet.

6 A case of this is from K.H., observing iron pill lesions in stomach biopsy https://twitter.com/Ho_Khanh_MD/status/963800933716123648.

7 For this formula please see https://github.com/keras-team/keras/issues/6444

8 Case at https://twitter.com/BinXu16/status/980404471833313280 “Kudo to @drkennethtang @luishcruzc and @DrGeeONE The answer of this case can be seen in the right corner of the 3rd picture. Dx: Echinococcus (hydatid cyst) with necrotizing pneumonia, abscess, and granulomatous inflammation. Additional high power pictures attached.”

